# Disruptors of sestrin-MAPK interactions rejuvenate T cells and expand TCR specificity

**DOI:** 10.1101/2024.05.17.594698

**Authors:** Alessio Lanna, Clara D’Ambra, Federica Rinaldi, Luisa Chocarro, Manuel Delpero, Melania Capitani, Michael Karin

## Abstract

Whereas treatments that reactivate exhausted T cells are available, strategies to rejuvenate terminally differentiated senescent lymphocytes are yet to be developed. Senescent T cells, with short telomeres and inactive telomerase, are different from exhausted cells, and may form due to defective telomere transfer reactions upon contact with antigen presenting cells (APCs). Senescent T cells are characterized by presence of sestrin-MAPK kinase activation complexes (sMACs), large immune-inhibitory protein assemblies of sestrins bound to a stress/energy sensing kinase (AMPK) and three functional effector kinases (ERK, JNK and p38 MAPKs). Here we described first in class Disruptors of the Sestrin-MAPK immune-inhibitory Complex (DOS), which target sMAC to ubiquitin-dependent proteasomal degradation, resulting in long-term sestrin transcriptional inhibition, increased T cell fitness, and generation of long-lived stem like memory features. Strikingly, the DOS generated stem T cells present *de novo* antigen-specific T-cell receptor DNA rearrangements that precede their future expansion. Although largely senescent at the point of treatment, the DOS regenerated T cells, with stem features and new TCRs, initiated immune-protective rejuvenation-dependent responses to new challenges, with or without vaccination. Therefore, it is possible to generate new T cell clones from formerly senescent cells and expand immune specificity.

**Highlights:**

- DOS are the first sestrin-MAPK binding disruptors
- DOS rejuvenate T cells (DOS-juvenation)
- DOS-juvenated T cells protect old mice from lethal infections, with or without vaccination
- DOS-juvenated T cells exist as stem-like cells that undergo antigen-specific TCR revisions

## Introduction

The aged human population is expected to reach over two billion by 2050^1^, with increased risks of age associated disorders due to deterioration of immunity^2^, and elevated health and social costs. Previous attempts to confer protective immunity to aged or vulnerable populations yielded modest results. Adjuvanted or high antigen influenza vaccines were developed, but the results are still suboptimal^3^; showing partial and heterogenous vaccine benefits, which barely exceed the prevention of influenza-related diseases compared to standard trivalent influenza vaccines^4^. Other approaches relying on pre-administration of metabolic tuning drugs prior to immunization were plagued by low compliance and failure to protect from respiratory infections^5^ or paradoxical activation of T cell senescence^6^. Furthermore, some of the pathways that were clinically investigated (e.g., mTOR) are completely absent from highly differentiated or senescent T cells^7^. Other strategies that eliminate senescent cells (senolytics)^8^ may have off-target effects resulting in destruction of the aged immune system. In addition, senoblockers^9^ would be mostly ineffective in T cells that are already senescent, whereas senomorphics^10^ can reverse specific aspects of the senescence associated secretory phenotype (SASP)^11^ without achieving broad rejuvenation. Finally, checkpoint inhibitor therapy targeting exhausted T cells during ageing is often short-lived^12–14^. As such, senescence and exhaustion are two distinct dysfunctional states of the T cell, and no long-term strategies that directly rejuvenate senescent T cells exist.

In humans, hypo-proliferative senescent T lymphocyte populations that lack CD28 and CD27 costimulatory receptors accumulate with age^15^. These cells are characterized by immune-inhibitory sMAC complexes^6,7^, formed when the sestrin stress proteins (sestrin1, 2 and 3) bind to, and activate, AMPK, that triggers scaffold assisted MAPK signals responsible for age-related loss of immunity^7^. Thus, AMPK, which is activated in response to increased catabolic demands^16^, and promote T cell survival^17,18^ likely through direct GATOR/mTOR inhibitory binding^19^, has opposite pro-senescent functions in the sMAC. Correspondingly, sestrins can be found in both sMAC or GATOR/mTOR complexes, but while regulation of GATOR/mTOR requires inhibitory interaction with the C-terminal domain of sestrin^20,21^, sMAC activation may require direct binding to the N terminal portion of the sestrins, which is involved in the oxidative stress response characterising senescent cells^21,22,23^.

Through simultaneous MAPK auto-phosphorylation reactions in fact, sMAC drives T cell senescence. Upon activation, each sMAC-bound MAPK operates at the same time, in the same T cell, being responsible for unique hallmarks of its age-related dysfunction^6,7^. Thus, albeit some compensatory, NK-like, innate T cell states have been ascribed to sMAC^24^, adaptive T cell immunity is suppressed by sMAC activation during ageing.

Here we identified a hitherto unknown pharmaceutical class that allows long-term immune rejuvenation through sMAC disruption and generation of de novo T cell receptor rearrangements in formerly senescent cells.

## Results

### Discovery of DOS

We designed simple *in vitro* assays using minimal sMACs composed of recombinant sestrin and AMPK incubated with 156 pentameric sestrin peptides previously proposed to mediate an interaction between the two proteins; such peptides spanning the N-term domain of sestrins not involved in GATOR/mTOR regulation (Figures S1A and S1B)^21,25^.

In these assays, ATP consumption by AMPK allows its activity assessment against a target substrate (SAMsite) via released luminescence, such that any inhibitory peptide would oppose sestrin action, resulting in an increased ATP content in solution, reduced luminescence, and disruption of AMPK activation in the sMAC (Figure S1C).

Upon peptide cyclisation, we identified the disruptor of sMAC (DOS) 46L, the lead compound (Figures 1A and S1D). To investigate possible DOS binding sites, we used Autodocking and AlphaFold algorithms^26^ that allow to predict protein structures via automated deep learning. DOS46L was predicted to bind to sestrins in a site that is distal to that required for GATOR/mTOR regulation^27^ (Figures 1B-Site 1- and S2A). To confirm this, we performed site directed sestrin 2 mutagenesis in primary T cells. We used phospho-flow and found that DOS failed to inhibit AMPK in mutant T cells bearing two punctiform sestrin aminoacid variations into alanine (436T>A and 306P>A), demonstrating these sites were required for DOS-driven sMAC suppression via sestrins (Figures S2B and S2C). By contrast, alanine mutation of less energetically favored sestrin-DOS interaction sites (docking predictions) had no effect.

**Figure 1.**
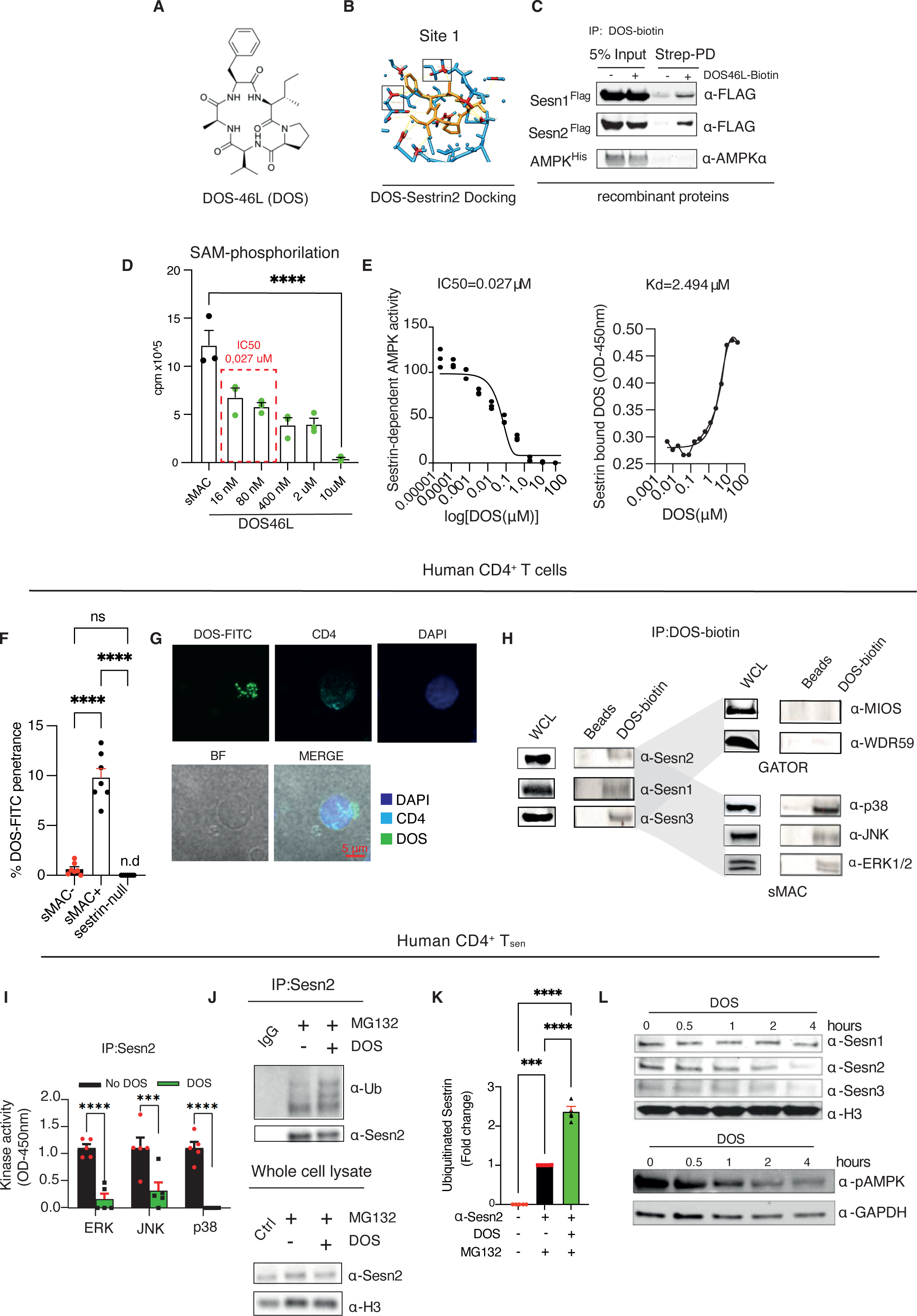
Discovery of DOS. (A) Molecular structure of the lead compound DOS46L (hereafter, DOS). (B) DOS binding site to Sestrin 2 predicted by AutoDock Vina and AlphaFold algorithms. (C) Co-Immunoprecipitation of biotinylated DOS46L (DOS; used at 10μM, throughout) followed by recombinant tagged protein detection, as indicated. (D) Recombinant sMAC activity investigated by liquid scintillation. Cpm, counts per minutes. (E) Half maximal inhibitory concentration (IC50) of DOS against recombinant sestrin-AMPK complexes by *in vitro* kinase assays with SAMS peptides and ADP-glo kinase assessment (left); equilibrium dissociation constant (Kd) of sestrin bound biotinylated DOS assessed by ELISA with streptavidin antibodies (right). (F) Quantification of selective DOS penetration in sMAC negative and sMAC positive primary human CD4^+^ T cells by flow cytometry, 30 minutes after DOS treatment (*n* = 7 donors). (G) Individual channels of representative confocal imaging (*n* = 3 donors) of DOS (green) uptake in human CD4+ T cells. Scale bar, 5 µm. (H) Co-Immunoprecipitation (IP, hereafter) of biotinylated DOS from human CD4^+^ T cell extracts after overnight treatment. Presence or absence of sestrin, p38, JNK, ERK (sMAC) or MIOS and WDR59 (GATOR) was confirmed by western blot in the IP. Whole cell lysates (WCL) controls are shown (same extracts; run in parallel). (I) ERK, JNK and p38 MAPK kinase activity (autophosphorylation) among T_sen_ sestrin 2 immunoprecipitates (sMACs) in the presence or in the absence of DOS for 30 minutes (*n* = 5 donors). Kinase activity was detected by ELISA assays. (J) Human T_sen_ were pre-treated with the proteasomal inhibitor MG132 (1μM) for 30 minutes followed by addition of DOS for 2 hours. Controls were not treated with DOS. Total lysates and sestrin 2 immunoprecipitates were then analysed by immunoblotting of ubiquitin and sestrin 2. H3, whole cell lysate control. (K) Pooled ubiquitination data (*n* = 5 donors) for experiments as in (I). Data are shown as fold change to the untreated control (NO DOS, in the presence of MG132), set as 1. (L) Time dependent assessment of sestrin expression (1, 2 and 3) by DOS in human Tsen (top) and AMPK phosphorylation in the same cells assessed by western blotting (bottom). Representative of 6 donors. In (E-F, I and K) one-way Anova with Bonferroni post-correction for multiple comparisons was performed. **P<0,01; ***P<0,001; ****P<0,0001. Error bars indicate SEM.

We next used recombinant proteins and biotinylated DOS, and confirmed direct DOS binding to sestrins (but not AMPK) via direct biotinylated DOS immunoprecipitation with streptavidin beads and immunoblot of the precipitated proteins with anti-flag antibodies (Figure 1C). Any impact on AMPK activity within the sMAC must then be mediated by direct DOS interaction with sestrin. Consequently, using radioactive P32 measurements (Figure 1D) or AMPK luminescence assays and recombinant sestrins (as per initial DOS screening), we found that sestrin-driven AMPK activation was robustly inhibited by nanomolar concentrations of DOS, and these AMPK inhibitory DOS concentrations were 20-30 times lower than those at which half of sestrins were bound by DOS in parallel enzyme-linked immunosorbent assays (ELISA)-based binding assays (IC50 *vs* Kd; Figure 1E). Thus, DOS triggers AMPK inhibition in the sMAC, even when only a fraction of sestrin is bound by the drug.

To study DOS entrance in primary T cells, we exposed purified (unfractionated) human CD4^+^ T cells to fluorescently conjugated DOS (FITC-DOS), followed by anti-CD3/CD28 activation, and performed flow cytometry assessment. We found that DOS selectively penetrated in the sestrin2^+^ p-p38^+^ (sMAC^+^) T cells within minutes of treatment (Figure 1F), especially in immune senescent CD27^-^ CD28^-^ CD4^+^ T cells (hereafter, T_sen_) that possess the highest levels of sMACs^7^ (Figure S3A). Some background DOS penetrance could also be detected in ∼ 1% sestrin2^-^ p-p38^-^ (sMAC^-^) T cells, likely due to DOS entrance in T cells where sMACs are not yet fully formed (for instance, beginning expressing only sestrin 1 or 3^7^). Correspondingly, DOS did not enter T cells that had been depleted of all three sestrins by shRNAs (sestrin-null T cells, Figure 1F). Likewise, early-stage (non-senescent) CD27^+^ CD28^+^ CD4^+^ T cells (hereafter, T_erl_), which possess very low sMAC levels^7^, were largely impermeable to the drug, perhaps due to differences in metabolism and membrane composition between senescent versus non-senescent cells (Figure S3A). Any effect of DOS on T cells must derive therefore from its effect on senescent or other sMAC containing lymphocyte populations that are destined to senescence^7^. Imaging studies confirmed intracellular DOS penetrance in activated CD4^+^ T cells (Figure 1G). However, DOS did not alter the extent of CD4 internalization upon TCR activation (Figure S3B). Finally, confocal imaging with antibodies to sestrin and the endoplasmatic reticulum (ER) marker KDEL further confirmed DOS penetrance, and showed its colocalization to the ER where the sMAC resides (Figure S3C).

To confirm DOS binding to primary T cell sMACs, we exposed human CD4^+^ T cells to biotinylated DOS, and immunoprecipitated DOS-bound sMACs with the streptavidin beads. This confirmed that DOS directly bound to the immunoprecipitated primary T cell sMAC composed of sestrin and MAPK kinases but not to GATOR/mTOR complex proteins (MIOS and WDR59; DOS-biotin, Figure 1H). Parallel assessment confirmed sestrin and MAPK and GATOR protein presence in the human CD4+T cell lysates subjected to the IP (whole cell lysate; Figure 1 H). Correspondingly, human T_sen_ had very low expression of GATOR/mTOR complex proteins as assessed by immunoblotting (Figure S3D).

We studied the effect of DOS in T_sen_ where the sMAC is abundant^7^, and immunoprecipitated the complex with anti-sestrin 2 antibodies followed by ELISA-based *in vitro* kinase assessment of the sestrin-bound MAPkinases, as described^7^ (Figure 1I). Because each MAPK in the sMAC undergoes auto-phosphorylation, we then measured the self-phosphorylation reactions^7^ denoted by the addition of exogenous ATP in the IPs, an established readout to assess sMAC function. We found that each MAPK kinase activity was suppressed by DOS treatment (Figure 1I), demonstrating that DOS inhibited the immunoprecipitated sMACs. We then derived sMACs from AMPK proficient or deficient T_sen_ cells (obtained by lentiviral shRNA delivery to AMPK) and did IP *in vitro* kinase assays to compare their response to DOS, as above. AMPK deficient T cells were resistant to DOS action and phenocopied the effect of the drug (Figure S3E). Similar results could be also obtained when AMPK activity was pre-inhibited by compound C^28^ (Figure S3F). Taken together, DOS binds to sestrins, but require AMPK function for sMAC inhibition.

Next, we assessed the expression of sestrin after DOS treatment in human T_sen_ and compared it with donor-matched T_sen_ not exposed to the DOS. Within 2 hours of DOS-driven sMAC inhibition, all three sestrins (sestrin 1, 2 and 3) dissociated from AMPK as assessed by immunoprecipitation with anti-AMPKalpha antibodies coupled to ELISA based sestrin detection in the IP reactions (Figure S3G); and underwent ubiquitin-dependent proteasomal degradation among sestrin precipitates from the same cells probed with anti-ubiquitin antibodies (anti-ubiquitin *vs* sestrin signal in the IP sample; Figures 1J and 1K).

DOS T cell pre-exposure to the proteasome inhibitor MG-132 preserved both sestrin and AMPK expression, demonstrating proteolytic degradation of key sMAC components in response to DOS treatment (Figure S3H). Correspondingly, within 4 hours of DOS treatment, immunoblotting revealed notable sestrin downregulation, and AMPK inactivation in the T_sen_ (Figure 1L). AMPK inhibition was also confirmed assessing the phosphorylation status of Acetyl-COA carboxylase (ACC), its main downstream target, by both phospho-flow and immunoblotting (p-Ser79 ACC down-regulation, Figure S3I). Specific ACC phosho-flow assessment was further confirmed by Compound C treatment, phenocopying DOS action. The sMAC was also inhibited by DOS in parallel flow cytometry experiments with p-p38 and sestrin 2 antibodies among senescent human CD27^-^ CD28^-^ CD8^+^ T cells (Figure S3J).

To further study sMAC disruption, we used recombinant proteins. AMPK is activated when ATP is unloaded from adenine nucleotide-binding sites in its γ subunit (at least one out of 4 cystathionine β-synthetase (CBS) sites, most likely CBS3)^29,30^, to accommodate AMP at the same sites. Recombinant sestrins directly bound to his-AMPK-γ as assessed by immunoprecipitation with anti-histidine tagging (Figure 2A). When we assessed ATP presence among the his-AMPK-γ precipitates by ELISA measurements, we found that sestrin strongly promoted ATP unloading in the IP reactions (Figure 2B), consistent with sestrin activating AMPK via its γ-subunit^16^. When we added DOS to these assays, the drug opposed sestrin action on AMPK-γ almost instantaneously, enforcing ATP levels in response to sestrin treatment (Figure 2C). As such, DOS inhibitory activity against the recombinant sMAC could be counteracted by an allosteric AMPK activator (A-769662), that operates better when AMPK is ATP bound rather than in ATP free conditions^31^ (Figure 2D). Similar results were obtained when we used MANT-ATP (2’/3’-O-(N-methylanthraniloyl) adenosine 5’-triphosphate)^29^, a fluorescent analogue of ATP which competes with ATP and specifically binds to the CBS1/3 adenosine exchanging sites, indistinguishably with full-length AMPK, or simply the AMPK regulatory γ sub-unit. Accordingly, we found that DOS opposed sestrin-driven ATP unloading evident as restored MANT fluorescence in these assays (Figure 2E).

**Figure 2.**
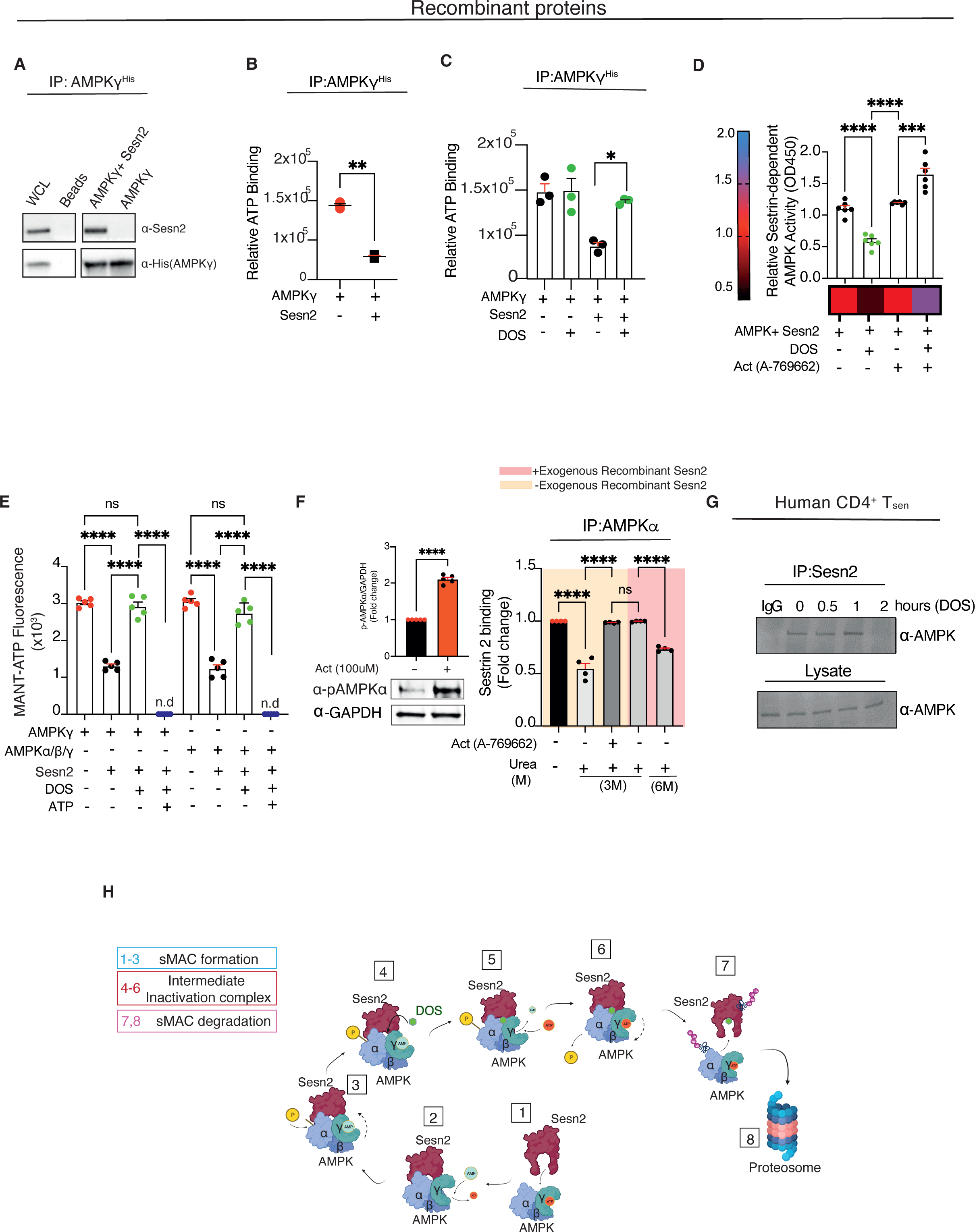
DOS-driven AMPK inhibition initiates sMAC dismantling. (A) Sestrin 2 and AMPKγ recombinant proteins were incubated at 30 °C. Direct binding was tested by immunoprecipitation of His-tagged AMPKγ using anti-His tag antibody followed by western blotting. (B) Investigation of sestrin-driven AMPK activation, assessed via direct ATP loading detection. Recombinant sestrin 2 and AMPK-γ were incubated as in (A). ATP was added to each experimental condition and detected after IP washing using the ATP Bioluminescence Assay Kit (*n* = 3 experiments). (C) AMPK-γ ATP binding in the presence or in the absence of DOS for 5 minutes. The reaction occurred as in (B). In control reactions, AMPKγ was not incubated with recombinant sestrin 2 but exposed or not to DOS (*n* = 3 experiments). (D) Recombinant sMAC activity assessed by ADP-Glo kinase Assay with or without the allosteric AMPK activator A-769662 (100 µM), in the presence or in the absence of DOS. Note that AMPK allosteric activation is paradoxically increased by co-treatment with DOS, reinforcing the notion^24^ that DOS treatment increases ATP loading on AMPK-γ (*n* = 6 experiments). (E) Fluorescent ATP-binding competition assay. MANT-ATP (10 µM) only fluoresces (448 nm) when bound and was reacted with full-length AMPK (5 µM) or the isolated AMPK-γ subunit (5 µM) in the presence or in the absence of sestrin, in a 1:4 ratio. Note that addition of DOS abolished sestrin-driven ATP unloading, evident as restored MANT-fluorescence in the competition reaction. ATP competition controls for the DOS reaction are shown (*n* = 5 experiments). (F) Effect of AMPK activity on its binding strength and affinity to sestrin. Non senescent T_erl_ were cultured for 3 days with or without allosteric AMPK activation by A-769662 followed by AMPK-α immunoprecipitation and AMPK and sestrin detection. The immunoprecipitates were exposed or not to a denaturating agent (Urea) or further exogenous sestrin 2 (50 ng), to evaluate sestrin binding strength, as indicated. Pharmacological activation of AMPK confirmed by immunoblotting (left), and binding disruption data expressed as fold to untreated controls, set as 1 (*n* = 4 donors, right). Data are expressed as fold change to the untreated control, set as 1. (G) Time-course of sestrin-AMPK dissociation upon DOS treatment of T cell sMACs obtained by sestrin 2 immunoprecipitation. Tsen cells were treated with DOS as indicated followed by sestrin 2 immunoprecipitation and AMPK-α immunoblotting. IgG, isotype control. (H) Schematic representation of DOS mechanism of action. DOS impedes sestrin-driven ATP unloading on AMPKγ leading to AMPK inactivation, loss of sestrin binding affinity, complex dissociation culminating with proteosomal degradation. In (B and F, left), a two-tailed paired t-test was used. In (C-F, right) one-way Anova with Bonferroni post-correction for multiple comparisons. *P<0,05; **P<0,01; ****P<0,0001. Error bars indicate SEM.

Using primary T cells, we enforced, or not, AMPK activation with the agonist A-769662 for 3 days in non-senescent T_erl_ that have very low spontaneous AMPK activation^6^ (Figure 2F, left), followed by AMPK immunoprecipitation with total anti-AMPK-α antibodies. We then incubated the IP reactions with increasing amounts of Urea, a denaturing agent that destroys protein binding. In parallel, recombinant sestrins were used in the IP reactions of agonist free T cells, to activate AMPK independently of A-769662 as a further activation control. Urea quickly disrupted sestrin-AMPK interactions in the IP from T_erl_ cells where AMPK was inactive, as detected by ELISA-based sestrin binding assessment. By contrast, sestrin-AMPK interactions were resistant to Urea in agonist activated T cells or in those agonist free T cells that had been exposed to the recombinant sestrin directly in the IP reaction (Figure 2F, right). These data show that activated AMPK has more sestrin binding affinity than inactive AMPK. Thus, once inhibited by DOS at the point of its adenosine exchanging sites, inactivated AMPK loses sestrin affinity, dissociates from sestrins (Figure 2G), and the individual sMAC components undergo proteasomal degradation (Proposed model; Figure 2H).

### DOS rejuvenates T cells

Next, we studied the effect of DOS in primary cultures. We used inverted light microscopy and confocal imaging and found reversal of expression of established senescence markers^32^ such as β-galactosidase (Figure 3A) and DNA damage associated foci (Figure S4A) in human T_sen_, cultured with ligation of the TCR (anti-CD3), recombinant human IL-2 and exposed to DOS for 7 days. Next, we assessed the effect of DOS on senescent T cell proliferation using sestrin deficient T cells (obtained by triple shRNA delivery to sestrins), which phenocopy DOS activation (thereby confirming DOS specificity of action), as well non-senescent human T_erl_ as a rejuvenation control. We found more cells in DOS treated T_sen_ cultures compared to T_sen_ not exposed to the DOS (cumulative population doublings; (PDs); Figure 3B left); these effects could not be simply explained by improved T cell survival because the cells had duplicated in number over the initial seedings, and to levels observed in DOS-free human T_erl_ (2-week cultures). These data show that DOS restores T_sen_ proliferation in culture. Furthermore, as expected, the proliferation effects were sestrin dependent because in parallel cultures with sestrin deficient T_sen_ cells, DOS-driven T cell proliferation was absent (Figure 3B right). Similar DOS-driven proliferative outcome (and sestrin reliance) was observed in sestrin proficient vs deficient T_sen_ analyzed by a cell dilution assay (Cell Trace Violet (CTV)) or bromodeoxyuridine (BrdU) incorporation into nascent DNA (Figures S4B and S4C). Moreover, analogue results were obtained by lentiviral depletion of AMPK in long term T_sen_ cultures (1-month; Figure S4D), where the effects of DOS were phenocopied (and thereby nullified) by genetic inhibition of AMPK. Hence, DOS action on T cells requires pre-existent sMAC activation.

**Fig. 3.**
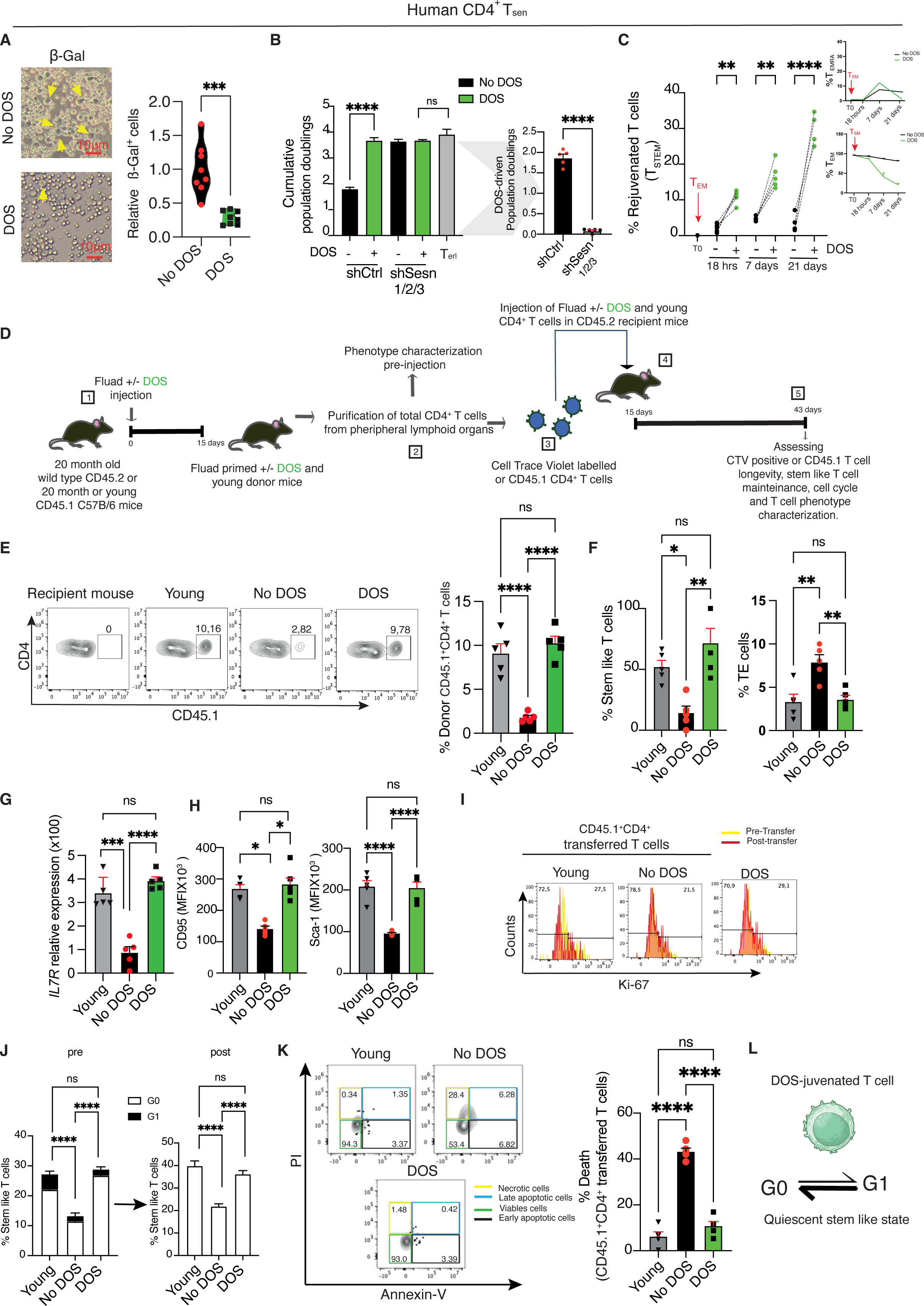
DOS-juvenation of T cells. (A) Senescence-associated β-galactosidase expression in DOS-treated (DOS-juvenated) or untreated T_sen_. Cells were purified and cultured for one week in the presence of anti-CD3 (0.5 μg/mL) and rh-IL-2 (5 ng/mL), then stained to detect β-galactosidase activity. Representative image on the inverted phase-contrast microscope (left) and relative quantification (right, *n* = 8 donors). (B) Population doublings of human T_sen_ (transduced with irrelevant scramble) and sestrin null CD4^+^ T_sen_ (transduced by triple lentiviral depletion of sestrins) and cultured as in (A) (left). Donor-matched CD27^+^ CD28^+^ CD4^+^ T cells (herafter, T_erl_) were cultured in parallel but activated with anti-CD3 and anti-CD28 (*n* = 5 donors). Cells were cultured over two weeks with restimulation every 7 days. DOS-driven population doublings (right) were calculated as delta between DOS treated and DOS untreated T cells, with or without depletion of sestrins. (C) DOS-juvenation of human CD4^+^ T cells. Terminally differentiated effector memory CD45RA^-^ CD28^-^ CD27^-^ CD4^+^ T cells (hereafter, T_EM_) were purified and cultured over 20 days, as in (A). At day 1 (18 hours), 7, and 21 cell phenotypes were assessed by flow cytometry. Quantifications of rejuvenated stem like (CD28^+^ CD45RA^+^ CCR7^+^ CD95^+^ CD62L^+^ TCF1^+^) among human CD4^+^ T cells are shown (left; *n* = 5 donors). Decay of T_EM_ and CD28^-^ CD27^-^ CD45RA^+^ CD4^+^ T cells (hereafter, T_EMRA_) undergoing rejuvenation is shown (right). (D) Adoptive transfer of DOS-juvenated T cells, experimental design. Donor T cells were derived from twenty-month-old mice 15 days after Fluad vaccination with or without DOS treatment (0.1 mg/Kg throughout), labeled with Cell Trace Violet (CTV) dye or congenic CD45.1 tracking, then transferred into young naïve CD45.2 recipients (3 months). In parallel, young mice (3 months) were used as young donor control. Recipient animals were rested for 28 days, then analysed for donor T cell persistence and maintenance of stem phenotype after transfer. (E) Maintenance of donor DOS-juvenated CD45.1 CD4^+^ T cells, their aged-matched controls, and that of young donor T cells, 28 days after transfer (day 43) in recipient mouse lymph nodes. Representative flow cytometry plots and poled data (*n* = 5 mice per group) are shown. (F) Assessment of mouse T cell memory programs in stem like transferred T cells (among CD45.1 CD44^-^ CD62L^+^ CD95^+^ CD4^+^ T cells) and terminally differentiated cells (TE, among CD45.1 CD44^-^ CD62L^-^ CD4^+^ T cells) following adoptive transfer as in (D) (*n* = 5 mice). (G) *IL7R* gene expression and (H) CD95 and Sca-1 mean fluorescent intensity (MFI; throughout) in stem cells derived from CD4^+^ CD45.1^+^ transferred stem T cells among lymph nodes of recipient CD45.2 mice, 28 days after transfer (*n* = 5 mice). (I) Representative plots of CD45.1 transferred cells in quiescent state before and after transfer assessed by cycle related intra-nuclear Ki67 staining. Representative of *n* = 5 mice per group. (J) G1 (Ki67^+^) to G0 (Ki67^-^) transition in stem like CD45.1 CD4^+^ T cells before and after adoptive transfer as indicated (*n* = 5 mice per group). (K) Assessment of T cell longevity following DOS-juvenation. Cells from lymph nodes were stained using the Annexin-PI Apoptosis detection Kit 28 days after transfer. Representative FACS plot (left) and quantification of dead CD45.1^+^ CD4^+^ transferred T cells (right). (L) DOS-juvenated T cell maintenance, *in vivo* model. In (A and B, right) two tailed paired T test was used. In (B, left-C, E-H and J-K) one-way Anova with Bonferroni post-correction for multiple comparisons was used, *p<0,05, **P<0,01; ***P<0,001; ****P<0,0001. Error bars indicate SEM.

To investigate immunological aspects, we derived CD27^-^ CD28^-^ CD4^+^ effector human memory T cells that lack CD45RA expression (T_EM_), which, along with CD27^-^ CD28^-^ CD4^+^ re-expressing CD45RA T cells (T_EMRA_), compose the highly differentiated senescent human T cell subset (T_sen_). These cells co-express the highest levels of sMAC^7^, β-galactosidase and p16, as revealed by flow cytometry with beta-gal and p16 antibodies among sMAC^+^ forming T_EM_ cells (p-p38^+^-sesn2^+^ T_EM_; Figure S4E). Strikingly, senescent T_EM_ cultured with DOS as above, switched to a CD45RA^+^ CD28^+^ CD62L^+^ CD95^+^ TCF1^+^ CD4^+^ stem like memory state^33^ that appeared among the naïve CD45RA^+^ CD28^+^ CCR7^+^ CD4^+^ T cell subset. Stem like switches were evident beginning from 18 hours in culture and kept growing, steadily, throughout the 21-day culture; in these experiments, DOS-induced stem generation likely occurs through a two-step reaction, first restoring CD45RA expression from T_EM_ (intermediate T_EMRA_ stage) followed by complete effector conversion into stem like T cells (Figure 3C). These data reveal that rejuvenation of terminally differentiated senescent T cells manifests as stem like T cell generation (DOS-juvenation). Correspondingly, upon DOS treatment, rejuvenated T cells demonstrated evidence of re-activation of well recognized stem-related pathways, including β-catenin signaling^33,34^ in isolated T cell cultures (without APCs; Figure S4F), as well restored telomere transfer^35^ from APCs at the Tsen synapse in immunological conjugates with subsequent telomere elongation (fluorescence-based APC telomere probe binding transfer coupled to quantitative polymerase chain reaction (qPCR); Figures S4G and S4H). Telomerase activity was not affected (Figure S4I).

Of note, DOS-driven T cell rejuvenation did not cause cell damage or death (Annexin-propidium iodide (PI) staining; Figure S4J). Finally, we used a Seahorse analyzer, and found that DOS treatment restored fatty acid oxidation (FAO) in T_sen_^36^, which is responsible for stem and juvenile maintenance in T cells. Importantly, these effects required sMAC disruption because DOS-driven metabolic reprogramming was not seen in T_sen_ that had been depleted all three sestrins by shRNAs prior to DOS treatment, and sestrin-null T cells also phenocopied the metabolic effects of the drug. These data further support DOS-driven rejuvenation of T cells upon sMAC dismantling, as previously observed in proliferating T_sen_ cultures exposed to the drug (Figure 2B *vs* Figure S4K).

Next, we studied the effects of DOS-juvenation of T cells *in vivo*. An adoptive transfer of old murine CD4^+^ T cells, with congenic surface tracking (CD45.1) or Cell Trace Violet (CTV) labelling^37^ in CD45.2 recipients, revealed that treatment with DOS restored T cell lifespan *in vivo* (Figures 3D and S5). In these experiments, old animals were treated with or without DOS in the presence of heat-inactivated flu vaccines (Fluad), rested 15 days, then donor T cells derived and transferred in young naïve recipients that were not vaccinated. A month later, both young and DOS-juvenated donor T cells were recovered in increased proportions (and similar numbers) in recipients, compared to old donor T cells not exposed to DOS treatment (Figure 3E). These cells had also switched to a CD44^-^ CD62L^+^ CD95^+^ stem like memory state (similar to young, transferred cells), with reduced numbers of CD44^-^ CD62L^-^ terminal effectors (TE) among the CD45.1 CD4^+^ T cells *in vivo*, as assessed by flow-cytometry phenotypic analysis (Figure 3F) and restoration of IL7-receptor transcripts as detected by qPCR (Figure 3G) and further stem like marker expression (CD95 and Sca-1; Figure 3H). Such *in vivo* stem-like memory T cell generation is totally in line with observation of DOS-driven telomere transfer at the immune synapse with APCs (Figure S4G), as recently described for this population of T cells^35^.

When transferred, most donor cells lacked expression of the cell cycle related Ki67 nuclear antigen and persisted in this resting state 28 days after transfer (Figure 3I). That is, donor T stem cells were quiescent at 28 days post transfer *in vivo* and were identified among the G_0_ T cell compartment by the intracellular flow cytometry staining, as Ki67 deficient cells (Figure 3J). Importantly, in the absence of DOS, old, transferred T cells largely died due to necrosis (PI^+^ Annexin^-^ T cells; Figure 3K). Therefore, absence or maintenance of T cells are not due to changes in proliferation after transfer. Instead, gain in survival (longevity) determines the persistence of the rejuvenated T cell in the host. Similar results were seen with adoptive transfer of old CTV CD4^+^ T cells into young recipients (Figure S5). We conclude that DOS-juvenation generates long-lived stem like memory T cells that are largely quiescent *in vivo* (Proposed DOS-juvenated T cell quiescent state; Figure 3L).

### DOS-juvenated T cells orchestrate renewed immunity

To study the impact of DOS-juvenation of T cells on aged immunity, we transferred (CD45.1) DOS-juvenated CD4^+^ T cells (DOS_juv_ T cells), old mouse CD4^+^ T cells or all other DOS treated, CD4^+^ depleted immune cells, into age-matched (CD45.2) recipients vaccinated with Fluad (Figure 4A). Young CD4^+^ T cells were also transferred, as a control. Moreover, young and old and DOS-treated, transfer-free animals were also vaccinated in parallel.

**Fig. 4.**
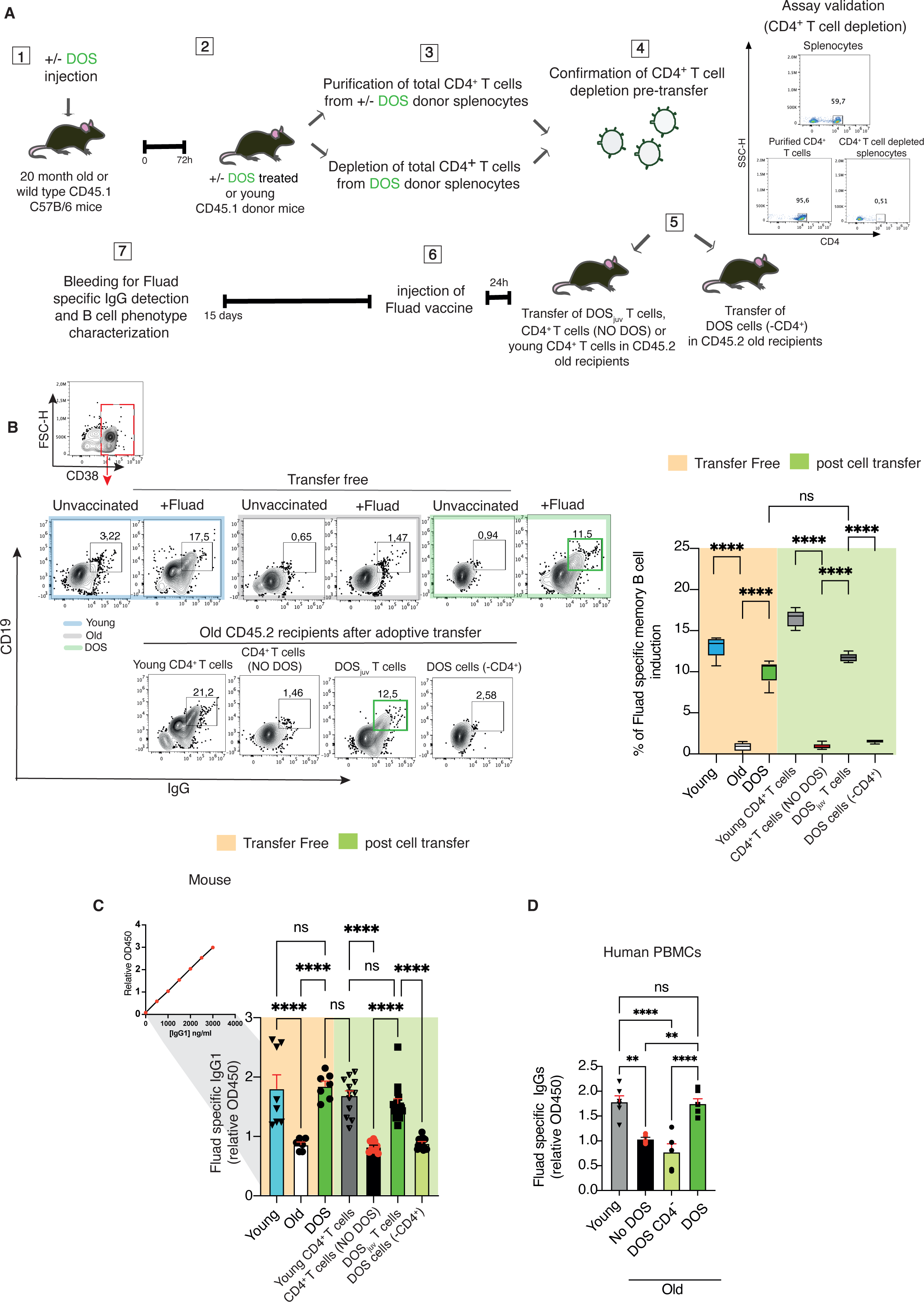
CD4^+^ T cell reliance of DOS. (A) Experimental design. Twenty-month-old CD45.1 mice were injected or not with DOS and sacrificed 72 hours later. Purified CD4^+^ T cells or all other CD4^-^ immune cells were derived from total donor splenocytes. The cells were then transferred into age-matched, old CD45.2 recipient mice vaccinated with Fluad after transfer. Young CD4^+^ T cells from naïve animals were transferred in parallel and compared against DOS_juv_ CD4^+^ T cells. Fifteen days later, Fluad-specific serum IgG1s and lymphoid B cell phenotypes were assessed. Young (3 months), old and DOS treated (transfer) mice were also vaccinated, and shown throughout as a control. (B) Fluad-specific induction of memory CD19^+^ IgG^+^ B cells in vaccinated old recipients subjected to adoptive transfer of DOS-juvenated CD4^+^ T cells (DOS_juv_ T cells), their DOS CD4^+^ T cell depleted splenocytes (DOS cells (-CD4^+^)), their aged matched CD4^+^ T cells from old donor animals not exposed to the DOS (CD4^+^ T cells (No DOS)) or young CD4^+^ T cell transfer from naïve donors. Animals with and without vaccination but not subjected to adoptive transfer are shown, with or without DOS treatment (transfer free). Memory cells were pre-gated on CD38, as shown. Representative plots (left) and pooled data from (*n* = 5 mice per group) are shown. Lymph node cells were studied. (C) Circulating Fluad specific IgG1 in the recipient immunized animals were assessed by ELISA 15 days after vaccination. Data are from *n* = 8 mice (transfer-free; young), *n* = 6 mice (transfer-free; old), *n* = 7 mice (transfer-free; DOS), *n* = 10 mice (CD4^+^ T cells (NO DOS)), *n* = 13 mice (DOS cells (-CD4^+^)), or *n* = 15 mice (DOS_juv_ T cells), or *n* = 12 mice (young CD4^+^ T cells) are shown. (D) Human PBMCs from old donors (age 65-80) were depleted or not of CD4^+^ T cells. Cells were then cultured for 14 days with Fluad vaccines in the presence or not of DOS. ELISA assay was performed as above described. Data from *n* = 6 donors (young), *n* = 5 donors (old No DOS), *n* = 5 donors (old DOS CD4^-^), *n* = 6 donors (old DOS) are shown. In (B-D) one-way Anova with Bonferroni post-correction for multiple comparisons was used, **P<0,01; ***P<0,001; ****P<0,0001. Error bars indicate SEM.

When we assessed reactive mouse lymph-nodes two weeks after Fluad vaccination, only DOS_juv_ T cells or young CD4^+^ T cells restored Fluad-specific CD19^+^ CD38^-^ GL7^+^ germinal center (GC) B cell (Figure S6A), CD19^-/low^ CD45R^-/low^ (B220) CD138^+^ plasma cell^38^ (Figure S6B), and isotopic switched CD38^+^ CD19^+^ IgG^+^ memory B cell generation^39^ (Figure 4B) in the old recipients. By contrast, CD4^+^ T cells not exposed to DOS had no effect. Similar results were obtained throughout in transfer-free animals, when assessing B cell responses from young or DOS-treated mice (but not their aged-matched DOS-free counterparts) immunized with Fluad vaccines (transfer-free controls). Among the DOS_juv_ (or young) transferred T cells, PD1^+^ CXCR5^+^ T follicular helper^40,41^ (Tfh) cells, known to support humoral responses (Figure S6C). Moreover, when we cultured mouse B cells with Fluad vaccines and congenic CD4^+^ T cells that had been depleted of all sestrins via triple shRNA delivery prior to DOS exposure *in vitro*, the drug had no effect on plasma cell generation in culture, showing that DOS-driven plasma cell generation was CD4^+^ T cell sestrin dependent (Figures S6D and S6E). Correspondingly, when we examined serum of the vaccinated animals by ELISA, we found that Fluad-specific immunoglobulines (IgG1s) were restored by the DOS-juvenated (or young) CD4^+^ T cells in the vaccinated recipients subjected to adoptive transfer (Figure 4C). These responses were comparable with transfer-free, vaccinated young mice (∼2000 ng/ml flu-specific circulating IgG1s), or those vaccinated animals that had been systemically exposed to DOS treatment once, with no adoptive transfer of T cells (Figure 4C). By contrast, DOS-free CD4^+^ T cell or DOS-treated, CD4^+^ T cell depleted immune cell transfer had no effect.

Of note, DOS also restored Fluad-specific IgG production in human immune cultures from aged individuals to levels observed in young, vaccinated cultures. As per mouse protocols, these effects were also CD4^+^ T cell dependent because removal of human CD4^+^ T cells by CD4 microbead depletion prior to DOS treatment, abolished DOS function and nullified Fluad-specific IgG restoration, as detected by ELISA in the human immune culture supernatants (Figure 4D). Taken together, CD4^+^ T cells are essential for DOS action upon vaccination.

To further study DOS_juv_ T cell function *in vivo*, we repeated CD45.1 T cell adoptive transfer experiments into aged-matched CD45.2 recipients vaccinated with Fluad and subjected to a lethal flu challenge two months later (Figure 5A). Young and old CD4^+^ T cells not exposed to DOS were also transferred, in parallel infections. Old mice receiving DOS_juv_ or young T cells were fully protected from a lethal infection, whereas those receiving old CD4^+^ T cells or those vaccinated without adoptive transfer largely succumbed to the infection (Figure 5B). As expected, transfer-free, vaccinated young mice could also resist to the infections, and showed responses largely overlapping with vaccinated old animals subjected to DOS_juv_ CD4^+^ T cell transfer.

**Fig. 5.**
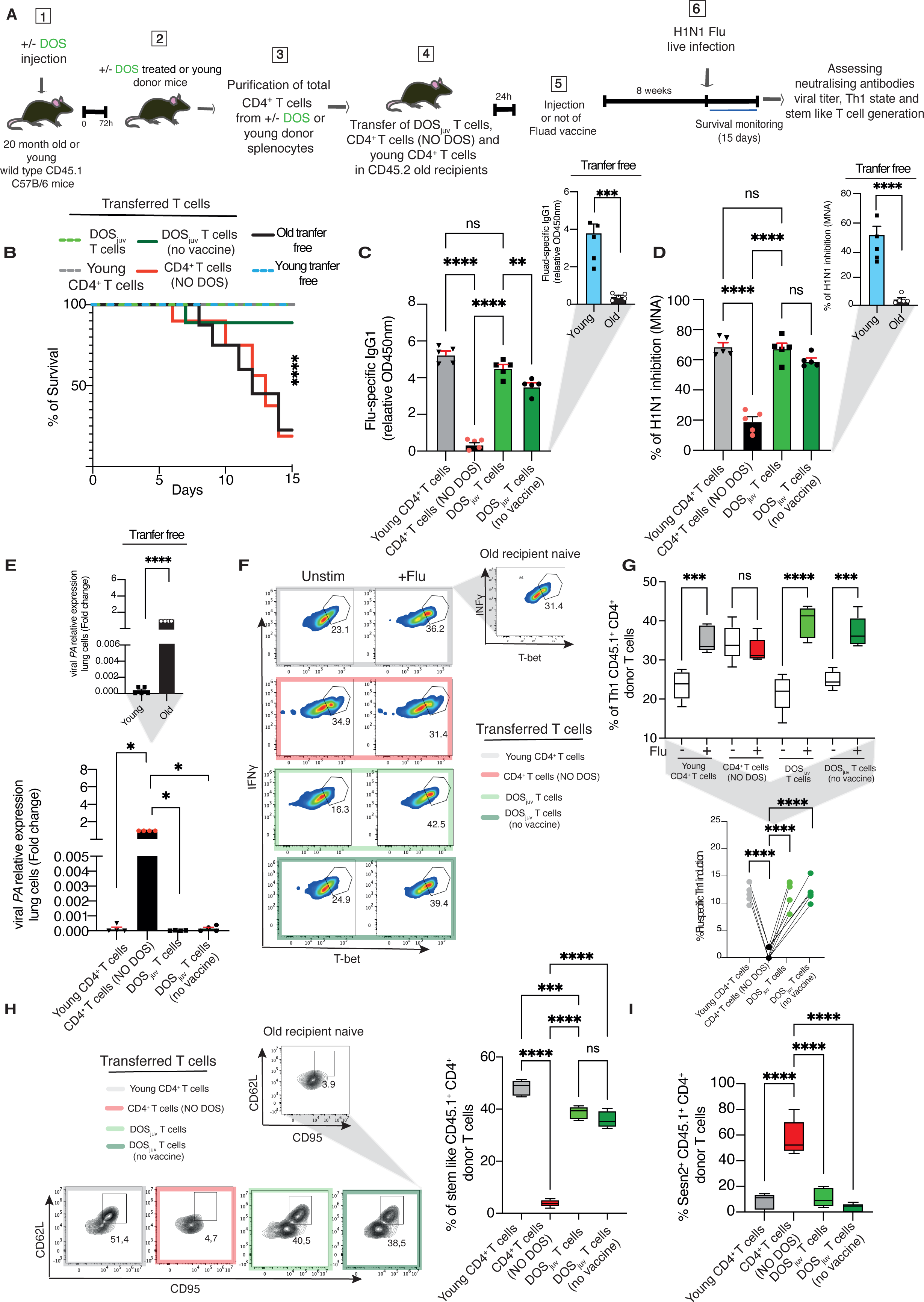
DOSjuv CD4^+^ T cell control of immunity with or without vaccination. (A) Adoptive transfer of DOS-juvenated T cells, experimental design. CD45.1^+^ CD4^+^ donor T cells were derived from twenty-month-old mice injected with and without DOS (0.1 mg/Kg throughout), then transferred into old naïve CD45.2 recipients (20 months) 72 hours later. In parallel, young (3-month-old) naïve CD45.1 mice were used as young CD4^+^ T cell donor control. Recipient animals were vaccinated, or not, with Fluad the day after, as indicated. Animals were subjected to a lethal H1N1 flu viral infection 8 weeks after transfer, and culled within 15 days of infection. Vaccinated young and old mice were also assessed, to determine background responses. (B) Survival of mice, Fluad vaccinated, or not, and then challenged with lethal H1N1 infection as in (A). Data are from *n* = 5 mice per group. (C) Fluad-specific IgG production in the same animals subjected to H1N1 infection. Data are from *n* = 5 mice per group. Transfer free vaccinated recipients are shown (top right; throughout). (D) Neutralization properties of Flu-specific IgGs from recipient mice as indicated were assessed upon overnight H1N1 infection of canine kidney cells (MDCK) *in vitro* by ELISA with anti-influenza A NP antibody. Data are from *n* = 5 mice per group. (E) Viral polymerase acidic protein (PA) assessment in lungs of the same mice upon H1N1 viral infection as in (A) by qPCR (*n*=5 mice per group). Data are expressed as fold change to the old infected control (transferred CD4+ T cells, no DOS), set as 1. (F) Representative FACS plots and (G) pooled data (*n* = 5 mice per group) of terminal primary antiviral Th1 responses among T-bet^+^ IFNγ^+^ CD45.1 donor mouse CD4^+^ T cells as in (A). Data are from *n* = 5 mice per group. (H) CD44^-^ CD62L^+^ CD95^+^ CD4^+^ T cells (stem-like T cells) in animals subjected or not to CD4^+^ T cell transfer and then infected in the same experiments as in (A); Old naive recipient examples are shown (*n* = 5 mice per group). (I) The fraction of sestrin expressing donor T cells among adoptively transferred CD45.1^+^ CD4^+^ T cells from age-matched (20 months) CD45.2^+^ recipients, six months after transfer as assessed by flow cytometry (*n* = 5 mice per group). In (C-E and G-I) one-way Anova with Bonferroni post-correction for multiple comparisons was used. In (C-E, top right), a two tailed paired T test was used. In (B) Mantel-Cox test was used. **\***P<0,05; **\*\***P<0,01; **\*\*\***P<0,001; **\*\*\*\***P<0,0001. Error bars indicate SEM.

Further examination revealed that DOS_juv_ T cells triggered H1N1 viral specific antibody production (ELISA assessment; Figure 5C), with neutralising activity (Micro-neutralisation assays with MDCK cells; Figure 5D) and eradication of viral titres *in vivo* (qPCR; Figure 5E). DOS_juv_ T cells also induced antiviral Th1 cell responses in the infected lungs (flow-cytometry; Figures 5F and 5G), well recognised events that confer anti-viral protection and sterilising immunity^42^. Moreover, flu-specific stem-like T cells were also observed among the transferred T cells that were exposed to the DOS, as assessed by flow-cytometry in the recipient lymphoid organs (spleens; Figure 5H).

In parallel, we also tested the capacity of DOS_juv_ T cells to control infections in identical transfer experiments where the old recipients were infected with no prior vaccination. Surprisingly, DOS_juv_ T cells controlled flu viral infections, in a way that was much similar with their function in a vaccinated host, albeit slightly less effectively than when combined with the vaccine (Figures 5B-5H). Furthermore, we found evidence of long-term (6 months) sestrin inhibition within the adoptively transferred CD45.1 DOS_juv_ T cells (Figure 5I), consistent with long-term T cell rejuvenation. These results were T cell autonomous because only the transferred CD45.1 T cells, but not the old CD45.2 recipients, had been exposed to the drug.

Similar results were obtained by subcutaneous DOS administration that protected vaccinated old animals against a lethal infection, restoring persistent flu-specific IgGs (Figure 6A to Figure 6D), with neutralizing activity (Figure 6E), even 6 months after the single DOS treatment. These experiments confirmed that DOS-deprived, aged humoral responses are defective *in vivo*, which can be reversed by the DOS. Stem like T cell induction was also restored by systemic DOS treatment *in vivo* (Figure 6F).

**Figure 6.**
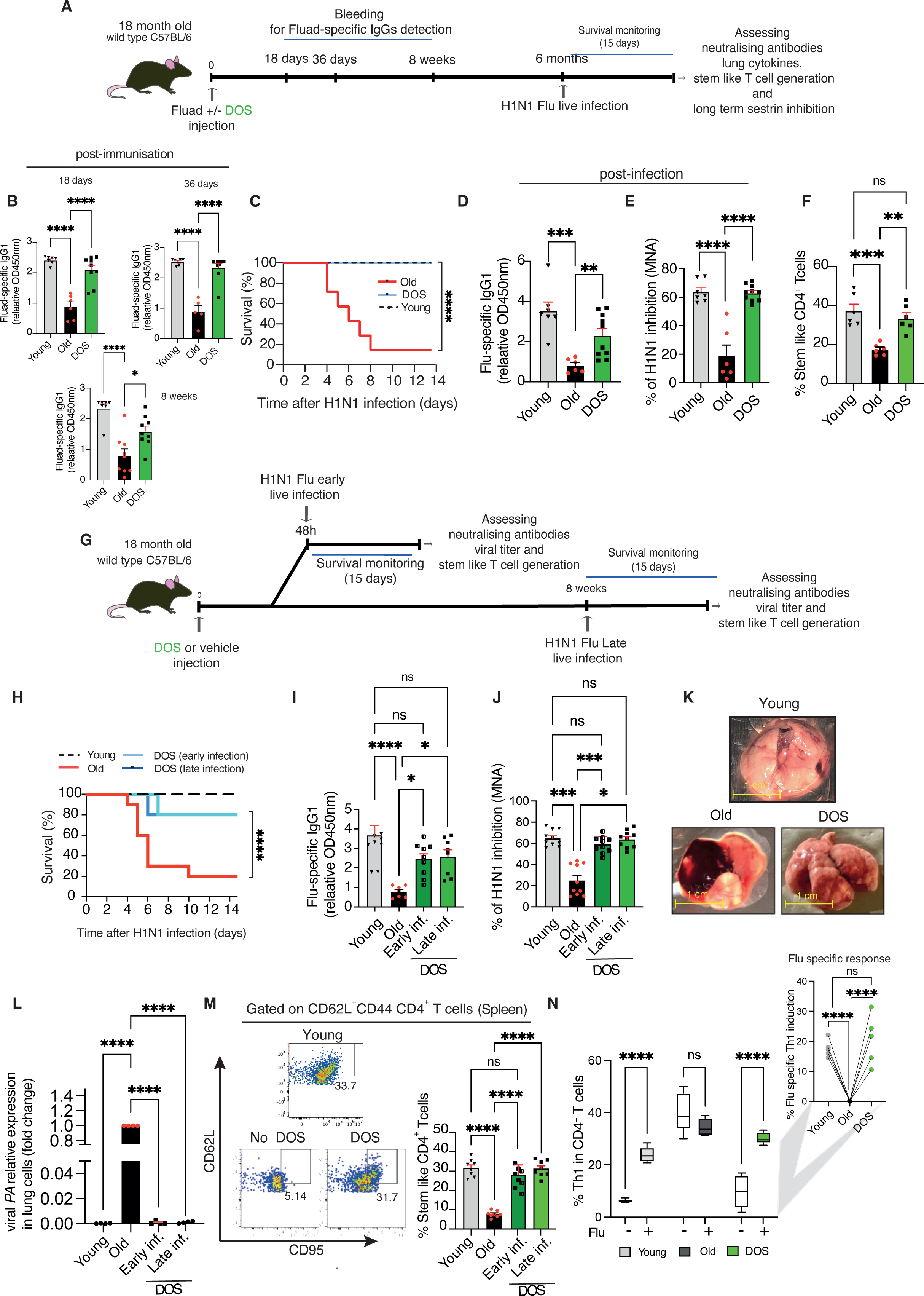
Phenocopy of DOS_juv_ CD4^+^ T cell transfer by systemic DOS administration. (A) Experimental design. Eighteen-month-old mice were vaccinated with Fluad and injected or not with DOS. After 18, 36 days and 8 weeks mice were bled to detect Fluad specific IgG production. Six months later, mice were infected with lethal H1N1 flu virus and then monitored for survival. Mice were culled within 15 days. Young mice (3 months) were used as control. (B) Antigen-specific IgG production in young, old and DOS-juvenated mice upon Fluad immunization. For 18 days post-immunization, data are from *n* = 7 mice (young), *n* = 6 mice (old), *n* = 9 mice (DOS). For day 36, data are from *n* = 7 mice (young), *n* = 5 mice (old), *n* = 9 mice (DOS). For week 8, data are from *n* = 7 mice (young), *n* = 10 mice (old), *n* = 10 mice (DOS). Mice were bled at the indicated time points and circulating serum vaccine-specific IgG levels were detected by ELISA. (C) Survival of mice, vaccinated and challenged with a lethal H1N1 infection, 6 months later as in (A) at 24 months of age. Data are from *n* = 7 mice per group (young and old) and *n* = 8 mice (DOS). (D) Fluad-specific IgG production in the same animals 15 days after H1N1 viral infection. Data are from *n* = 7 mice (young), *n* = 6 mice (old) or *n* = 10 mice per group (DOS). (E) Neutralization properties of Flu-specific IgGs were assessed upon overnight H1N1 infection of canine kidney cells (MDCK) *in vitro* by ELISA with anti-influenza A NP antibody. Results with heat-inactivated sera from DOS-juvenated mice, their aged matched controls without DOS, and young animals are shown. Data are from *n* = 7 mice (young), *n* = 6 mice (old) or *n* = 10 mice per group (DOS). (F) Terminal assessment of CD44^-^ CD62L^+^ CD95^+^ CD4^+^ T cells (stem-like T cells) 231 days after DOS treatment, in splenocytes from old animals infected as in (A). Aged matched and young infected animals not exposed to the DOS are shown (*n* = 6 mice per group). (G) Experimental design. Eighteen-month-old mice were injected with or without DOS, in the absence of Fluad vaccination. Mice were then infected with H1N1 virus after 48 hours, or, in parallel, after 8 weeks to assess survival and anti-viral immune responses to either early or late infection, respectively. Young mice (3 months) were used as control. (H) Survival of animals as in (G) was assessed (*n* = 10 mice). (I) Detection of Flu-specific IgGs among the same animals infected in the early and long-term protocol. ELISA plates were coated with H1N1 viral extracts, rather than Fluad, to detect viral specific antibodies. Data are from *n* = 10 mice (young), *n* = 7 mice (old), *n* = 8 mice (DOS early infection (inf. throughout)) or *n* = 9 mice (DOS late inf.). (J) Neutralization properties of Flu-specific IgGs from DOS-juvenated mice serum, age-matched infected animals not exposed to the DOS or young, infected animal serum (as in G), with no prior immunization, as in (I). (K) Images of autoptic lungs from representative old mice treated with or without DOS then infected with H1N1 8 weeks later. Young, infected lungs are shown. Organs were collected 6 days after infection. Note absence of lung necrosis with the DOS only regimen (no vaccination). (L) Viral polymerase acidic protein (PA) assessment in lungs of young, old or DOS juvenated mice upon H1N1 viral infection (as in A; *n*=4 mice per group). Data are expressed as fold change to the old infected control (no DOS), set as 1. (M) Assessment of stem-like T cells upon DOS juvenation after H1N1 viral infection in the early and long-term protocol. Representative FACS plots (left) and quantification (right, *n*=8 mice per group). (N) Primary antiviral Th1 responses among T-bet^+^ IFNγ^+^ CD4^+^ T cell of young, old or DOS juvenated mice to H1N1, 3 days after infection (*n* = 5 mice). In (B, D-F, I-J and M, right-N) one-way Anova with Bonferroni post-correction for multiple comparisons was used. In (C) and (H) Mantel-Cox test was used. *P<0,05; **P<0,01; ***P<0,001; ****P<0,0001. Error bars indicate SEM.

Likewise, DOS only, vaccine-free prophylaxis protocols were effective at protecting old animals against a lethal (future) infection when the flu virus was inoculated either immediately (2 days) or 2 months after the single DOS treatment (Figure 6G and 6H) and elicited analogous flu-specific neutralizing antibodies (Figure 6I and 6J). Protective responses were also visualized in the lungs from animals that had been exposed to the DOS prior to infection, with no sign of lung necrosis (Figure 6K) and absence of viral particles among the infected lungs demonstrating viral clearance *in vivo* (qPCR; Figure 6L). Moreover, both stem-like T cell induction (Figure 6M) as well flu-specific T cell responses (Th1; Figure 6N) could be restored after infection to the levels found in young mice not exposed to the drug. Taken together, DOS treatment leverages on CD4^+^ T cells to confer long-term immune protection against infection *in vivo*.

Immune-protection may be also linked with long-term T cell rejuvenation evident up to 6 months after the single DOS treatment and manifested as persistent stem-like T cell induction and sestrin downregulation to the levels found in CD4^+^ T cells derived from young animals. This latter was observed consistently both at the transcript and protein level by qPCR, flow cytometry and immunoblotting of sestrins in peripheral T cells that had been recovered from blood and spleen (Figures S7A-S7D). These effects are likely consequence of long-term deacetylation^43^ of the *sestrin* gene, as assessed by T cell Chromatin immunoprecipitation with anti-histone antibodies coupled to qPCR based *sestrin* gene amplification (Figure S7E) and enhanced histone deacetylase activity (HDAC1; Figure S7F). In these experiments, there was reduced *sestrin* gene amplification among acetylated T cell H3 histones derived from mice subjected to single dose systemic DOS treatment. Furthermore, when we incubated the immunoprecipitated T cell HDAC1 from these mice with aged-matched T cell DNA from DOS free animals, we confirmed direct histone 3 deacetylation that is associated with *sestrin* gene inhibition, as revealed by ELISA assays (anti-acetil-H3K27; Figure S7F). Additionally, systemic DOS treatment supported Tfh responses upon vaccination, further analogy with earlier adoptive transfer of DOS_juv_ T cells (Figure S8 *vs* Figure S6C). Therefore, DOS-juvenation of T cells can occur with or without vaccination and strongly reinforces the adjuvanted clinical vaccine alone, resulting in long term immune rejuvenation.

Searching for a mechanism of immune protection by the rejuvenated T cells, we discovered increased levels of T cell receptor (TCR) excision circles (TRECs) ^44^ among lymph nodes of flu infected, DOS treated old animals by real time PCR (Figure S9A), suggesting recent DOS-driven TCR rearrangements with new specificities^45^. Accordingly, flow cytometry examination revealed that DOS-juvenated stem cells from these infected animals demonstrated expression of new TCRβ variants (Figure S9B). As such, augmented TREC generation was also detected in adoptively transferred DOS_juv_ CD45.1 CD4^+^ T cells recovered from infected old CD45.2 animals (Figure S9C), as well in purified human CD4^+^ T cells exposed to DOS treatment in culture (Figure S9D). These data indicate that DOS treatment induces TCR rearrangements in mature lymphocytes, and in a cell autonomous way, suggesting a new means for immune protection by DOS_juv_ CD4^+^ T cells, via immune repertoire expansion.

To test if DOS-juvenation of T cells expands antigen recognition, we returned to the human immune system and used yellow fever (YF), to which Western populations have not been exposed^46^. We exposed, or not, human CD4^+^ T cells to DOS and cultured them with autologous APCs pulsed with different concentrations of YF vaccines, using antigen free cultures as a control. We found that DOS_juv_ T cells specifically produced IFN-γ, exclusively in the stem like comportment that had been exposed to the YF vaccine, indicating generation of new YF-specific responses by the rejuvenated T cells (Figure S9E). This response reminded that of sestrin deficient T cells with increased TCR sensitivity such that they best act at lower antigen concentrations^7^. However, the new YF specific responses could not be simply explained by increased TCR sensitivity. In fact, in these experiments, T cells that had not been exposed to DOS treatment remained always unreactive, regardless of the relative YF antigen dosing (over 50-fold dosing; Figure S9E).

To further test induction of YF specific responses, we treated CD4^+^ T cells with or without DOS, followed by conjugation with YF pulsed APCs, as above. After overnight culture, we derived purified stem like, CD45RA^-^ CD28^-^ effector memory (TEM) and CD45RA^-^ CD28^+^ central memory (TCM) CD4^+^ T cells from the cultures, rested cells for 5 days, and performed YF restimulation with autologous APCs. Supernatant analysis confirmed IFN-γ production and release by the same YF specific stem T cells exposed to DOS (but not in the supernatants from the DOS free cultures; Figure S9F). By contrast, T_EM_ or T_CM_ from the same cultures had no YF specific response, even in the presence of DOS.

Similar YF specific stem-like T cell responses could be also detected using specific YF peptide loaded APCs, instead of the vaccine itself, by flow-cytometry (absolute antigen specific induction; Figures S9G and S9H), whereas an irrelevant peptide had no effect. Furthermore, de novo generation of DOS-driven YF specific responses could be detected in stem-like CD4^+^ T cells derived from OT-II mice. These transgenic animals express only an ovalbumin (OVA)-specific TCR, indicating that any new response needs to derive from purposely recombined TCRs (Figure S9I). Thus, DOS treatment can confer new antigen specificity both in the presence or in the absence of vaccination that may result in prophylactic activities for antigens that will be encountered in the future. These data are suggestive of *de novo* TCR recombination, extending previous observations of TCR revision programs in peripheral T cells^45,47–49^.

We then performed next generation sequencing (NGS)^50^ and examined the TCR β gene in human T cells exposed, or not, to DOS and conjugated with autologous, YF peptide pulsed APCs. Antigen free cultures were used as control, to derive antigen-driven rearrangements upon background subtraction, with or without DOS.

Eighteen hours later, we compared the TCR β reads amplified by the sequencing in both DOS exposed and DOS free T cells derived from the YF cultures. We found rapid induction^51^ of YF related TCR revision programs (Figure 7A), consisting of enhancement of TCR repertories and, strikingly, generation of new TCRs with antigen pockets (DJ recombination) that did not exist before DOS treatment (Figure 7B). Individual TCR rearrangements were also detected by the sequencing upon antigen stimulation in cultures where T cells had not been exposed to the DOS yet to a lower extent (Figure S10A); however, TCR β pocket rearranging largely required presence of DOS. Importantly, examination of TCR clonotypes revealed DOS-driven rearrangements to be exclusively evident amongst the rarest TCR β variants (Figure S10B), suggesting occurrence in rare populations such as antigen specific stem cells.

**Figure 7.**
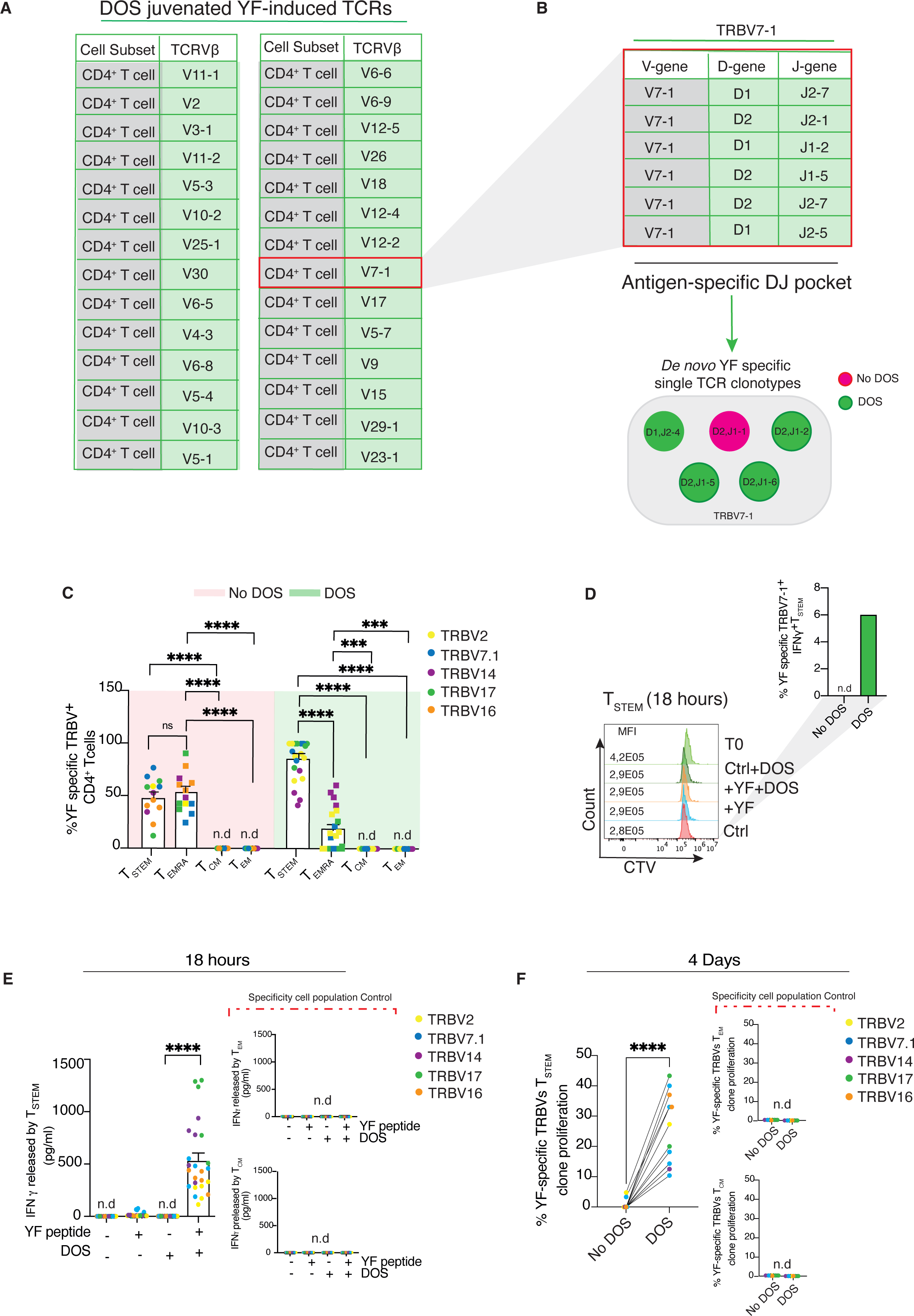
DOS-juvenated T cells present *de novo* antigen-specific TCR rearrangements. (A) Purified human CD4^+^ T cells and autologous APCs (CD3-depleted PBMCs) were pre-treated or not with DOS or YF peptides (YF), respectively, then co-cultured 18 hours in 1:2 ratio. The day after, post-synaptic CD4^+^ T cell genomic DNA was derived from the conjugates and processed for Next generation sequencing (NGS) of the *TCRVB* gene. The table shows inductions of *T cell receptor Beta Variant* (TRBV) rearrangements (V-DJ recombination) from a single donor upon YF stimulation of DOS-juvenated CD4^+^ T cells. Unstimulated backgrounds were subtracted. For experimental details, please see Methods. One representative donor of 4 is shown. Further genomic clonotypes, Supplementary information 10. (B) Antigen pocket rearrangements among TRBV7-1 variant in T cells derived from the same donor as in (A). Example of single YF-specific TCR clonotypes (among TRBV7-1) that are only seen in T cells exposed to YF-pulsed APCs, with or without DOS (bottom). Note that most antigen pocket rearranging requires presence of DOS, indicating active D-J recombination. (C) Single cell presence of YF-specific TRBVs among the different T cell subsets of the 4 donors subjected to NGS. The expression levels of each TRBV were evaluated in individual cells by flow cytometry against the unstimulated background treated with DOS (to isolate YF-specific TCRs induced by the DOS); or the background treated without YF and DOS (to isolate YF specific TCRs in cells that are not subjected to DOS-juvenation). Each dot is a single TRBV in the single cell flow cytometry assessment. Note absence of YF-specific TRBVs in T_CM_ and T_EM_ cells. Results from 4 experiments are shown. (D) Absence of early YF-specific proliferation (18 hours) of the same stem T cells as shown by CTV tracking. An example of functional, YF-specific TRBV (7.1) clonotypes among the same stem-like T cell clones, at the same time, is shown (intracellular flow IFN-γ detection; top right). CD4^+^ T cells were purified from the same donor as in (A) and conjugated with or without autologous YF-pulsed APCs, in the presence or in the absence of DOS. Antigen free synapses were subtracted. (E) IFNγ release by the indicated TRBV expressing T stem clones after 18-hour YF peptide stimulation and (F) antigen-specific proliferation of the same clones, 4 days after culture as assessed by BrdU incorporation. Cell culture conditions as in (A). Specificity cell population controls (T_CM_ and T_EM_) are shown. In (C and E) One-way Anova with Bonferroni post-correction for multiple comparisons was used. In (F) statistical significance was determined by two tailed unpaired student T test. ***P<0,001; ****P<0,0001. Error bars indicate SEM.

To validate NGS findings, we used specific anti-TCR variant antibodies and flow cytometry and assessed TCR variant usage among the same donor T cells subjected to the NGS. Strikingly, we found that DOS_juv_ T cells, in culture with YF pulsed APCs for 18 hours, effectively expressed the NGS identified TCRs among the stem-like T cell compartment (Figure 7C). Some T_EMRA_ cells also showed new TCR variant expression in these experiments, suggesting TCR revision programs may begin early in the DOS-juvenation reaction, when these cells arise from senescence prior to stem like conversion (Figure 7C). By contrast, T_EM_ or T_CM_ lacked YF-specific TCRs. In parallel, using CTV dilution assays, no trace of YF specific proliferation (CTV dilution induced by the YF antigens) could be detected at this early assessment (Figure 7D). However, these DOS-treated, TRBV expressing T cells produced IFN-γ in response to YF activation at this time, as assessed by flow-cytometry with anti-IFN-γ antibodies among the stem like compartment (Figure 7D top right). Therefore, appearance of new antigen-specificity (e.g. specific IFN-γ production) precedes clonal expansion, further reinforcing that TCR variant expression in DOS_juv_ stem T cells was not due to amplification of rare, previously existing clones.

To further confirm generation of new YF specific responses, each TRBV expressing stem clone was purified by fluorescence activated cell sorting, and re-challenged in antigen specific assays five days later, as above. There was YF-specific IFN-γ release after 18-hour activation (Figure 7E), as well 4-day YF-specific proliferation (Figure 7F), respectively detected by ELISA in the culture supernatants, and BrdU incorporation in the stem like T cell clones that had been exposed to the DOS. Other effector or central memory subsets had no specific responses, confirming that the acquisition of new TCR specificity upon DOS treatment was strictly related to generation of the newly TRBV expressing rejuvenated stem like T cell clones.

To assess for a possible mechanism of TCR revision, we measured the expression of Rag proteins in mature CD4^+^ T cells exposed or not to DOS for 18 hours. Rag proteins have low expression in mature T cell populations^47,52^: however, DOS treatment potently enhanced Rag 1 and 2 protein levels in activated CD4^+^ T cells as detected by immunoblotting with anti-Rag antibodies (Figure S11A). Moreover, we used chromatin immunoprecipitation (Chip) and purified the Rag complex derived from T cells, exposed or not to DOS, and activated by anti-CD3 ligation or autologous, YF-pulsed APCs (following conjugate dissolution and isolation of post-synaptic T cells, as described^37^). With both activations, we found that DOS treatment potently induced Rag binding to the T cell DNA, and in antigen specific way (Figures S11B and S11C).

Next, we generated Rag deficient T cells by direct siRNA delivery, using an irrelevant, non-targeting siRNA as control (flow cytometry validation; Figure S11D). We subjected these Rag deficient T cells to YF-specific activation by autologous APCs, with or without DOS. Parallel antigen free cultures served to derive YF specificity. When we analyzed these cultures by flow cytometry with anti-TRBV antibodies and stem like T cell markers (CD95 and TCF1 among otherwise naïve T cells), as expected, we found that DOS-driven YF-specific TCRs were recombination products because their expression was abolished in these Rag deficient T cells (Figure S11E). Accordingly, generation of YF-specific TRECs (detected by qPCR upon background subtraction), was disrupted by Rag deficiency in these cells (Figure S11F). These responses, which were obtained just with DOS prophylaxis and no prior vaccination of the subject, are in line with previously reported YF specific T cell activities among YF vaccinated individuals^53^, similarly to vaccine free, prophylactic treatment of T cells or mice (DOS only *in vivo*; Figure 5 and Figure 6). Specific rearrangements were also obtained with common Epstein Bar viral (EBV) antigens and overlapped in part with YF activation (Figure S12A), further suggesting generation of stem TCRs with flexible pockets against a variety of challenges, in analogy with recently described ‘sponge’ TCRs^54^ that can protect against multiple insults. As such, generation of different DJ antigen pockets correlated with acquisition of *de novo* antigen-specific function by the productive TRBV stem like T cell clones (*de novo* IFN-γ production; Figure S12B). We conclude it is possible to use DOS and rejuvenate senescent T cells, generating new stem like T cell clones with expanded TCR specificity. A DOS-juvenation program is proposed (graphical abstract).

## Discussion

Here we described DOS, first in class sMAC disruptors that selectively rejuvenate T cells and allow antigen specific TCR revisions. As such, DOS treatment can confer protection against challenges that are yet to be encountered. Vaccination can also be strongly improved but is not essential for DOS action.

Although our data identify stem-like T cell generation from formerly senescent cells as key pathway to T cell rejuvenation, these findings are not in contrast with recent observations of iteratively stimulated T cells (ISTCs) persisting for years in vivo. Rather, DOS-driven T cell rejuvenation only required one round of activation to access stem state, illuminating stemness as the gateway to access rejuvenation for senescent T cells. In the future, these rejuvenated T cells, with long telomeres and proliferative capacity, may sustain repeated rounds of activation, converging toward immortal-like state, similarly, to recently described ISTCs^55^.

It was thought initially that T cells underwent some form of peripheral TCR revision, to eliminate anergic clones, and to comply with defective expression of the antigen receptor on the cell surface^46–49,56,57^. However, recent studies revealed that such CD3^low^ cells actively participate in immune response^58^ and undergo TCR shedding from their antigen specific synapses to unleash telomere transfer from APCs, initiating an antigen specific stem switch^37^. An exciting possibility would be TCR revision-based chromatin remodeling to be supported by IL7^48^ in the telomere recipient T cells; and this cytokine signaling is also a feature of the stem T cells with new TCRs generated by DOS. It remains to be tested whether DOS effects may be broader than rejuvenation of the immune system. For instance, T cell rejuvenation by sMAC inhibition might trigger additional rejuvenation cascades in non-lymphoid tissues, due to newly arising T cell roles in controlling organ homeostasis *in vivo*^58^. A possible caveat in these approaches could be spontaneous autoimmunity upon DOS-driven TCR rearranging; however, we are yet to see any autoimmunity case in our large pre-clinical cohorts (over 400 animals), perhaps due to thymus reactivation, and elimination of auto-reactive clones. Thus, immunity can be rejuvenated by DOS, and this may initiate new forms of immunotherapies.

**Figure S1 – DOS design**

(A) Schematic representation of DOS compound design strategy.

(B) Workflow of the *in vitro* screening of DOS compounds.

(C) *In vitro* screening of DOS compounds. Suppression of AMPK activation upon addition of recombinant sestrin was used as a readout.

(D) Cyclic DOS46L synthesis, the lead compound.

**Figure S2 –DOS-sestrin2 binding, molecular docking**

(A) 3D representation of sestrin 2 showing the two possible DOS46L binding sites. The interacting amminoacids are shown. Highlighted aminoacids were subjected to inactivating mutagenesis.

(B) Predicted DOS binding site to sestrin 2. Boxes indicate intervention points for mutagenesis.

(C) Confirmation of sMAC disruption sites by DOS was performed by intracellular flow cytometry and showed as fold change to the untreated control (no DOS), set as 1. Alanine scanning mutagenesis was performed in primary human T_erl_ cells (*n* = 3 donors).

**Figure S3 - Exemplification of DOS discovery**

(A) Representative FACS plot (left) and quantification (right) of DOS penetration in human T_sen_ compared to T_erl_ (*n* = 8 donors).

(B) DOS treatment does not alter CD4 internalization in purified human CD4+ T cells followed by anti-CD3/CD28 activation (*n* = 4 donors).

(C) Confocal imaging of sestrin2, DOS46L and the reticulum endoplasmatic KDEL marker. Colocalization in human CD4^+^ T cells and its individual channels are shown. Representative of *n* = 3 donors.

(D) Expression of GATOR2 components (MIOS and WDR59) in T_erl_ (CD27^+^ CD28^+^), T intermediate (T_int_; CD27^-^ CD28^+^) and T_sen_ (CD27^-^ CD28^-^) human CD4^+^ T cell subsets, was assessed by immunoblotting, as indicated. GAPDH, loading control. Representative of 4 experiments with 4 donors. WDR59 in HEK293T, positive control (run in parallel).

(E) sMAC IPs were derived by sestrin 2 pull down from human CD27^-^ CD28^-^ CD4^+^ T cells (hereafter, Tsen) transduced as indicated. The IPs were treated or not with DOS for 30 minutes in the presence or in the absence of ATP (200 μM) prior to ELISA based *in vitro* kinase reactions with anti-p-p38.

(F) p38 activity among sestrin 2 IPs (sMACs) derived from human T_sen_ exposed or not to DOS directly in the IP reaction, in the presence or absence of compound C (10 µM). Note that DOS inhibitory action is phenocopied by compound C and no further inhibition of sMAC can be obtained when combining DOS with the AMPK inhibitor (*n* = 3 donors). *In vitro* kinase assays detecting p38 MAPK self-phosphorylation reactions upon subtraction of ATP free conditions are shown.

(G) Quantification (*n* = 5 donors) of sestrin-AMPK binding within sMACs (wash-in, top) and their release from the complex (wash-out, bottom). Heatmap of sestrin presence in wash-in and wash-out fractions is shown. sMAC complexes were immunoprecipitated with anti-AMPKα antibodies and incubated with or without DOS for 2 hours, followed by ELISA with anti-sestrin antibodies (sestrin 1, 2 and 3) of both wash-in and wash-out fractions in human CD4^+^ T_sen_. Data are expressed as fold change to the untreated control, set as 1.

(H) Expression of T_sen_ sMAC components (sestrin 2 and AMPK) with or without DOS (4 hour exposure), in the presence or in the absence of MG132 (10μM) pre-treatment (30 minutes) assessed by immunoblotting. H3, loading control.

(I, left) AMPK activity measured assessing phosphorylation of its downstream target, Acetyl-CoA carboxylase (ACC). Human T_sen_ were treated or not with DOS for 2 or 4 hours and the levels of phosphorylated (p)-ACC and total ACC were evaluated by flow cytometry. AMPK activity was measured as the ratio between p-ACC and total ACC. Initial AMPK activity (time 0) was set as 1 (100%). Representative plots and pooled data from (*n* = 5 donors) are shown. Compound C treatment was used as control to phenocopy DOS action. (I, right) Confirmation of reduced ACC phosphorylation (Ser79) by immunoblotting, 4 hours after DOS treatment in human CD4+ Tsen. Pooled data from (n = 3 donors) are shown. Data are expressed as fold change to the untreated control (no DOS), set as 1.

(A) (J) Representative plot (left) and quantification (right) of sMAC (sestrin^+^-p-p38^+^) presence among human senescent CD28^-^ CD27^-^ CD8^+^ T cells, assessed by flow cytometry with or without DOS treatment (4 hours; *n* = 6 donors).

In (B, D and E) One-way Anova with Bonferroni post-correction for multiple comparisons was used. In (A, F and H-I) two-tailed paired t-test. *P<0,05; **P<0,01; ***P<0,001; ****P<0,0001. Error bars indicate SE.

**Figure S4 - Effects of DOS-juvenation**

(A) T_sen_ were treated with or without DOS for one week in the presence of anti-CD3 and recombinant human (rh)IL2 (*n* = 5 donors). Representative image (left) obtained by confocal microscopy and quantification (right) of H2AX intensity (MFI, Mean Fluorescence Intensity). Scale bar, 20 µm.

(B) Purified human T_sen_ were activated as in (A) for 48 hours and then depleted or not of all sestrins using lentiviral particles. Five days after transduction, cells were stained with CTV dye, activated as above then cultured for additional 3 days in the presence or absence of DOS prior to CTV dilution analysis. Representative FACS histograms of *n* = 9 donors.

(C) Human T_sen_ were activated and depleted of sestrins, as above. Five days later, proliferation was assessed by BrdU incorporation for 72 hours in the presence or in absence of DOS (*n* = 5 donors).

(D) Effect of DOS in AMPK null senescent T cells shown by CTV staining (*n* = 3 donors). Human T_sen_ were genetically modified to deplete AMPK, cultured for one month and then stimulated in the presence of CTV for another 3 days. Cells were activated every 7 days (4 round of activations as above) prior to CTV staining to examine proliferation of terminally differentiated cells in culture. Representative FACS histograms (top) and quantification of CTV low proliferating cells (below).

(E) Senescent markers (p16 and β-Galactosidase) expression in human terminally differentiated T_EM_ directly *ex vivo*, in sMAC^+^ or sMAC^-^ cells. Representative FACS plot (left) and quantification (right) are shown (*n* = 6 donors).

(F) Levels of stem related intracellular signaling (β-catenin) with or without DOS treatment in human stem-like T cells derived from purified T_sen_ cells (*n* = 4 donors).

(G) Representative FACS plots (left) and quantification (right) of telomere transfer in T_sen_ forming synapses with Ag loaded APCs in the presence or in the absence of DOS. Purified human APCs were labeled with fluorescent telomere probes (Cy3 PNA) and allowed to form synapses with autologous T_sen_ (3:1 ratio; *n* = 5 donors). data are shown as fold change to the untreated control (no DOS), set as 1.

(H) T_sen_ were treated or not with DOS then conjugated with autologous APCs pulsed with Cytomegalovirus (CMV) and Epstein Bar Virus (EBV) antigens (1:50 each) overnight. Post-synaptic T_sen_ and APC genomic DNA was assessed for telomere measurements. Telomere length was calculated from the ΔCT according to the formula suggested by the kit.

(I) Assessment of telomerase activity in T_sen_ 3 days after DOS treatment (*n* = 5 donors). Telomerase activity was detected according to the TeloTAGGG Telomerase PCR ELISA Kit.

(J) Cellular toxicity in human CD4^+^ T cells in the presence or in the absence of DOS as assessed by Annexin V and PI (*n* = 4 donors). H2O2 treatment, positive death control.

(K) Fatty acid oxidation (FAO) levels detected in non-senescent human T_erl_ or in T_sen_ treated or not with DOS (left) or in the presence or in the absence of Sestrins (middle). FAO rate was calculated by etomoxir subtraction. Absence of DOS-driven FAO metabolic reprogramming is shown (right). Data are from *n* = 6 donors.

In (A, E, G-I and K, left) two-tailed paired T test was used. In (C-D and K, right) one-way Anova with Bonferroni correction for multiple comparisons. *P<0,05; **P<0,01; ***P<0,001. Error bars indicate SEM.

**Figure S5 – Phenocopy of CD45.1 T cell longevity *in vivo***

(A) Phenocopy of CD45.1 tracking studies. Representative FACS plots, and frequencies of donor CTV CD4^+^ T cells recovered from young CD45.2 naïve recipients subjected to adoptive transfer as per Figure 2E, 28 days after adoptive transfer (day 43; *n* = 6 mice per group).

Representative pre (B) and 28-day post (C) transfer phenotypes of CD4^+^ CTV^+^ donor T cells from young, old (no DOS) and DOS-juvenated donor mice before adoptive transfer into young recipients. Pre-transfer T cells were assessed 15 days after donor vaccination with Fluad in the presence or absence of DOS as in Fig. 2d. CD95 FMO controls are shown.

(A) (D) Frequency of CD44^-^ CD62L^+^ CD95^+^ stem like T cells and that of terminally differentiated CD62L^-^ CD44^-^ lymphocytes among donor CTV^+^ CD4+ T cells. Donor T cells were analyzed 28 days after transfer (day 43).

(B) (E) Sca1 and CD95 protein levels in CTV^+^ CD4^+^ stem T cells in the same recipient mice, 28 days after cell transfer (*n* = 5 mice per group). MFI values are shown.

(C) (F) *IL7R* gene expression among sorted CTV^+^ CD4^+^ T cells purified form lymph nodes of recipient mice as indicated (*n* = 5 mice per group).

(D) (G) FACS histograms of the cell cycle marker Ki-67 among CTV^+^ CD4^+^ donor T cells before and after adoptive transfer. An example of identical Ki67 staining before and after transfer is shown. In (A and D-F) one-way Anova with Bonferroni correction for multiple comparison. ***P<0,001; ****P<0,0001. Error bars indicate SEM.

**Figure S6 - DOS_juv_ CD4^+^ T cells are essential for Ag-specific B cell generation**

Representative plots (left) and quantification (right; *n* = 5 mice per group) of Fluad-specific induction of (A) CD19^+^ CD38^-^ GL7^+^ germinal center (GB) B cells and (B) CD19^-/low^, CD45R^-/low^ (B220) CD138^+^ plasma cells in vaccinated old recipients subjected to adoptive transfer. Recipient animals were injected with DOS-juvenated CD4^+^ T cells (DOS_juv_ T cells), their DOS CD4^+^ T cell depleted splenocytes (DOS cells (-CD4^+^)), their aged matched CD4^+^ T cells from old donor animals not exposed to the DOS (CD4^+^ T cells (No DOS)) or young CD4^+^ T cell transfer from naïve animals. Recipient animals were vaccinated with Fluad one day after adoptive transfer. Young, old and DOS treated animals vaccinated without adoptive transfer are also shown, as per Fig. 3. Lymph node cells were studied two weeks after vaccination (day 15).

(A) (C) CXCR5^+^ PD1^+^ Tfh detection among CD45.1+ CD4^+^ T cells subjected to adoptive transfer in old CD45.2 recipients and analysed 15 days after recipient Fluad vaccination. Representative FACS plots (left) and pooled data (right) from (*n* = 5 mice per group) are shown. A background example of vaccinated old transfer-free animals is shown.

(B) (D) Experimental design. CD4^+^ T cells were purified from total splenocytes of unvaccinated old mice (20 months) then transduced with sh-Scramble or triple sh-Sestrins (sh-sestrin1, sh-sestrin2, sh-sestrin3). After 96 hours, transduced CD4^+^ T cells were co-cultured with autologous APCs in the presence or in the absence of Fluad (1:100 of the human dose). After additional 10 days, plasma cells were characterised. Antigen specific plasma cells are derived by subtraction of stimulated samples against unstimulated background controls, with or without DOS.

(C) (E) Antigen-specific plasma cell generation in immune cultures as in (D). Note that sestrins silencing in old CD4^+^ T cells phenocopies DOS action. As such, no further induction of antigen-specific plasma cells can be obtained when DOS is added to cultures with sestrin deficient CD4^+^ T cells (*n* = 5 mice per group). Young B cell cultures were used as control.

In (A-C and E) one-way Anova with Bonferroni post-correction for multiple comparisons. *P<0,05; ***P<0,001; ***P<0,0001. Error bars indicate SEM.

**Figure S7 – Single DOS administration induces long-term sestrin inhibition *in vivo***

(A) Time course of sestrin 2 expression (MFI) *in vivo* between DOS treated and vehicle injected mice. Old mice (*n* = 5 mice per group) were sub-cutaneously injected with DOS, then bled for CD4^+^ T cell sestrin 2 assessment at the indicated time points. Young CD4+ T cells are shown. MFI, mean fluorescence intensity.

(B) Immunoblot analysis of sestrin 2 inhibition (left) and quantification (right) among circulating mouse CD4^+^ T cells treated or not with DOS and assessed 72 hours later. Representative of *n* = 4 animals per group. Data are expressed as fold change to the untreated control (no DOS), set as 1.

(C) Sestrin 2 levels in splenocytes of young, old and DOS-juvenated mice in the presence of Fluad vaccines (10 days post treatment). Quantification of *sestrin 2* expression levels assessed by qPCR. Data are from *n* = 4 mice (young), *n* = 4 mice (old) or *n* = 6 mice (DOS).

(D) Sestrin protein levels six months after systemic DOS treatment. Representative FACS plots and pooled data from (*n* = 5 mice per group) are shown.

(E) Investigation of *sestrin* acetylation (H3K27ac in the *sestrin 2* locus) among splenocytes of old infected animals (20 months) treated or not with DOS and assessed 10 weeks later (*n* = 6 mice per group). Young (3 months) infected animals served as control. The *Sestrin* 2 gene *locus* was analyzed by qPCR after chromatin immunoprecipitation (ChIP) with anti-H3K27ac antibody.

(F) Evaluation of HDAC1 activity. CD4^+^ T cell HDAC1 was immunoprecipitated from lymphoid organs of mice as in (E) and incubated for 30 minutes (30 °C) with exogenous T cell DNA purified from old mice not exposed to DOS (*n =* 5 mice per group). HDAC1 activity (direct deacetylase) was assessed by ELISA assay by detecting H3K27 acetylation levels in target DNA.

In (A and C-F) one-way Anova with Bonferroni post-correction for multiple comparisons. *P<0,05; **P<0,01; ***P<0,001****P<0,0001. Error bars indicate SEM.

**Figure S8 – DOS treatment restores Tfh responses upon vaccination**

(A) Representative FACS plots (left) and cumulative data (right; *n* = 6 donors) of primary Tfh cell responses defined as PD1^+^ CXCR5^+^ CD3^+^ CD4^+^ T cells among splenocytes of old mice 14 days after DOS treatment, with or without Fluad vaccination. Antigen specific responses are denoted by light greying throughout. Young and old control mice with and without Fluad vaccination are shown. Fluad specific Tfh induction is shown (right).

**Figure S9 – DOS_juv_ T cells present functional TCR revisions**

(A) TCR rearrangements measured as TREC quantification from CD4^+^ T cell genomic DNA purified from lymph-nodes of old H1N1 infected mice (20 months) treated or not with DOS (0.1mg/kg). TREC levels were assessed by qPCR. Young infected mice (3 months) served as control.

(B) Quantification (left) of TCRvϕ3 variant usage in CD4^+^ T cells purified from lymph-nodes of young, old and DOS-juvenated mice (as above) assessed by flow-cytometry 2 weeks after infection. Viral specific TCR inductions are shown (right).

(C) TREC quantification in DOS_juv_ CD45.1^+^ CD4^+^ transferred T cells derived from lymph-nodes of old CD45.2 mice subjected to lethal H1N1 infection in the absence of vaccination and assessed 2 weeks later.

(D) Time course of TCR rearrangements assessed by TREC quantification in human CD4^+^ T cells treated with DOS. CD4^+^ T cells were activated with anti-CD3/CD28 and rhIL2 and assessed at indicated time points. Data are from *n* = 12 donors (18 hours; 3 days), or *n* = 9 donors (7 days). Absolute raw data are shown.

(E) Generation of YF-specificity upon DOS treatment. CD4^+^ T cells were pre-treated with or without DOS for 4 hours then conjugated with autologous APCs preloaded with different YF-vaccine dilutions, as indicated. IFNγ production was investigated in the T_STEM_ compartment by flow cytometry 18 hours later (*n* = 6 donors). Note absence of YF responses in T cells not exposed to DOS, regardless of the antigen dosing. Specificity T cell population controls (T_CM_ and T_EM_) did not produce any YF response (top right).

(I) (F) T_STEM_ were derived from YF-vaccine primed APC-T cell conjugates 18 hours after culture, rested for additional 5 days then re-challenged with autologous YF-vaccine loaded APCs for 18 hours. IFNγ release was quantified by ELISA in the culture supernatants (*n* = 4 donors). Specificity T cell population controls (T_CM_ and T_EM_) are shown.

(II) (G) Example of human T_STEM_ gating strategy for YF peptide stimulation with or without DOS (note TCF1 staining confirming 100% stem like origin) Individual gatings and positive (PMA-ionomycin induced) IFNγ induction controls are shown. Yellow shadows, YF specific interval response.

(H, top) quantification of IFNγ production among the same stem-like human T cells. Autologous APCs and T cells were pre-treated with peptides or DOS respectively for 4 hours prior to co-culture. Yellow fever specific IFNγ producing stem cells are derived by subtraction against unstimulated backgrounds, with or without DOS (H, bottom). No response could be detected using a negative peptide control (actin; 2μg/mL), confirming induction of YF specific responses. Absolute IFNγ producing T_STEM_ counts are shown (*n* = 5 donors).

(I) Generation of YF-specific responses, transgenic T cell controls. OT-II T cells were conjugated with congenic YF pulsed APCs for 18 hours followed by stem-like CD4^+^ T cell sorting. Five days later, purified stem-like T cells were restimulated as above and YF-specific IFNγ release was quantified among culture supernatants by ELISA (top). In control experiments, OT-II T cells were derived from animals that had been immunised with ovalbumin, 15 days after vaccination. The recall response to OVA was used as a control to quantify antigen-specific IFNγ release against the transgenic T cell antigen (OVA; bottom). OVA, ovalbumin. *n* = 5 mice per group throughout.

In (A-C and H-I) One-way Anova with Bonferroni correction for multiple comparisons. In (D) F test. In (E-F) two tailed paired T test was used. *P<0,05; **P<0,01; ***P<0,001; ****P<0,0001. Error bars indicate SEM.

**Figure S10 – DOS-driven YF-specific TCR clonotypes**

(A) Yellow Fever (YF) induced TCRs table list referred to Donor A in Fig.5.

(B) Single TCR clonotypes among different conditions in the Donor D

**Figure S11 – DOS juvenation restores Rag function in peripheral T cells**

(A) Representative immunoblots (left) and quantification (right) of Rag protein expression from human CD4^+^ T cells treated or not with DOS and activated with anti-CD3/anti-CD28 for 18 hours are shown (*n =* 3 donors). Data are expressed as fold change to the untreated control (no DOS), set as 1.

(B) Rag 2 proteins were immunoprecipitated from the chromatin of human CD4^+^ T cells treated as in (A) then followed by ELISA with anti-Rag antibodies. Data are expressed as fold change to the untreated control (no DOS), set as 1.

(C) Chromatin immunoprecipitation of Rag1 protein from T cells derived upon dissolution of immune-conjugates with autologous YF-pulsed APCs, with or without DOS. YF-specific TCR recombination factor binding to T cell DNA was calculated subtracting endogenous Rag factors bound to T cells forming conjugates in the absence of antigens (*n* = 3 donors).

(D) Representative FACS plot validation (left) and quantification (right; *n* = 3 donors) of Rag knock-down in human CD4^+^ T cells. HEK293 cells were used as negative Rag expression control (bottom left). Note that Rag silencing reduces Rag expression to background fluorescence of HEK293 cells naturally lacking Rag proteins, validating known down efficiency.

(E) Rag deficiency abrogates DOS-driven NGS identified TCR revisions. Human CD4^+^ T cells were transfected with siCtrl or siRAG1/siRAG2 for 3 days. The T cells were then treated with or without DOS for 4 hours, prior to culture with YF-pulsed or antigen-free APCs for additional 18 hours. The YF specific DOS-juvenated TCRs derived by background subtraction with no YF peptides are shown (*n* = 4 donors).

(F) Effect of Rag protein silencing on the YF specific TCR rearrangement, assessed by TREC quantification (*n* = 4 donors).

In (A-C and E) two tailed paired T test was used. In (D) One-way Anova with Bonferroni correction for multiple comparisons. In **f,** F test. *P<0,05; **P<0,01; ***P<0,001; ****P<0,0001. Error bars indicate SEM.

## Supporting information

Graphical abstract

## Materials and Methods

**Table 1.** List of unconjugated antibodies.

| Antibody | Supplier | Clone | Catalogue number |
| --- | --- | --- | --- |
| p-Ampk α-Rabbit mAb | Cell Signaling | 40H9 | 2535S |
| P-p38 MAPK - Rabbit mAb | Cell Signaling | D3F9 | 4511S |
| P-SAPK/JNK Rabbit mAb | Cell Signaling | 81E11 | 4668S |
| SESN1 Rabbit pAb | Gene Tex | / | GTX118141 |
| SESN2 Rabbit pAb | Gene Tex | / | GTX116925 |
| SESN3 Rabbit pAb | Gene Tex | / | GTX112277 |
| Histone H3 Rabbit pAb | GIANNI Srl | / | AMab1791 |
| Anti-rabbit IgG, HRP-linked Antibody | Cell Signaling | / | 7074S |
| WDR59 Rabbit mAb | Cell Signaling | D4Z7A | 53385S |
| MIOS Rabbit mAb | Cell Signaling | D12C6 | 13557S |
| p-mTOR Rabbit mAb | Cell Signaling | D9C2 | 5536S |
| JNK Rabbit pAb | Cell Signaling | / | 9252T |
| H2B Rabbit pAb | Cell Signaling | V119 | 8135 |
| AMPK total pAb | Cell Signaling | / | 2532S |
| P38 total mAb | Cell Signaling | D13E1 | 8690S |
| ERK total mAb | Cell Signaling | 137F5 | 4695T |
| Anti-FLAG pAb | GeneTex | / | GTX115043 |
| Anti-Myc mAb | GeneTex | / | GTX29106 |
| p-ERK Rabbit mAb | Cell Signaling | D13.14.4e | 4370S |
| GAPDH Rabbit mAb | Cell Signaling | D16h11 | 5174S |
| anti-ubiquitinated<br>proteins mAb | EMD Millipore | FK2 | 04-263 |
| Rag1 Rabbit mAb | Cell Signaling | D36B3 | 3968S |
| Rag2 Rabbit pAb | Invitrogen | XJ3737974B | PA5-88580 |
| α-CD3 Ab | Invitrogen | OKT3 | 14-0037-82 |
| Acetyl-Histone H3<br>Rabbit mAb | Cell Signaling | D5EA | 8173 |
| 5-Methylcytosine<br>Rabbit mAb | Cell Signaling | D3S2Z | 28692 |
| HDAC1 Rabbit mAb | Cell Signaling | D5C6U | 34589 |
| $\beta$ -Galactosidase | Cell Signaling | E2U2I | 27198 |
| P-16 | Cell Signaling | E6N8P | 18769 |
| $\beta$ -Catenin | Cell Signaling | D10A8 | 8480 |

**Table 2.**
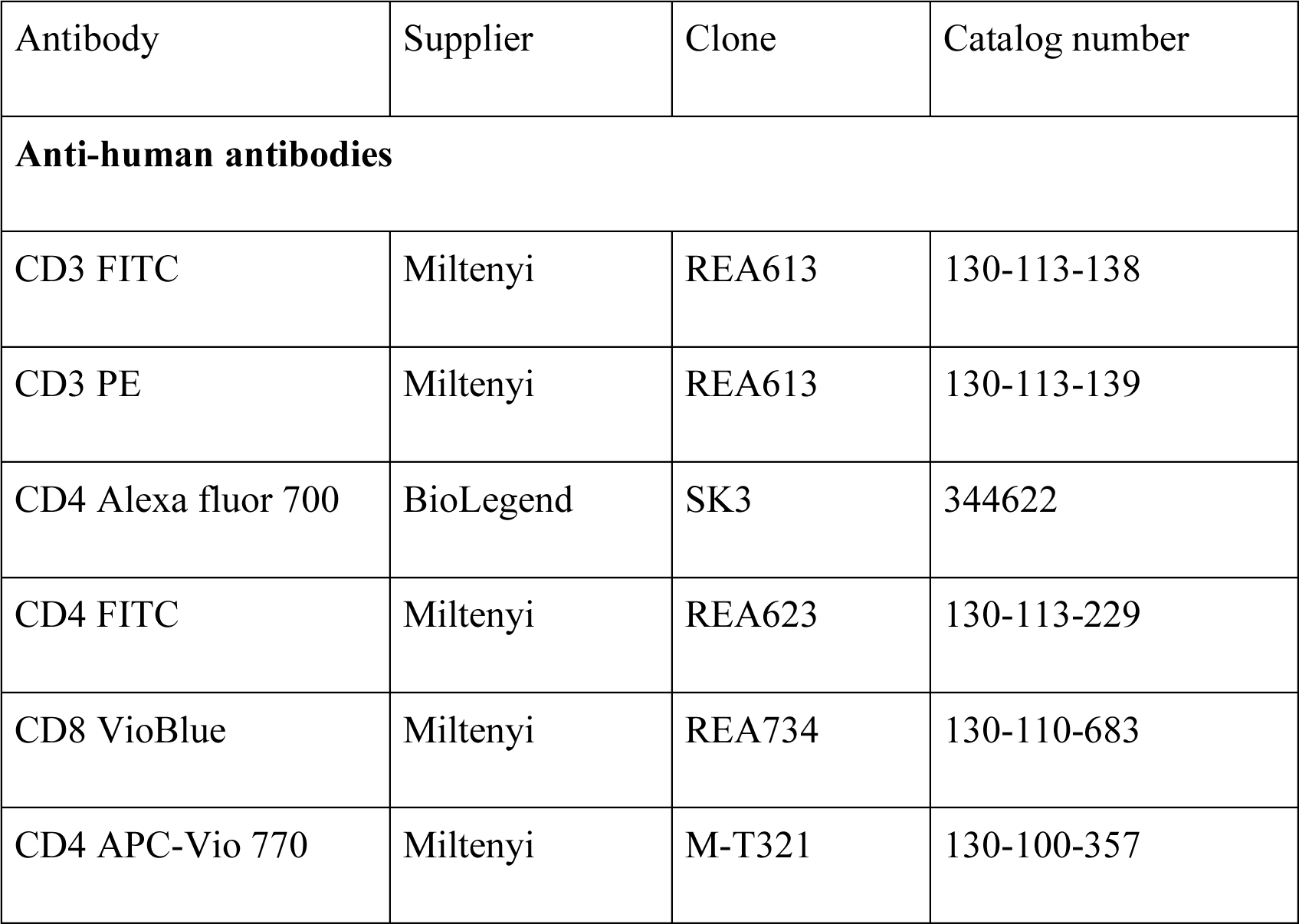
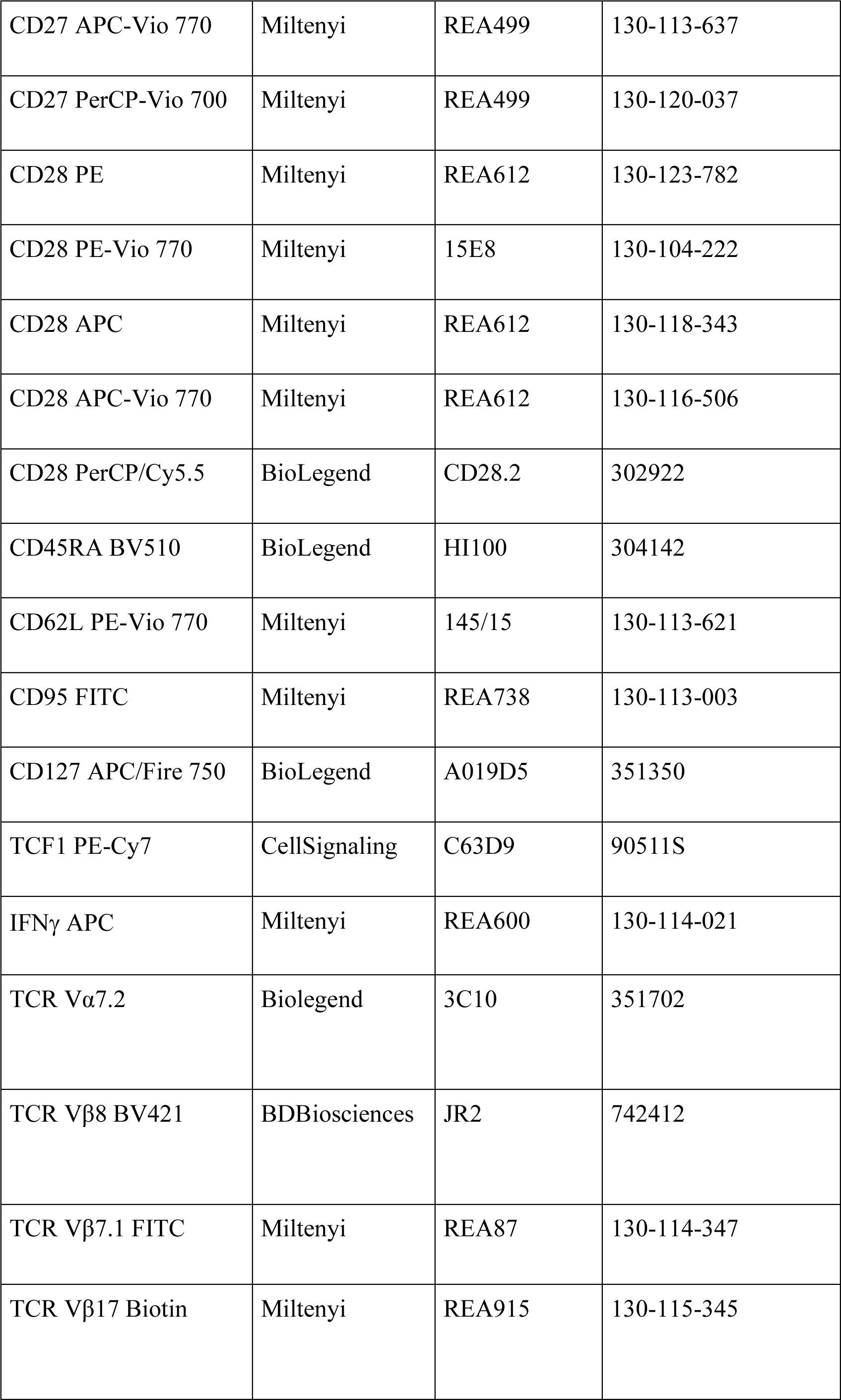
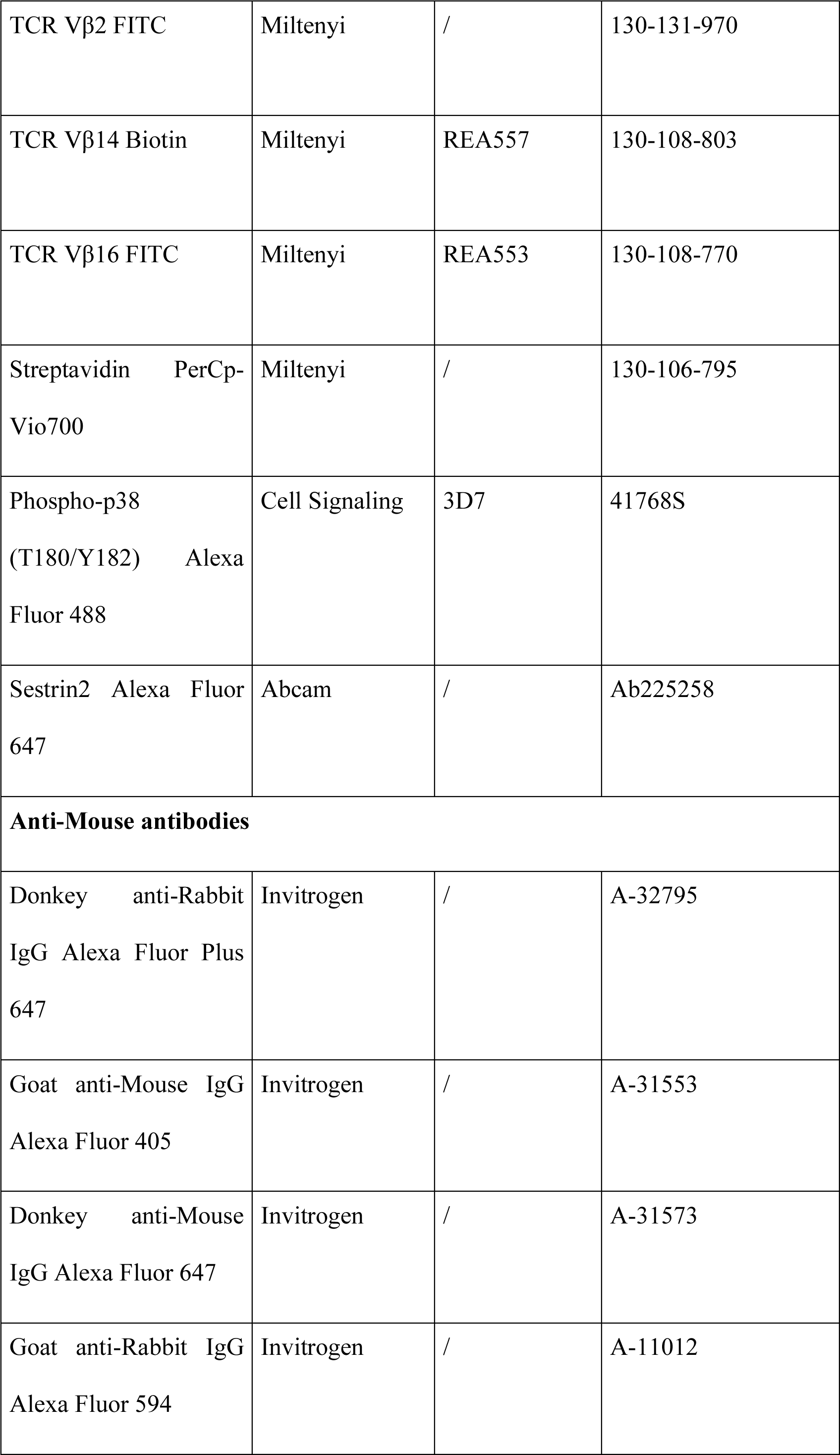
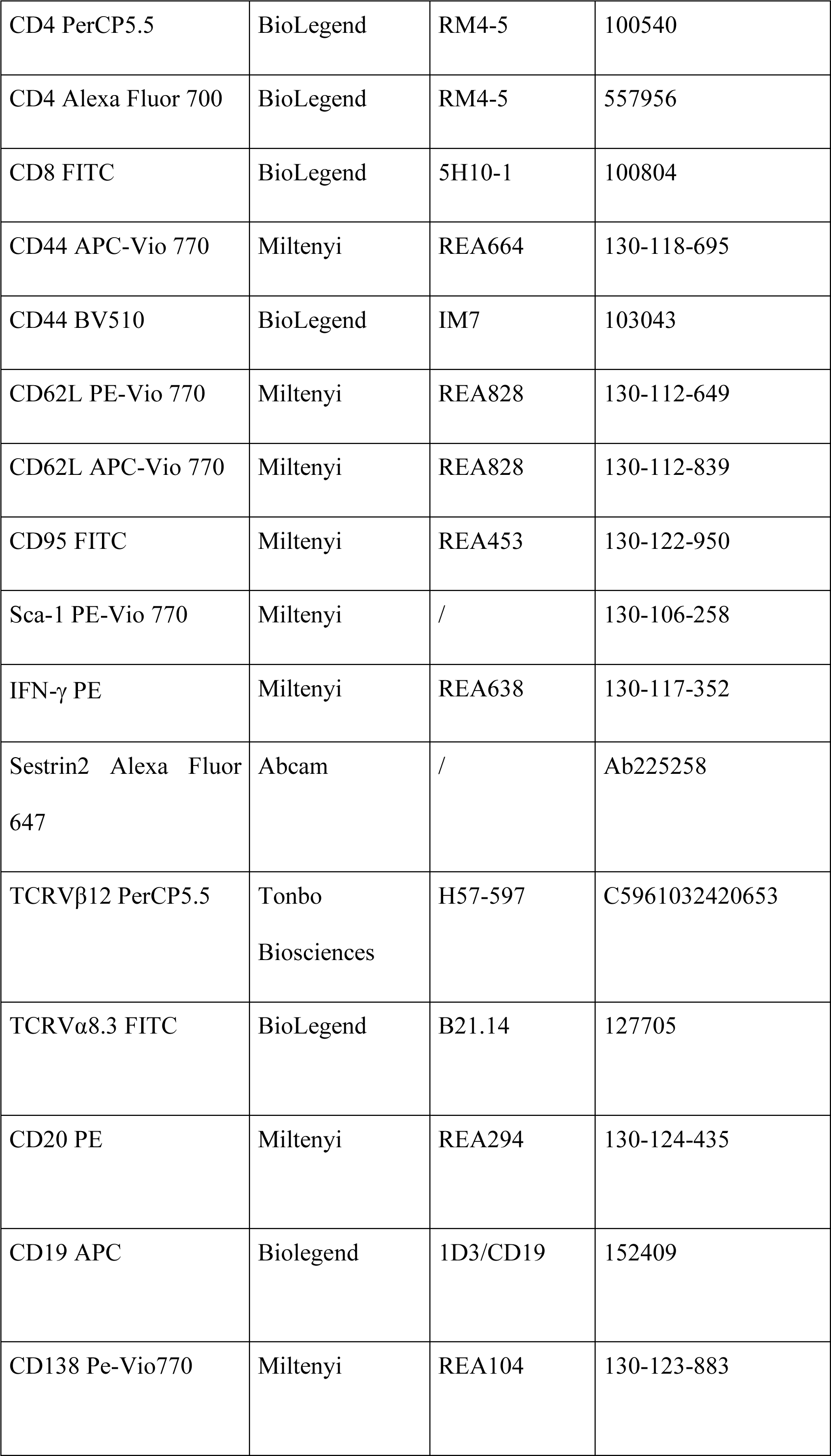
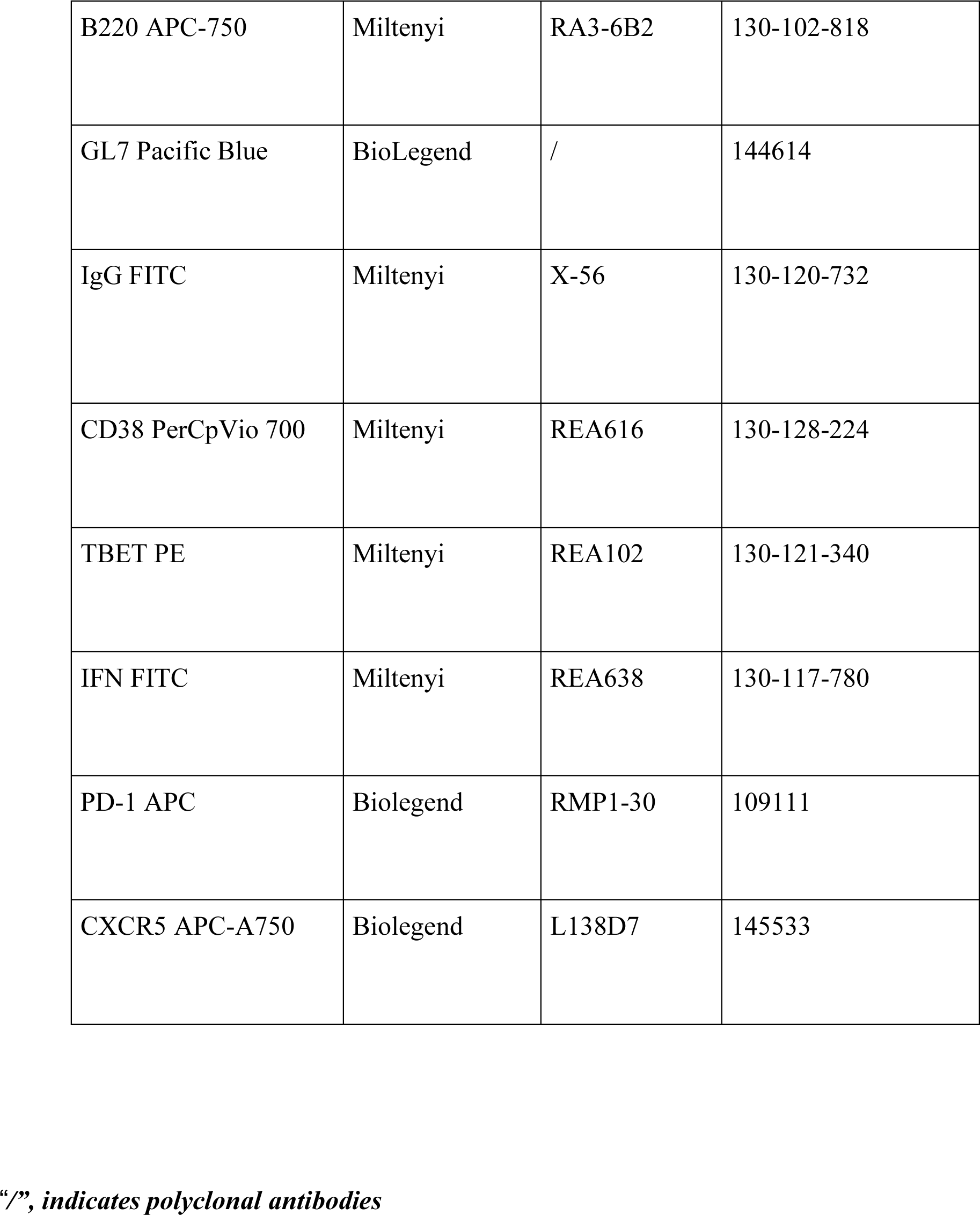
List of fluorescent antibodies.

### Human and mouse studies

All human specimens were obtained from de-anonymized volunteers unless otherwise specified with the approval of Fondazione Santa Lucia Ethical Committee and voluntary informed consent in accordance with the Declaration of Helsinki in all cases. No compensation was offered. For mouse studies, we received approval from the Italian Ministry of Health. Mouse husbandry was controlled by standard circadian rhythms (12 hours of dark and 12 hours of light; fixed temperature 25°C, humidity between 40-60%). For ageing studies, old donors were 65-80 years old.

### DOS synthesis and formulation

The synthesis of c-DOS46L consisted of four stages. In stage 1 the naked linear peptide, CDOS1, was generated. Specifically, the first amino acid (Fmoc-Val-OH) was loaded on the 2-CTC resin and then the Fmoc was deprotected using Piperidine in DMF. The remaining 4 amino acids were coupled one by one in predetermined sequence using HOBt and DIC. Followed deprotection of Fmoc group using Piperidine in DMF, after each amino acid protection to avail for next amino acid coupling to elongate the chain. In the end of this step the crude linear peptide was obtained with a charge of 2%TFA /DCM to knock off the resin.

Stage 2 was aimed to obtain the crude cyclic peptide, CDOS2. This was obtained by cyclization of the linear peptide by using EDC. HCl/HOBt.H_2_O/ DIPEA in DMF as a solvent.

In stage 3 the CDOS2 was charged on the Reverse phase column to yield the pure cyclic peptide, CDOS3. In the end, in stage 4, the column pooled fractions were distilled to minimum desired level and then filtered, the material was then dried at 55’5°C under vacuum to obtain the targeted pure cyclic peptide, CDOS4. CDOS4 means the C-DOS46L used in all the paper.

Other DOS compounds were synthetized using the same procedure, NH_2_COOH cyclisation and FMOC deprotection.

The DOS used in the paper was resuspended in 100% of DMSO (Sigma); subsequential dilution were done in PBS. For pharmacokinetic (PK) studies, DOS was instead resuspended in a solution of 80% PG (propylene glycol) and 20% PEG (Polyethylenglycolum 400). In all the experiments, DOS refers to 10 µM of cyclic DOS46L unless otherwise specified.

### DOS peptide library in vitro screening by inhibition of AMPK1

The screening was performed using the kit ADP-Glo kinase Assay (V6930; Promega). Specifically, 250 ng of the recombinant AMPK1 (P47-110GH-10; Signal Chem) were pre-incubated with 10 µM of DOS peptide, before the addition of the pre-incubated complex containing 1 µg of sestrin1 (GWB-5DFF50; Genway) or sestrin2 (GWB-66FAFC; Genway), ATP (200 µM; V915B; Promega), AMP (200 µM; V506B; Promega) and SAMsite (0,2 µg/µl; S07-58-1MG; Signal Chem). In addition, in some experiments testing the effect of DOS on the AMPK activity we added to the reaction the AMP-activated protein kinase activator A-769662 (100 µM; ab120335), as indicated.

AMPK activity was then detected by adding 40 µl of kinase detection reagent according to manufacturing protocol. Luminescence was acquired at the Varioskan reader (ThermoFisher). Background AMPK activity without sestrin was subtracted.

### Cell isolation and culture

Peripheral blood mononuclear cells (PBMCs) were isolated by Ficoll (17-1440-03; GE Healthcare) density gradient centrifugation from blood of healthy volunteers.

Primary human CD4^+^ T cells were isolated using CD4 MicroBeads (130-045-101; Miltenyi); primary human senescent CD4^+^ CD27^-^ CD28^-^ T cells (thereafter T_sen_) were isolated using CD4^+^ T Cell Isolation Kit (130-096-533; Miltenyi) followed by depletion of CD27^+^ and CD28^+^ T cells using CD27 MicroBeads (130-051-601; Miltenyi) and CD28 MicroBead Kit (130-093-247; Miltenyi). CD3^-^ cells (thereafter APCs) were isolated by depleting CD3^+^ cells using human CD3 MicroBeads (130-050-101; Miltenyi). The CD27^+^ expressing population of human CD4^+^ T cells that co-expresses also CD28^1, 2^ was used as early differentiated (non-senescent) T cell (T_erl_) control.

For isolation of mouse T cells, CD4 (L3T4) MicroBeads (130-117-043; Miltenyi) were used.

All the isolations were conducted following the manufacturer#s protocols.

T_erl_ cells were cultured in RPMI1640 (R2405; Sigma-Aldrich) supplemented with 10% FBS (F9665; Sigma-Aldrich) and 1% Pen-Strep (DE17-602E; Lonza). Dynabeads Human T-Activator CD3/CD28 (11161D; Gibco; 1:100) or plate bound with anti-CD3 and anti-CD28 antibodies (0.5µg/ml each; Invitrogen and Miltenyi) were used to activate these T cells as indicated by the manufacturer#s protocol. For primary human highly differentiated senescent T_sen_ cell cultures, plate bound CD3 and recombinant human IL2 (5ng/mL) were used instead as previously described^1,2^. For primary cultures, all T cells were incubated in RPMI 1640 media (R8758-500ml; Sigma-Aldrich) supplemented with 10% FBS (fetal bovine) in an incubator (37 $C and 5% of Co2).

### Western blotting protocol

Analysed samples were lysed with RIPA buffer plus phosphatase and protease inhibitors. Protein lysates were quantified by Lowry protocol and then denatured with Lemmli buffer + β-Mercaptoethanol. 7.5 µg of proteins were separated on polyacrylamide gel and then transferred on nitrocellulose membrane by the Trans-Blot dry system (Biorad). Membranes were blocked on milk (5%) or BSA (5%) depending on if the protein to detect were phosphorylated or not.

A comprehensive list of primary and secondary antibodies used for western blotting is reported in table S1. Primary antibodies were used 1:1000 dilution unless otherwise specified. For secondary antibodies 1:5000 dilution was used.

### Immunophenotypic analysis

All extracellular stainings were performed resuspending cells in the necessary antibody mix (1:100) in PBS with 0.5% BSA and 2mM EDTA for 30 minutes at 4°C in the dark. If intracellular staining was necessary, cells were then fixed with BD Cytofix Fixation Buffer (554655; BD Bioscience) for 15 minutes at 37°C and permeabilized with BD Phosflow Perm Buffer III (558050; BD Bioscience) for 30 minutes at -20°C. Cells were then stained with either primary conjugated antibody (30 minutes at 4°C) or unconjugated primary antibody (1 hour at 4°C in the dark) followed by a secondary conjugated antibody diluted 1: 500 (30 minutes at 4°C in the dark). Cells were analysed on a CytoFLEX flow cytometer (Beckman Coulter). Dead cells were excluded. A comprehensive list of antibodies used for flow cytometry is reported in Table 2.

### Confocal imaging

To investigate intracellular localization of DOS, 1*10^6^ of purified CD4^+^ T cells were activated with anti-CD3/CD28 and treated with 10 µM of DOS-FITC for 4 hours. Cells were then collected and plated in coverslips pre-treated with poly-lysine and fixed with formaldehyde 4%. Cells were then stained with the primary antibody anti-CD3 (1:100; 14-0037-82; eBioscience) and with a secondary anti-mouse antibody conjugated with AF-647 (1:250; Invitrogen).

To check the co-localization of DOS with the ER, 1*10^6^ of purified human CD4^+^ T cells were treated with DOS-FITC for 4 hours and then stained with primary antibody anti-KDEL (1:100; ab12223; Abcam) and anti-sestrin 2 (1:100; Gentex). Secondary antibodies used were anti-mouse conjugated-AF647 (1:250; Invitrogen) or anti-rabbit conjugated-AF594 (1:250; Invitrogen).

To evaluate the impact of DOS on DNA degradation, T_sen_ were purified as above described and then treated or not for one week with DOS. 5*10^5^ cells were then plated in a poly-lysine pre-coated coverslip and surface-stained with the anti-CD4 antibody APC-Vio 770 (1:100; Miltenyi). Cells were then fixed with 4% formaldehyde and permeabilized with 0,25% Triton (T8787; Sigma). Cells were firstly intracellularly stained with primary rabbit anti-human H2AX antibody (1:100; 2595S; Cell Signaling); and then with secondary anti-rabbit conjugated-AF488 antibody (1:250; ThermoFisher). For all imaging experiments, sections were mounted in a glass slide with ProLong gold antifade mountant with DAPI (P36931; Invitrogen) and fluorescence images were acquired on an Axio Observer confocal microscope (Zeiss).

### Analysis of DOS incorporation

PBMCs were treated with 10 µM DOS-FITC conjugate for 30 minutes, washed, fixed and then immediately analysed by flow cytometry. Surface staining to CD4, CD8, CD27 and CD28, CD62L, KLRG1 receptors and sestrin2 and p-p38 (sMAC) intracellular staining allowed investigation of incorporation of DOS in the different T cell subsets.

### Inhibitor concentration 50 (IC50)

To study the IC50 of DOS toward sestrin driven AMPK activity, a mix containing DOS peptides (in 5-fold dilutions between 50 µM and 1 pM), 5 ng/µl recombinant sestrin 2, 50 µM DTT, 0.2 µg/µl SAMsite, 200 µM ATP and 200 µM AMP and 5 ng/µl recombinant AMPK was diluted in kinase buffer as per initial *in vitro* DOS screening. Reaction volume was 25µl coating a 374 well plates. The mix was incubated for 1 hour at room temperature and the inhibition of sestrin driven AMPK activity was measured following the ADP-Glo kinase Assay kit. Data were analyzed following the manufacturing information of the kit; specifically, the IC50 value was extrapolate via Sigmoidal, 4PL, X is log(concentration) non-linear regression analysis model-GraphPad Prism software.

### Sestrin-driven AMPK activity assay by radiolabeled ATP

Commercial recombinant AMPK protein (P47-110GH-10; Signal Chem) was used in combination with recombinant sestrin 2 protein (GWB-66FAFC; Genway). The reaction was performed with 200μM AMP, 200μM ATP (0.5 μCi 32P-ATP per reaction) 50μM SAMS, 250 ng AMPK and 1ug of Sestrin2 in the presence of a kinase buffer (40 mM EDTA, 5mM MgCl2, 0,025% BSA, 0,8mM DTT). After 30 minutes of incubation at 30°C, the reaction was stopped by adding 80μl of H3PO4 1%. Reaction mixture was then transferred to 96 well Multiscreen plate (MAPHNOB50; Millipore) followed by detection in a TopCount® NXTTM.

### KD titration curve

Recombinant Sestrin2 was diluted in TBS, pH 7.4, 2 mM MgCl_2_ at a final concentration of 5 µg/ml and used to coat 96 well plates overnight at 4°C. Coated plates were washed twice with TBS to remove residual soluble proteins and blocked in TBS 1% BSA solution for 1hour at room temperature. A serial DOS-biotin dilutions were prepared in duplicates in TBS 1% BSA and incubated in Sestrin2 coated plates for 90 minutes at room temperature followed by washing steps to remove unbound DOS. The binding between DOS-biotin and Sestrin2 was fixed by using 2,5% of Glutaraldehyde in HBS (50 mM HEPES/NaOH, pH 7,4; 150 mM NaCl). Sestrin2-bound DOS was quantified by ELISA. Primary antibody (streptavidin-HRP 1:2000) was incubated in 50 µl of TBS 1% BSA 90 minutes at room temperature. To detect antibody, 50 µl of TMB substrate (N301; ThermoFisher) was added to each well, incubated for 30 minutes, and reaction was stopped with stop solution (N600; ThermoFisher). The optical density (OD450) was measured by an ELISA on a micro-plate reader (AMR-100, Biolab). KD was calculated in GraphPad Prism by performing a !Non-linear regression statistical analysis” with the following equation^3^:

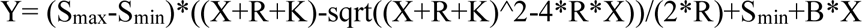

Where:

Y being the signal value S

X being the concentration of added DOS

R being the concentration of immobilized protein

K being the dissociation constant

B being the background slope

### *In vitro* pull down with biotinylated-DOS

0.1 µg/µl of recombinant tagged proteins (sestrin1-FLAG-tagged; sestrin2-FLAG-tagged; AMPK1-His-tagged) were incubated with 10 µM of DOS-biotin overnight at 4°C. Streptavidin-agarose beads (20353; Thermo Scientific) were washed with PD buffer (20mM Hepes pH 7.5, 100mM NaCl, 10mM KCl, 1mM MgCl_2_ 50 µM DTT, 10% glycerol and 0,1% BSA) and blocked overnight with 5% BSA before incubation with reaction substrates (recombinant tagged proteins plus biotinylated-DOS) for 1 hour at room temperature. Streptavidin beads combined with recombinant sestrins were then washed with PD buffer and proteins were eluted by addition of Laemmli buffer 1x. Followed western-blot assay, proteins were detected with anti-FLAG and anti-Myc antibodies. In parallel, an *in vitro* pull down with DOS-biotinylated was performed directly on CD4^+^ T cell lysates and presence of Sestrins, p38, JNK, ERK, MIOS and WDR59 was assayed by western-blot.

### *In vitro* Kinase Assay

Purified human senescent CD4^+^ T cells were lysed in IP buffer (a solution of Hepes (20 mM), NaCl (100 mM), KCl (100 mM), MgCl_2_ (1 mM), Glycerol (10%) and NP-40 (0.5%) and protease inhibitor (1X)) for 30 minutes on ice followed by centrifugation at 20000g for 30 minutes. The supernatant was then incubated with anti-Sestrin 2 antibody overnight (5 µg). The following day, protein A/G agarose beads (20422; Thermo Scientific) were incubated together with the lysate with anti-sesn2 antibody for 3 hours at 4°C on a rotor.

For analysis of the ability of DOS to inhibit MAPK, the immunoprecipitates were treated or not with DOS and ATP (200μM; 9804; Cell Signaling) for 30 minutes at 30°C. The resulting kinase reaction product was then coated in an ELISA plate at 4°C overnight. The plate was then blocked with 5% milk in PBS-Tween 0.5% for 1 hour. To assess the MAPK activity, the plate was incubated for 2 hours at room temperature with one of the following antibodies: anti-p-p38 (1:100), anti-phospho-SAPK/JNK (1:100) and phospho-ERK 1/2 (1:100). The secondary anti-rabbit IgG-HRP conjugated antibody (1:500) was incubated for 1 hour at room temperature. Autophosphorylation within each individual MAPK kinase in the sMAC was derived by subtraction of the corresponding *in vitro* reaction in the absence of ATP.

To prove that DOS action is AMPK dependent, the immunoprecipitates were derived from senescent T cells that were depleted or not of AMPK by lentiviral vectors. Ninety-six hours after transduction, T cell extracts were subjected to immunoprecipitation with anti-sestrin antibodies (sestrin 2), then treated with or without DOS and ATP (200 μM) for 30 minutes at 30°C. sMAC activity was assessed with anti-p-p38 antibodies by ELISA after overnight coating on a 96 well plate at 4°C. Alternatively, Compound C (10 µM; 171260; Sigma Aldrich) was added directly to the kinase reaction or to the live cultures of senescent T cells for 2 hours in the presence or in the absence of DOS prior to identical ELISA assays. After overnight blocking (as above), plates were incubated with the primary anti-p-p38 antibody (1:100) for 1 hour at room temperature in the dark, followed by incubation with secondary antibody (1:500) 1 hour in the same conditions. For both the assays, TMB substrate was used for detection. Absorbance was detected at 450 nm on a micro-plate ELISA reader (AMR-100, Biolab).

### *In vitro* sMAC disruption

To study sMAC disruption after DOS treatment, AMPK immunoprecipitates obtained as described above were coated on an ELISA plate at 4°C overnight. The plate (wash-in) was then treated with DOS and ATP (200 μM) for 2 hours at 30°C. After the reaction, the wash-out was used to derive wash-out plates by identical overnight coating at 4°C. Wash-in plates and wash-out plates were incubated with anti-sestrin1, anti-sestrin2 or anti-sestrin3 antibodies (1:100) as primary antibodies and detected by ELISA reader following secondary antibody detection. The same experiment was conducted to study sMAC disruption after AMPK inhibition with Compound C; in this experiment the plate was treated with or without Compound C (10 µM) for 2 hours at 30°C, followed by AMPK detection.

### Analysis of sMAC degradation

For the analysis of sestrin2 ubiquitination upon DOS treatment, 5 x10^6^ senescent CD4^+^ T cells were lysed with 100μL of denaturing lysis buffer (a water solution containing SDS (1%), EDTA (5 mM) and β-mercaptoethanol (10 mM)) and boiled for 5 minutes. Samples were then diluted with 900 µl of non-denaturing washing buffer. After lysis, cell extracts were incubated with anti-Sestrin2 antibody (5 µg) at 4°C overnight. In parallel, 200 µl of protein A/G beads were washed 4-5 times in IP buffer and blocked in IP buffer overnight at 4°C. The day after, blocked protein A/G beads were added to the sample and incubated for 3h at 4°C. Protein-beads conjugates were washed with IP buffer and then boiled in presence of Laemmli buffer. Samples were then loaded into polyacrylamide gel and analysed with antibodies to ubiquitinated proteins. For analysis of proteasomal degradation of sMAC, purified human T_sen_ were pre-treated with or without the proteasomal inhibitor MG132 (10 µM; M7449; Sigma-Aldrich) for 30 minutes at 37°C, prior to DOS exposure for 2 hours in culture. Cells were then lysed as described above and analysed by direct immunoblotting for sestrin2 and AMPK.

### AMPKγ-ATP loading assay

Recombinant AMPK γ1 chain (AMPKɣ; AR51158PU-S, OriGene) and Sestrin 2 (GWB-66FAFC-50UG, Aviva) were diluted in IP buffer (Hepes 20 mM, NaCl 100 mM, KCl 100 mM, MgCl_2_ 1 mM, 10% Glycerol) at a final concentration of 5 µg/ml and 20µg/ml respectively. ATP (200 µM) was used in the reaction. For IP loading, AMPKɣ was immunoprecipitated with Anti-His tag monoclonal antibody (TA100013, OriGene) after 5 minutes of *in vitro* reaction at 30 °C. Reaction volume was 50μl.

In order to follow in the window of the ATP detection, the samples were serial diluted with dilution buffer in the range 10^-6^ to 10^-12^ M. ATP was detected using the ATP Bioluminescence Assay Kit (HS II-11 699 709 001-Roche) following the manufacturer’s instructions. Luminescence was acquired at the Varioskan LUX Multimode Microplate reader (ThermoFisher).

### Fluorescent ATP-binding detection

5µM of recombinant total AMPK (P47-110GH-10; Signal Chem) was mixed with 20µM of recombinant sestrin 2 (Genetex) according to the stoichiometry ratio 1:4. In parallel, recombinant AMPKγ protein (5µM; AR51158PU-S, OriGene) was mixed with 20µM of sestrin (Genetex). Condition without sestrin was used as control. The binding reaction was carried out in 50 µl of Tris-HCl buffer (10mM; PH8); incubated at 30$C for 5 minutes in presence or in absence of DOS. After this time, 10µM of MANT-ATP (NU-202S; Jena Bioscience) was added to each reaction and the released fluorescence was quickly registered at 448nm. Fluorescence signal is proportional to the amount of ATP-bound at the AMPK CBS sites (1 and 3). Addition of 200µM of ATP post MANT-ATP binding disrupted MANT-ATP fluorescence confirming ATP binding.

### Sestrin-AMPK binding strength assay

Human T_erl_ cells were treated with A-769662 (AMP-activated protein kinase activator, ab120335) at 100 µM and incubated for 3 days.

After this time, cells were lysed and total AMPK protein was immunoprecipitated with the anti-AMPKα antibody (2532S, Cell Signaling).

Fifty ng/well of the T cell immunoprecipitates were coated into an ELISA plate and incubated at 4 °C overnight. The plates were then blocked with 4% milk in PBS-Tween 0.5% for 1hour before adding recombinant Sestrin 2 (50 ng/µl) for 2 hours at 30°C.

After this time, urea (3M; 57-13-6, Merck Millipore) was added for 10 minutes to break the AMPK-Sestrin2 binding followed by incubation with the anti-Sestrin2 antibody (1:100, #8487, Cell Signaling) and secondary anti-rabbit IgG-HRP conjugated antibody (1:500). TMB substrate was used for detection. Absorbance was detected at 450 nm on a micro-plate ELISA reader (AMR-100, Biolab). In some experiments, the immunoprecitates were incubated with exogenous sestrin, washed and then analysed for binding strength with or without Urea at various concentration (3M or 6M), to monitor interaction strength.

### Molecular docking simulations

To predict the potential binding site(s) of DOS46L to Sestrin-2, protein-peptide molecular docking was performed using AutoDock Vina 1,2. As the available Sestrin-2 crystal structures in Protein Data Bank do not have a complete defined tertiary structure (PDB code: 5CUF; 5DJ4; 5T0N; 6N0M), structure prediction of Sestrin-2 was fetched from AlphaFold Protein Structure 3 Database. Sestrin-2 structure was then optimized for docking, using Dock Prep tools by UCSF Chimera 4. Briefly in this step, hydrogen atoms were assigned and gasteiger charges and net charges were calculated. Processed Sestrin-2 structure was saved in a PDB file. The 3D-structure of DOS 46L was then generated by CycloPs 5, a GUI python program for building cyclic peptide structures from peptides sequences. Energy minimization and optimization for docking were performed in UCSF Chimera, similarly to Sestrin-2 optimization/following the steps of Sestrin-2 optimization. Optimized structure of DOS 46L was saved in a Sybyl Mol2 file for docking. After protein and ligand preparation, AutoDock Vina with default parameters was used. Models of the complexes of DOS46L interacting with Sestrin-2 were obtained and UCSF Chimera was used to plot the modelling figures.

### Generation of DOS disruption sestrin mutants

Sestrin-α (436T-A; 306P-A) and β (94W-A; 74G-A) mutants and wild type constructions were cloned into pDUAL-PuroR, pDUAL-BlastR and pDUAL-PuroR vectors under the transcriptional control of the spleen focus-forming virus (SFFV) promoter by standard cloning techniques.

GFP, Puromycin (PuroR) and Blasticidin (BastR) resistance genes were expressed under the transcriptional control of the ubiquitin promoter WPRE and CPPT sequences present in the pDUAL vector increase transgene expression system.

Standard cloning techniques were followed. Briefly, sestrin-α and β mutants were first amplified on KanR cloning plasmids, then the plasmid was digested by BamHI-NotI 10U/μL (Thermo Fisher Scientific) restriction enzymes in Fast Digest Buffer for 1hour at 37°C to purify the transgene of interest. Wild type sestrin construction was generated inserting the AatII-NotI fragment of the β construct into the α construct to eliminate mutations. AatII-NotI (10U/ul) enzymatic digestion was conducted in Fast Digest Buffer for 1hour at 37°C to purify the transgene of interest. Fragments were purified by Qiagen cloning standardized protocol and ligated to the pDUAL of interest by the T4 ligase enzyme (Promega) for 1 to 24 hours at room temperature. Ligations were transformed into Echerichia coli XL/1 Blue competent bacteria and the DNA of interest was purified by miniprep. Then, the positive minipreps were transformed into Echerichia coli XL/1 Blue competent bacteria the DNA of interest was purified by midiprep. Qiagen cloning standardized protocols were followed. DNA concentration was measured by Nanodrop spectrophotometer. The size of the purified plasmid was analised by electrophoresis in 1.5% agarose gel (1.5% agarose, RedSafe™ Nucleic Acid Staining Solution (20,000x) (iNtRON) in the TAE 1X buffer). To verify the correct structure of the plasmids, the purified DNA was digested with the restriction enzymes BamHI-NotI 10U/μL (Thermo Fisher Scientific), or with HindIII in Fast Digest Buffer (Thermo Fisher Scientific) for 1 or 2 hours respectively, at 37°C. The product of digestion was run in 1.5% agarose gel. It was verified that the sizes of bands obtained were as expected. All insert sequences were confirmed by Sanger Sequencing.

### Lentivirus production and purification

At day 0, HEK293 T cells (CRL-11268; ATCC) were plated in T75 flasks. The day after, cells (70% confluent) were transfected with 10μl Lipofectamine LTX (15338100; ThermoFisher) combined with PLUS reagent in a ratio 1:1 and 4μg of DNA with the following three plasmids: pCMV-VSV-G envelope vector (RV-110; Cell Biolabs); p.891 packaging vector; p-SIREN-scramble/p-SIREN-shsesn1/ p-SIREN-shsesn2/ p-SIREN-shsesn3 (7), 48 and 72 hours later, supernatants from transfected HEK293 T cells containing lentivirus were collected. Lentiviral particles were then concentrated according to Lenti-Pac Lentivirus Concentration Solution protocol (LT007; GeneCopoeia).

### Cell trace violet (CTV) staining in sestrin null senescent T cells

One million of purified human CD4^+^ T_sen_ was activated with an anti-CD3 antibody coated-plate (0.5μg/ml) plus rhIL-2 (5 ng/ml; 11 011 456 001; Sigma). Forty-eight hours post-activation, cells were transduced with p-SIREN-sh lentivirus particles to target sestrins or with the control p-SIREN scramble at a multiplicity of infection (MOI) of 10. Ninety-six hours after transduction, cells were treated with 5 µM of Cell Trace Violet Cell Proliferation Kit (C34557; ThermoFisher), incubated for 10 minutes at 37°C, then washed and resuspended in media in the presence or not of DOS. Cells were then cultured for additional 3 days and re-activated as above, followed by CD3 and CD28 staining, and CTV dilution detection. Gating was performed among the GFP^+^ effectively transduced population (reporter gene) and analysed at the CytoFLEX flow cytometer.

### CTV in shAMPK DOS treated senescent T cells

Purified human CD4^+^ T_sen_ cells were transduced with lentiviral vectors containing shScramble (as control) or shAMPKα siRNA expression construct and maintained in culture for 21 days. Cells were activated every ten days with anti-CD3 antibody and recombinant rhIL-2, as above. After one month of culture, cells were treated with DOS and stained with 5 µM Cell Trace Violet, then reactivated for additional 3 days. CTV Analysis on transduced GFP^+^ T cells was then performed at the cytofluorimeter.

### *In vitro* BrdU proliferation assay

Purified human CD4^+^ T_sen_ cells were stimulated for 48 hours with anti-CD3 antibody (0.5μg/ml) plus rhIL-2 (10 ng/ml), then transduced with shSestrin1/2/3 lentiviral vectors as previously described (6). Ninety-six hours post-transduction, senescent CD4^+^ T cells were treated with BrdU (5-bromo-2′-deoxyuridine; 10µM; B23151-ThermoFisher) for 72 hours; then permeabilized and stained with anti-BrdU primary antibody (1:100; ab6326; Abcam) to assess proliferation by flow-cytometry.

### β-galactosidase staining

Purified T_sen_ were cultured for one week in a plate coated with anti-CD3 antibody (0.5 μg/ml) with the addition of rhIL-2 (5 ng/ml), in absence or not of DOS. One week later, 2*10^5^ cells were transferred on poly-lysine coated 96 well plates and stained using Senescence β-Galactosidase staining kit, following the manufacturer#s protocol (9860; Cell Signaling Technology). Cells were then imaged on a Zeiss-Primovert inverted phase-contrast microscope.

### AnnexinV staining

For analysis of cellular toxicity, human CD4^+^ T cells were treated with DOS, H_2_O_2_ (500mM; positive death control) or left untreated. Cells were stained using the Annexin V-FITC Apoptosis detection Kit (ab14085; Abcam) following the manufacturer protocol. Annexin levels were acquired by flow-cytometry. Annexin V staining was used also for detection of dead cells in CD45.1 mice adoptive transfer experiments.

### Seahorse assay for cell metabolism

To measure fatty acid oxidation, purified senescent CD4^+^ CD27^-^ CD28^-^ T cells were activated with with anti-CD3 antibody (0.5 μg/ml) plus rhIL-2 (5 ng/ml) and treated overnight with or without DOS. The day after, the media was replaced with XF Base Medium Minimal DMEM (103334-100; Agilent) composed of basic DMEM plus 0.5 mM glucose, 1 mM glutamine and 1% FBS. Cells were incubated with this media for 4 hours before plating with the assay media (composed of XF Base Medium Minimal DMEM supplemented with 2 mM glucose and 0.5 mM L-carnitine; Agilent) in the XF96 well plate (Agilent) pre-treated with poly-lysine at 2*105 cells/well. Cells were then incubated for 1 hour at 37⁰C without CO2. Just before the assay, cells were treated with 1X palmitate-BSA or with 1X BSA. According to the Palmitate Oxidation Stress Test protocol (Agilent), during the assay, cells were treated at different time points with Etomoxir (4 µM; 103672-100; Agilent), Oligomycin (1.5 µM; 103672-100; Agilent), FCCP (1 µM; 103672-100; Agilent) and Rotenone/Antimycin A (0.5 µM; 103672-100; Agilent). OCR values were acquired by Seahorse Machine (XFe96) over a time of 120 minutes.

For fatty acid oxidation, subtraction of etomoxir treated conditions was used to derive lipid burning as recommended by the manufacturer (Agilent). In some experiments, to further confirm DOS specificity of action, metabolic assessment was performed in senescent human CD4^+^ T cells that had been shRNA-depleted, or not, of all sestrins prior to overnight DOS treatment.

### Yellow fever specific T cell stimulation

Purified human CD4^+^ T cells were pre-treated with or without DOS for 4 hours; in parallel, autologous APCs were preloaded with or without yellow fever vaccine (1:200 of the human dose, Stamaril; Sanofi Pasteur) or specific YF peptides (2ug/ml: PepMix YF-NS4B; Innovative Peptide Solutions). In control experiments, APCs were loaded with negative control peptides (2ug/ml PepMix Human-Actin; Innovative Peptide Solutions). T cells and antigen-pulsed APCs were then co-cultured for 18 hours, in a 1:2 ratio. The next day, cells were treated for 4 hours with Brefeldin A (1 μg/mL; Sigma-Aldrich). CD4^+^ T cells stimulated for three hours with PMA (50 ng/ml) and ionomycin (1 ug/ml) were used as a control. Cells were stained for CD3, CD4, CD28, CD45RA, CD95, CD62L, and for intracellular TCF1 and IFN-γ detection. Antigen specific events were derived by subtraction of the unstimulated background, with or without DOS. Analysis was carried out by flow cytometry.

### TREC investigation

For polyclonal analysis of human TCR recombination, purified human CD4^+^ T cells were plated overnight, for 3 days, or for one week with CD3/CD28 Dynabeads, in the presence or not of DOS. For antigen specific TCR recombination instead, human CD4^+^ T cells were pre-treated with DOS for 4 hours and APCs pre-loaded with YF peptides (2ug/ml; PepMix YF-NS4B; Innovative Peptide Solutions) for the same time. The T cells were then co-cultured to allow synapse formation with APCs, in a 1:2 ratio. Antigen-specific TRECs were analyzed 18 hours later by background subtraction of antigen free immune conjugates with or without DOS, as above described.

For both experiments, DNA was isolated following TRIzol reagent protocol (15596018; ThermoFisher) and quantified with NANO-DROP spectrophotometer (Thermo-Scientific). qPCR was performed with 100 ng of gDNA, 900nM primers, 250nM TaqMan probe per sample according to the SensiFAST Probe Hi ROX kit (BIO-82005 Bioline).

To investigate TRECs, the following primers were used:

Fw CACATCCCTTTCAACCATGCT

Rw GCCAGCTGCAGGGTTTAGG

Probe [6FAM] ACACCTCTGGTTTTTGTAAAGGTGCCCACT [TAM]

The qPCR reaction was conducted by using !StepOne Plus System” machine (Applied Biosystems).

For analysis of mouse TCR recombination, murine CD4^+^ T cells were purified from peripheral lymphoid organs of mice treated or not with DOS. Cells were left to rest in culture overnight and the day after they were assayed for the TREC expression according to the above described procedure for human TREC investigation. For mouse TRECs, the following specific primers were used:

Fw CCAAGCTGACGGCAGGTTT

Rw AGCATGGCAAGCAGCACC

Probe [6FAM] TGCTGTGTGCCCTGCCCTGCC[TAM]

The same procedure was used to measure TRECs in CD45.1^+^ CD4^+^ adoptively transferred T cells in CD45.2 recipients.To investigate how TREC production could be affected by depletion of Rag proteins, 2 millions of purified CD4^+^ T cells were electroporated in the presence of siRNA for RAG1 (sc-42962; SantaCruz) and RAG2 (sc-36371; SantaCruz). siRNA scramble (sc-44236; SantaCruz) was used as control. Electroporation was performed according to the Human T cell nucleofactor kit protocol (Lonza) as previously described^2^. Cells were left for 60 hours in culture post electroporation with exposure or not to DOS for the remaining 4 hours of silencing prior to conjugation with autologous YF pulsed APCs, in a 1:2 ratio. For APC antigen loading, YF peptide mix was used (2ug/ml; PepMix YF-NS4B; Innovative Peptide Solutions). In parallel, conjugates were left unstimulated with or without DOS. After the overnight coculture, CD4^+^ T cells were purified and genomic DNA was extracted according to TRIzol reagent protocol (15596018; ThermoFisher). The DNA was quantified with NANO-DROP spectrophotometer (Thermo-Scientific). TREC levels were investigated by qPCR according to the protocol above described. Antigen specific TRECs were derived subtracting the raw data originated from unstimulated cultures with or without DOS, to identify the events occurring specifically in response to the DOS when combined with the YF peptides, or any YF specific events without DOS respectively.

### TCR Sequencing

Human APCs and CD4^+^ T cells were purified from PBMCs. Six million APCs were preloaded with YF peptides (2ug/ml; PepMix YF-NS4B; Innovative Peptide Solutions) or left unstimulated for 4 hours and then co-cultured with three million CD4^+^ T cells that had been pre-treated or not with DOS for the same time.

In parallel donor matched cultures, EBV peptides (1:50; peptivator EBV consensus;) were used instead of YF. After the overnight co-culture, CD4^+^ T cells were purified and genomic DNA was extracted according to the “PureLink DNA mini kit” (Invitrogen). DNA concentration was acquired at the NanoDrop spectrophotometer (ThermoFisher). This experiment has been performed in parallel in four donors to have biological replicates.

DNA samples were sent out for direct *TCR*β sequencing. *TCR*β amplification was carried out according to Robins et al. (2009)^4^ by using Qiagen Multiplex PCR kit and the reported primers. The method is capable to detect unique TCR repertoire, including the rarest receptor sequences. For indexing, KAPA HiFi HotStart ReadyMix (Roche) and Dual Index Set (NEB) were used. The libraries were sequenced on an Illumina MiSeq platform using 2×250bp paired-end sequencing with a coverage of ten million reads.

Fastq files were first subjected to quality control using FastQC (v0.11.9)^5^. TCR repertoire analysis and error correction were then performed using MiXCR (v4.2.0)^6^. Filter for low quality reads and adapter trimming was further performed using trimmomatic software (v0.39). Plots such as heatmaps were generated using the R programming language^7^. To identify antigen-specific (YF or EBV) TCR rearrangements, antigen stimulated or DOS-juvenated samples were subtracted of the unstimulated background controls. To distinguish antigen-specific DOS-driven TCR V-DJ and DJ rearrangements, DOS treated samples without antigen were considered as background. This analysis allowed to isolate the YF or EBV specific TCR modifications of either untreated or DOS-treated samples. Similar TCR genomic analysis were described elsewhere^8^. In addition, individual VDJ clonotypes and clonotype usage, as well examples of unique *de novo* antigen-specific clonotype formation are also shown across all conditions. Software generated VDJ tools (Java V1.2.1) were used for clonotype analysis.

### TRBV clone validation

Donor-matched YF-specific conjugates were formed as above described (“**Yellow fever specific T cell stimulation**”) for 18 hours, in the presence or in the absence of DOS. The day after, cells were treated with brefaldin A (1:1000; Sigma-Aldrich) for 4 hours. Cells were stained for the surface proteins CD4, CD28, CD45RA, CD62L, CD95, and intracellularly, for TCF1 and IFNγ, as indicated. In addition, expression of donor-matched YF-specific TCR variants identified during Next Generation TCR Sequencing was validated by flow cytometry with variant specific antibodies to TCRvβ2, 7.1, 14, 16 and 17.

For the mouse side, TCR rearrangements were assessed in CD4^+^ T cells purified from lymphoid organs of old mice treated or not with DOS and then infected or not 2 months later with H1N1 virus (3.5*10^5^ PFU). Cells from young mice infected with the same virus were used as control. All mouse T cells were stained for the surface TCR variants vα8.3 and vβ12, previously reported to be associated with the anti-Flu response^9^.

For the purification of individual TRBV clones, NGS-matched human CD4^+^ T cells were pre-treated with or without DOS then co-cultured with autologous APCs pre-loaded with YF-vaccine or peptides for 18 hours, as indicated. Post-synaptic CD4^+^ T cells were then gently detached from the conjugates using a 2 ml syringe with cold PBS and processed for sorting. For cell sorting, post-synaptic CD4^+^ T _STEM_ were stained with antibodies directed to CD45RA, CD28, CD95 and CD62L surface markers in combination with a single TRBV clonotype antibody as follows: TCRvβ2^+^, TCRvβ7.1^+^, TCRvβ14^+^, TCRvβ16^+^, TCRvβ17^+^. In parallel, TEM and TCM for each TCRvβ clonotype were also sorted based on the relative expression of CD45RA and CD28 surface markers. Cells were sorted with a MoFlo Astrios-EQ flow cytometer (Beckman Coulter). Isolated TRBV clones (2*10^5^ each) were then rested for 5 days followed by re-stimulation with YF peptide (2 ug/ml) or YF-vaccine (1:200 of the human dose) for 18 hours. Cell supernatants were assessed with Human IFN γ ELISA Kit (abcam; ab46025) according to the manufacturer’s instructions. T_EM_ and T_CM_ IFNγ supernatants were measured as negative controls.

To assess YF-specific TRBV T_STEM_ clone proliferation, CD4^+^ T_STEM_ (2*10^5^ each) from every TCRvβ clonotype subset that had been derived and sorted as above were incubated with BrdU labeling solution for 4 hours, followed by re-challenging with autologous YF-pulsed APCs for additional 4 days. TCRvβ CD4^+^ T_STEM_ cells were tested with the BrDU Cell proliferation ELISA kit (abcam; ab126556) according to the manufacturer’s instructions. Sorted TCRvβ T_EM_ and T_CM_ were tested in parallel as negative controls.

### Telomere live transfer

Ten million of purified human APCs with live labelled telomeres were allowed to form synapses with autologous T_sen_ (3:1 ratio) in the presence or in the absence of DOS and stimulated overnight with a 1:1 mix of CMV and EBV antigens (Peptivator CMV pp65 130-093-438 and Peptivator EBV Consensus 130-099-764; Miltenyi). The day after were stained for CD3, CD4, CD27, CD28 receptors and analysed by flow-cytometry for transferred APC telomere fluorescence into recipient T cells.

### Telomere Length Quantification Assay

Purified human APCs (4*10^6^) and T_sen_ (1*10^6^) were conjugated with CMV and EBV antigens (3:1) and treated or not overnight with DOS. The day after, cells were detached from conjugates with a 2ml syringe and cold PBS. After a CD4^+^ T cell purification as above described, genomic DNA was extracted from senescent T cells according to PureLink Genomic DNA Mini Kit protocol (K182001; Invitrogen). qPCR reaction was performed in the extracted DNA according to the protocol suggested by the !Absolute human telomere length quantification qPCR Assay kit” (8918; ScienCell Research Laboratories). Telomere length was calculated from the ΔCT, according to the formula suggested by the kit.

### Telomerase Activity

CD4^+^ T_sen_ were purified from human PBMCs. Purified T_sen_ were plated in an anti-CD3 coated plate in the presence of IL-2 (5ng/ml) and left in culture for 72 hours with or without DOS. Cell pellets from the three different cell types were assayed for telomerase activity according to the TeloTAGGG Telomerase PCR ELISA Kit (11854666910; Roche).

### DOS_juv_ CD4^+^ T cell Adoptive transfer

Twenty months old mice (C57BL/6 wild type CD45.1) were primed with Fluad (1:20 of the human dose; Seqirus) and subcutaneously injected with or without DOS (0,1mg/Kg). After 15 days, mice were culled and peripheral lymphoid organs collected. Splenic donor CD4^+^ T cells were purified, stained with CTV and phenotype before injection into young naïve congenic recipients (3 months). Mice not receiving donor cells were used as negative transfer control. Twenty-eight days later, mice were sacrificed, organs collected and CTV^+^ cells analyzed at the CytoFLEX flow cytometer. Alternatively, aged-matched CD45.1 mice (20 months) from vaccinated donor animals were derived as above described and injected without CTV labelling into CD45.2 young recipients (3 months) to allow tracking after transfer independently of CTV. No vaccination of recipients was performed in either protocol, with similar results. Ten million or 5 million cells were used for adoptive transfer, with similar results, as indicated.

To study the role of CD4^+^ T cells on driving DOS effects, old CD45.1 mice (20 months) were injected or not with DOS. Seventy-two hours later animals were culled and CD4^+^ T cells were isolated from their spleens and then intravenously injected into aged-matched CD45.2 recipient mice (5×10^6^ CD4^+^ T cells per mouse). Vaccination with Fluad was performed or not 24 hours after transfer as indicated below.

Eight weeks later, all mice were exposed to the H1N1 flu virus (5×10^5^ PFU) and then daily monitored for the next 15 days. During this period, clinical parameters were acquired: fur condition, posture, movement, breathing, eye state, and weight loss to assess a clinical score.

Once reached a total score of 13 or more, mice were humanely euthanized accordingly to the animal ethical guidelines; animals with good parameters were instead euthanized 15 days post infection. Several tissues (blood, lung, spleen, and lymph-nodes) were collected from all the animals for further analysis.

Experimental mouse groups:

- 5 old CD45.2 mice (20 month old) receiving age-matched DOSjuv CD45.1^+^ CD4^+^ donor T cells and vaccinated with Fluad.
- 5 old CD45.2 mice (20 month old) receiving age-matched CD45.1^+^ CD4^+^ donor T cells (DOS-free) and vaccinated with Fluad
- 5 old CD45.2 mice (20 month old) receiving young CD45.1^+^ CD4^+^ donor T cells and vaccinated with Fluad
- 5 old CD45.2 control mice (20 month old) transfer free injected with vehicle and vaccinated with Fluad
- 5 young CD45.2 mice (3 month old) transfer free injected with vehicle and vaccinated with Fluad without DOS
- 5 old CD45.2 mice (20 month old) receiving age-matched DOS_juv_ CD45.1^+^ CD4^+^ donor T cells without vaccination

### sMAC expression in human T cells

Purified human CD4^+^ T cells were stained for the surface proteins CD4, CD28, CD45RA receptors and for the intracellular proteins sestrin 2 and phopsho-p38 directly *ex vivo* and analysed by cytofluorimetry. sMAC expressing cells as considered as sestrin and p-p38 positive populations.

### *In vitro* antigen-specific IgG production

PBMCS were treated with or without DOS then cultured for 14 days with Fluad vaccine (1:50 human dose; Seqirus). In some experiments, human CD4^+^ T cells were depleted from the immune cultures by magnetic isolation prior to Fluad stimulation, to assess impact of these cells on the vaccine response upon DOS treatment. Fluad vaccines (1:40 of the human dose) were coated on 96 wells overnight at 4°C and then blocked with 4% milk in PBS 0.5% Tween-20, 2 hours at room temperature. 100ul of supernatant was then added and left incubating overnight at 4$C. Negative controls were coated with PBS 1X instead of Fluad vaccine and used as background throughout. Furthermore, in parallel coating, unvaccinated serum was used as further negative control for presence of unrelated IgGs. The plate was washed and then incubated with anti-mouse IgG1 biotin conjugated antibody (1:250; 130-095-879; Miltenyi) for 2 hours at room temperature. After washing and incubation with streptavidin-HRP conjugated antibody (1:3000; 1610381; Biorad) for 1 hours at room temperature, TMB substrate was added. The reaction was stopped with the stop solution. The absorbance was detected at 450 nm at the ELISA micro-plate reader (AMR-100, Biolab).

### RNA extraction, retro-transcription and qPCR

Total RNA from CD4^+^ T cells purified from mouse spleen, lymph nodes or lung cells was extracted using the !PureLink RNA Mini Kit” (12183018A; Invitrogen). The RNA concentration was acquired by NANO-DROP spectrophotometer (Thermo-Scientific). One µg of total RNA was retro-transcribed to cDNA following the protocol advised from the kit !qScript cDNA synthesis kit” (733-1174; Quanta Biosciences). qPCR reaction was performed with 100 ng of cDNA per sample using the kit SensiFAST SYBR Hi ROX kit (BIO-92005; Bioline).

For investigation of mouse *sestrin 2* the following primers were used:

Fw TCTCGGCACTTTGAGGACAC

Rw AACCATGGTCTTCCCAGCAG

In parallel, to investigate the expression levels of other genes of interest, the TaqMan approach was used. In this context, qPCR reaction was performed with 100 ng of cDNA per sample using the kit SensiFAST Probe Hi ROX kit (BIO-82005; Bioline). The expression of *IL7RB* was assessed with the TaqMan probe Mm00434295 (ThermoFisher), while for the house-keeping gene, β*-actin*, the probe Mm02619580 was used (ThermoFisher).

To investigate the presence of the H1N1 virus in the lung of infected mice, total DNA was purified from the tissue according to the TRIzol reagent protocol (15596018; ThermoFisher), quantified and amplified for the PA viral gene of H1N1 virus with the following primers:

Fw CGGTCCAAATTCCTGCTGA

Rw CATTGGGTTCCTTCCATCCA

The following primers were instead used to amplify the house-keeping gene, *RPL34,* used as control:

Fw GGTGCTCAGAGGCACTCAGGATG

Rw GTGCTTTCCCAACCTTCTTGGTGT

All qPCR reactions were conducted by using !StepOne Plus System” machine (Applied Biosystems).

### Chromatin Immune Precipitation (Chip)

For detection of murine HDAC1 activity, chromatin was immunoprecipitated from peripheral lymphoid organs of old mice (20 months) treated or untreated with DOS and from their young controls. All mice were H1N1 infected 2 months after DOS treatment (DOS only protocol, see above). Chip was performed according to the manufacturer’s instructions with anti-HDAC1 antibody. The Chip reactions were then incubated for 1 hour at 30$C with an exogenous DNA purified from peripheral lymphoid organs of naïve old mice (20 months). HDAC1 activity was measured by ELISA assay detecting the level acetylated Histone 3 (anti-acetil-H3; K27; 8173S; Cell Signaling).

### *Sestrin* 2 gene acetylation

To investigate epigenetic changes induced by DOS treatment *in vivo,* the chromatin was immune-precipitated from peripheral lymphoid organ cells of old mice treated or untreated with DOS and from their young controls following the manufacturing protocol of the Pierce Chromatin Prep Module kit (26158; Thermo Scientific). *Sestrin 2* gene is mainly acetylated at histone 3, for this reason an anti-histone 3 antibody (anti-acetil-H3; K27; 8173S; Cell Signaling) was used for the immuneprecipitation. The antibody anti-5 methylcytosine was used in parallel as control. After DNA purification *sestrin 2* expression was assessed by qPCR with primers specific for the mouse *sestrin 2*. The following primers were used: Fw AGGAGTGTCTATCGCCGTAGCG

Rw TGGGACACCCGAGGGGTCTAGGT

qPCR reaction was performed using the kit SensiFAST SYBR Hi ROX kit (BIO-92005; Bioline) on a “StepOne Plus System” machine (Applied Biosystems).

### DOS kinetics

Mice (strain C57BL/6) were divided in 2 groups:

1. 5 old mice (18 month old) injected subcutaneously with DOS 46L at 0.1mg/kg
2. 5 old mice (18 month old) control subcutaneously injected with vehicle
3. 5 young mice (3 month old) control

Mice were bled at 48, 72, 96 hours and 10 days post injection. Sestrin2 levels were monitored by flow-cytometry.

### Mouse Survival

C57BL/6 mice were subcutaneously injected with Fluad vaccine (1:20 human dose; Seqirus) and divided in the following groups:

- 7 old mice (18 month old) vaccinated with Fluad without DOS
- 8 old mice (18 month old) vaccinated with Fluad + DOS 46L (0,1 mg/Kg)
- 7 young mice (3 month old) vaccinated with Fluad without DOS

Six months post vaccination, mice were then infected with H1N1 flu virus (6*10^5^ PFU) and then monitored for the following 15 days: body weight, disease progression, clinical score and survival data were acquired. After 15 days, mice were sacrificed and tissues were collected (blood, lung, spleen and thymus). Clinical score included: fur condition, posture, movement, breathing, eye state, and weight loss according to ethics and under veterinary supervision. Animals were sacrificed when reaching a clinical score of 13 or above (grade of severity accepted from the ethic commission), or in any case of severe dyspnea.

### Determination of antigen specific IgG production *in vivo*

Identification of IgGs levels on mouse plasma was performed at 18, 36 days and 8 weeks after mouse immunisation with Fluad vaccine (1:20 of the human dose; Seqirus) by ELISA assay as above described. For viral specific antibodies, H1N1 live particles were used for coating instead of Fluad. IgGs were investigated at the same way in mouse blood of the DOS-juvenated adoptive CD4^+^ T cell transfer experiment.

### Micro-neutralization Assay

Mouse heat-inactivated sera were diluted 1:200 and pre-incubated with H1N1 virus (1:10000 dilution) for 1 hour at 37$C. MDCK cells (CCL-34; ATCC) were added to sera-viral mix at concentration of 15.000 cells per well (96 well plate) and incubated overnight at 37$C. The following day, cells were first fixed with cold Acetone 80% in PBS and then incubated for 1 hour at room temperature with anti-influenza A NP antibody (1:250; ab20343; Abcam). Secondary antibody was goat anti-mouse IgG HRP labelled (1:2500; Millipore). The reaction was developed by ELISA as above. Percentage of death inhibition was calculated setting the OD450 value of positive dead cell control as 0% and OD450 value of negative null death as 100%.

### Mice treated only with DOS and infected with H1N1 virus

C57BL/6 mice were divided in the following groups:

- 10 old mice (18 months old) treated with DOS46L (0.1mg/kg)
- 10 old mice (18 months old) not injected with DOS
- 10 young mice (3 months old) not injected with DOS

Mice were intra-nasally infected with H1N1 viral particles (3.5*10^5^ PFU) either 48 hours (immediate infection) or 8 weeks (delayed infection) after subcutaneous DOS administration (0.1 mg/Kg). Mice were monitored for 15 days and clinical score parameters were recorded: fur condition, posture, movement, breathing, eye state, and weight loss. Animals were sacrificed according to clinical score threshold of 13 or in any case of severe dyspnea, as above. Tissues were collected from each mouse (blood, lung, spleen, lymph-nodes and thymus).

### Adoptive transfer for B cell phenotype

Twenty-month-old mice (CD45.1 C57BL/6 strain) were subcutaneously injected or not with DOS. After 72 hours, mice were culled and spleens were collected; CD45.1 CD4^+^ T cells and all other immune cells depleted of CD4^+^ T cells were purified. Complete CD4^+^ T cell depletion was confirmed into the CD4^-^ T cell pool by flow-cytometry. Each CD45.2 recipient mouse received 5*10^6^ of purified CD45.1 CD4^+^ T cells or CD4^-^ cells via caudal vein injection. Young CD4^+^ T cell from naïve animals were also transferred. Young and old and DOS treated mice, free of transfer, were used as control. The day after transfer, CD45.2 recipient mice were subcutaneously vaccinated with Fluad (1:20 of the human dose). Fifty-teen days later, recipient mice were bled to investigate Fluad specific IgGs, before proceeding with animal sacrifice and collection of the peripheral lymph nodes. B cells from lymph nodes were stained for the following cell surface markers: CD19, CD45R (B220), CD138, GL7. CD3^+^ CD4^+^ and CD3^+^ CD8^+^ T cells were excluded from gating.

For IgG1 quantification, a standard curve was obtained with well-defined recombinant IgG1 protein (ab180055; abcam) concentrations and compared against IgG1-containining mouse sera from vaccinated animals, run in parallel.

### *In vitro* B cell phenotype

CD4^+^ T cells were derived from twenty-month old mice (C57BL/6 strain) and stimulated for 48 hours with CD3/CD28 Dynabeads (11452D, Gibco). The cells were then transduced with p-SIREN-sh lentivirus particles to target sestrins or with control p-SIREN scramble vectors, at a multiplicity of infection (MOI) of 10. After 72 hours, the transduced CD4^+^ T cells were cultured with autologous purified APCs in a 1:2 ratio. The co-cultures were stimulated with Fluad vaccine (1:100 of the human dose) and treated or not with DOS. After 10 days of culture, B cells were stained for the following surface proteins CD19, B220, CD138, CD38. T cells were excluded by gating out CD3^+^ CD4^+^ CD8^+^ lymphocytes.

### Cytokine production and T cell subpopulation induction in splenocytes

For primary Tfh responses, C57BL/6 mice were vaccinated with Fluad in the presence or in the absence of DOS and analyzed 14 days later. Splenocytes were then isolated from the spleen of the vaccinated young (3 months), old or DOS-treated mice (20 months) and stained for CD3, CD4, CXCR5 and PD1 markers. For the antiviral response, T cells from DOS only treated, H1N1 infected mice were re-exposed to H1N1 viral particles and Th1 state was analyzed 3 days later for the intracellular IFNγ and Tbet expression among CD3^+^ CD4^+^ T cells. Young mice were used as control. The same approach was used to investigate CD45.1 Tfh and Th1 responses in the DOS-juvenated adoptive CD4^+^ T cell transfer among CD45.2 recipient lymph nodes or lungs, as indicated.

### IFNγ production in response to Yellow Fever in mouse splenocytes

Splenocytes were collected from young mice (C57BL/6 strain; 3 months) 1 month after vaccination with YF vaccine (1:20 of the human dose; Stamaril-Sanofi). Unvaccinated young animals were used as control. Splenocytes were plated and left unstimulated or stimulated with YF peptide mix (2ug/ml; PepMix YF-NS4B; Innovative Peptide Solutions) or with the control peptide mix (2ug/ml; PepMix Human-Actin; Innovative Peptide Solutions). After overnight incubation, cells were treated with brefaldin A (1:1000; Sigma-Aldrich) for 4 hours and then stained for the T cell surface markers CD4, CD44, CD62L, CD95 and for the intracellular IFNγ. Data were acquired at the flow cytometer. In some experiments, induction of de novo YF specificity was further proved in OT-II transgenic mice that only respond to OVA. T cells were derived from OTII splenocytes and challenged with YF pulsed APCs in the presence or in the absence of DOS for 18 hours. The day after, stem T cells were purified by FACS sorting, rested for 5 days and re-challenged with APCs loaded with YF peptides. IFN gamma release was measured in the culture supernatant by ELISA (ab100689; abcam) 18 hours later. Positive control IFN gamma T cell release derived from OVA immunised animals, whose OT II CD4+ T cells were re-challenged with OVA in vitro 15 days after vaccination.

### Statistical analysis

A maximum of nine (human experiments) or 10 (mouse work) biological replicates for each experiment, with at least of three independent biological replicates throughout, as indicated. GraphPad Prism v9 was used to perform statistical analysis. For pairwise comparisons in human experiments, a two-tailed, paired Student#s t-test was used. For pairwise comparison in *in vivo* mouse experiments, a two-tailed, unpaired Student#s t-test. For multiple comparisons, a one-way analysis of variance (ANOVA) with a Bonferroni post-test correction was used. For survival analysis, Mantel-Cox. For analysis of small antigen-specific events (TREC monitoring and intercellular telomere transfer), an analysis of variance with F test was used. Similar statistic assessment was described previously^1, 2, 12^. Error bars indicate SEM throughout.

## Data availability

The data generated or analysed in this study are included in the manuscript, and its Supplementary Figures. Next generation TCR sequencing metadata are provided at submission. These data will be available with no restriction for reanalysis and reuse at the point of publication. Source data are available.

## Contributions

A.L. invented the DOS and further conceived of, designed and directed the study, analysed and interpreted the data, provided fundings and laboratory infrastructures and wrote the manuscript. C.D.A. F.R. and M.C. performed experiments and analysed individual data sets. L.C. synthesized sestrin mutant vectors. M.D.P. analysed human TCR sequencing under guidance of A.L. M.K. provided suggestions. All authors read and approved the final version of the manuscript.

## Acknowledgments.

We thank A. Sewell for discussions, and D. Escors and L. Fernandez Rubio and Sentcell for technical support. This work was funded by Sentcell ltd. The funder had no role in study design or decision to publish the manuscript. A.L. is a Honorary Professor of the University College London and the Chief Executive Officer of Sentcell ltd.

## Conflict of interest

A.L. is a shareholder to Sentcell ltd and the sole inventor of the DOS pharmaceutics where Sentcell ltd figures as the Applicant (PCT/IT2021/000059). M.K. serves as a scientific advisor to Sentcell ltd. A.L. M.C and F.R. are supported by Sentcell ltd.

