## Supplementary figures and images for "Disruptors of sestrin-MAPK interactions rejuvenate T cells and expand TCR specificity"

### Graphical abstract

#### Mechanism of DOS-juvenation

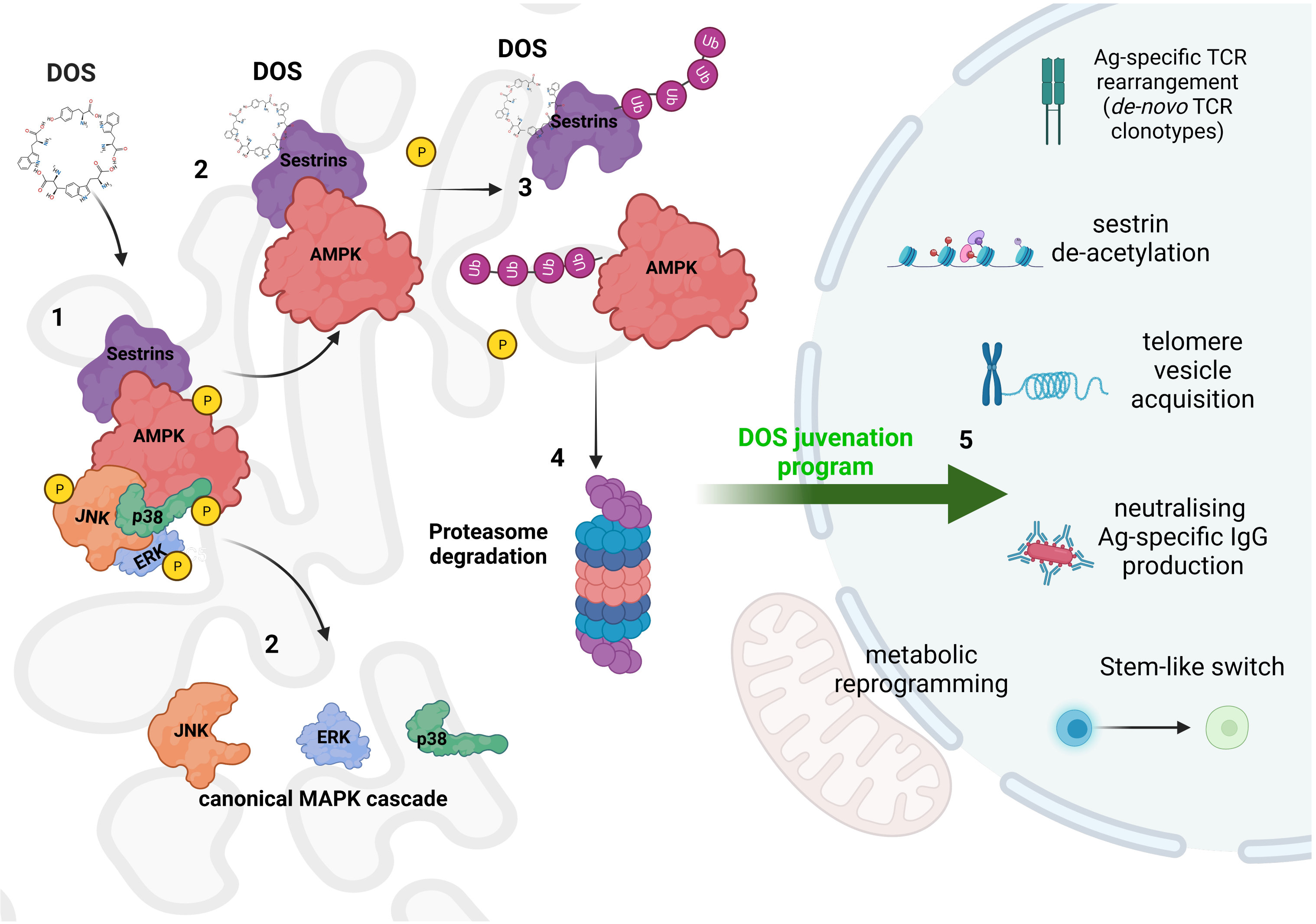

### S1

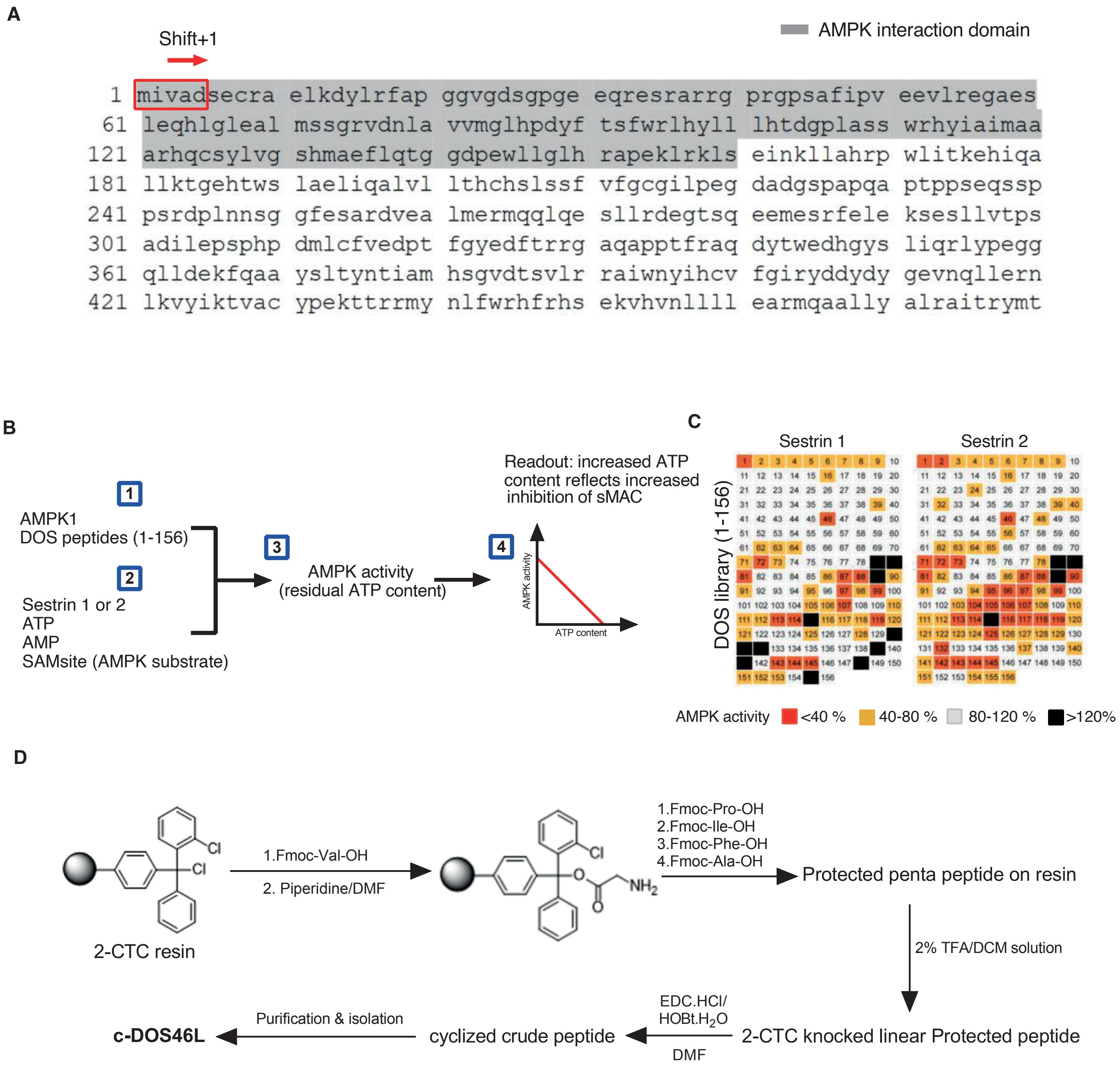

### S2

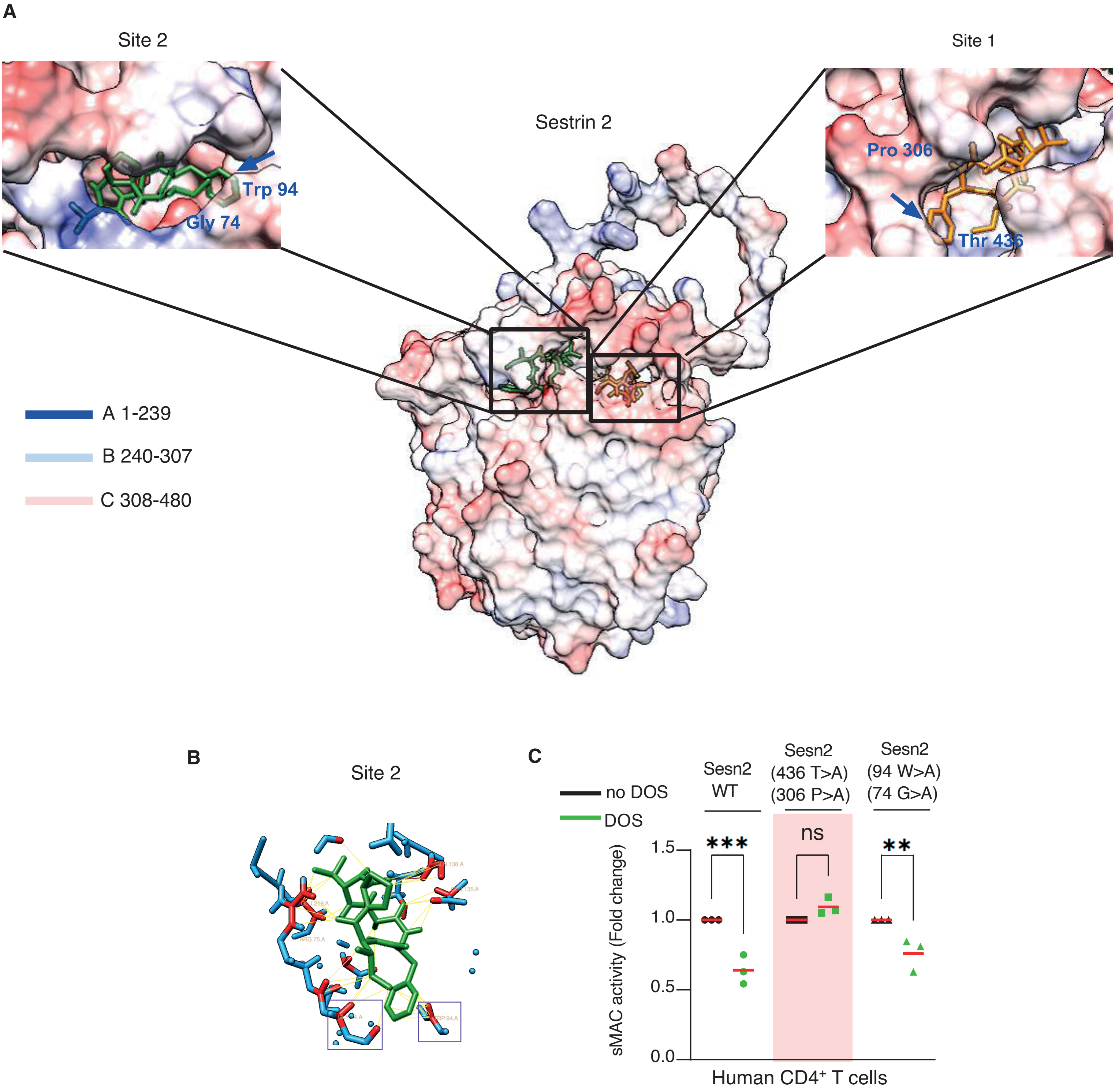

### S3

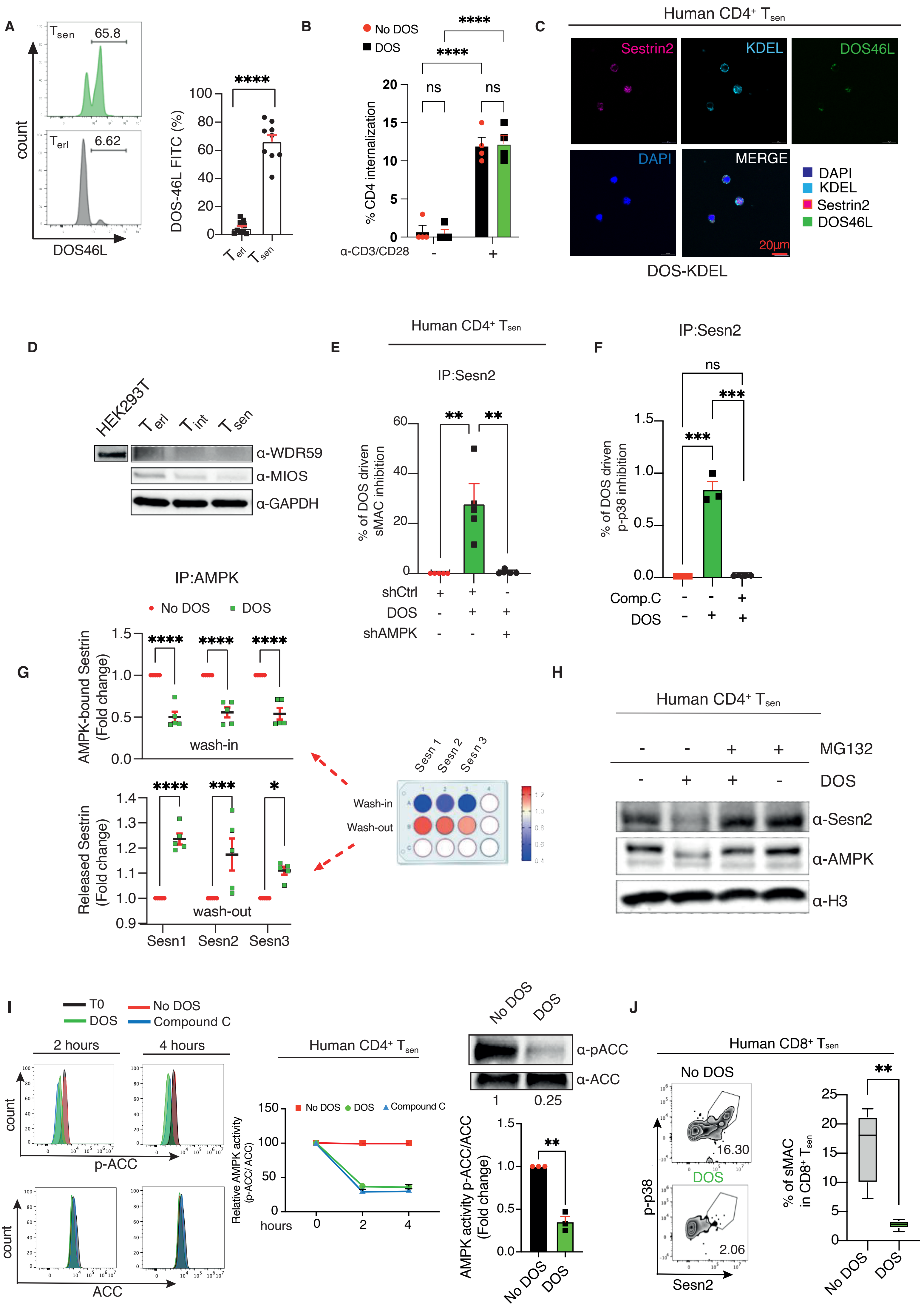

### S4

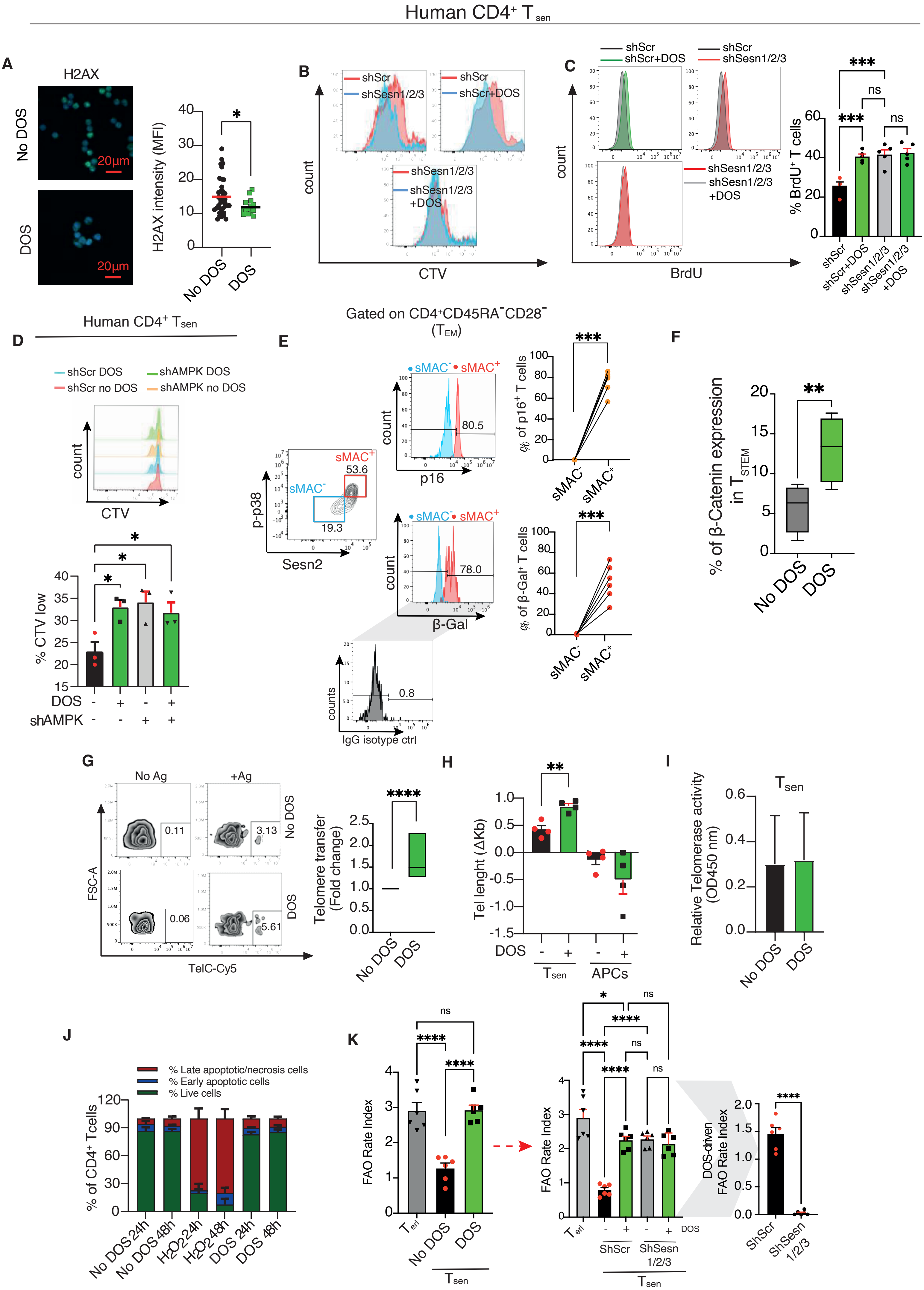

### S5

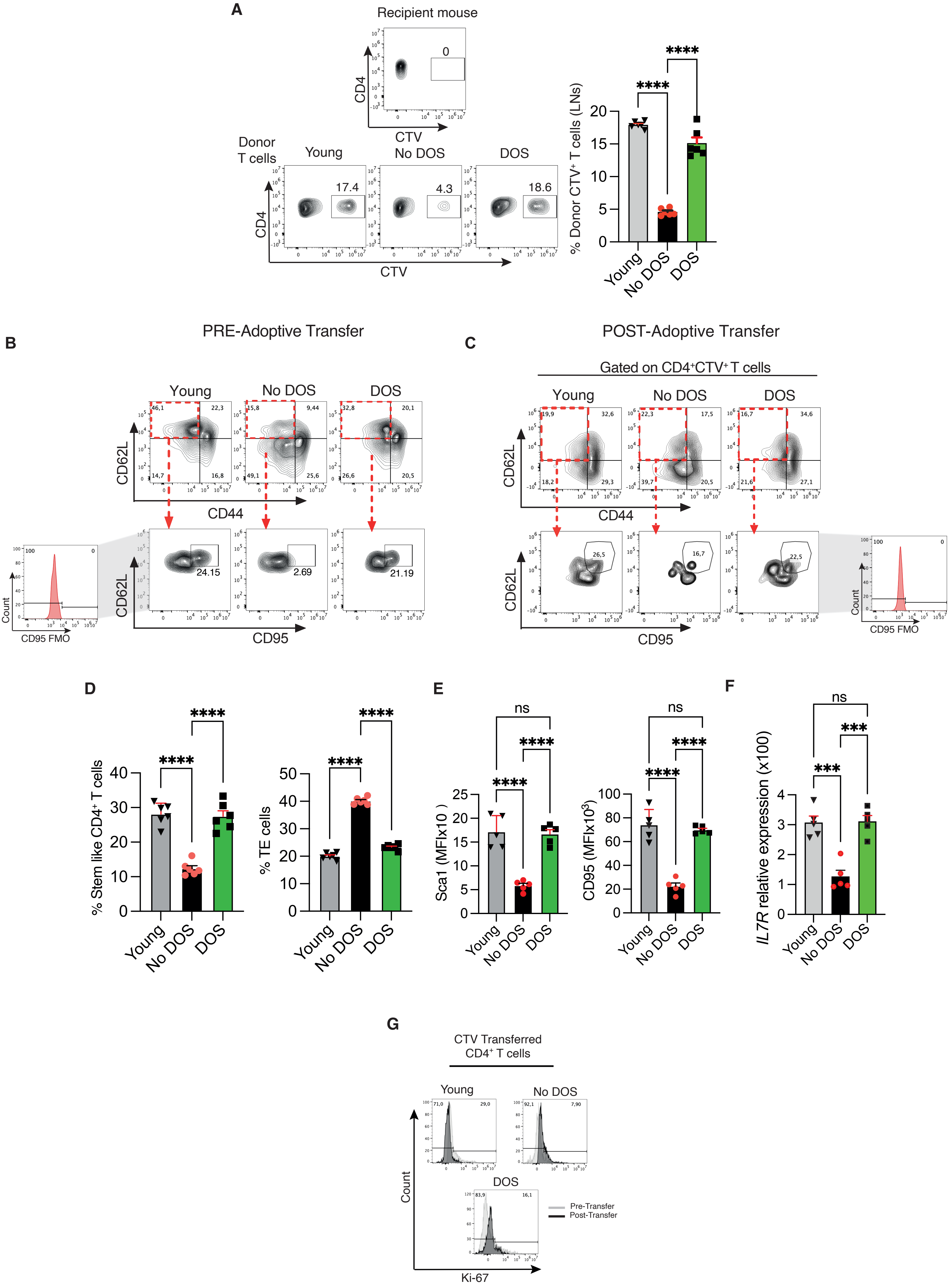

### S6

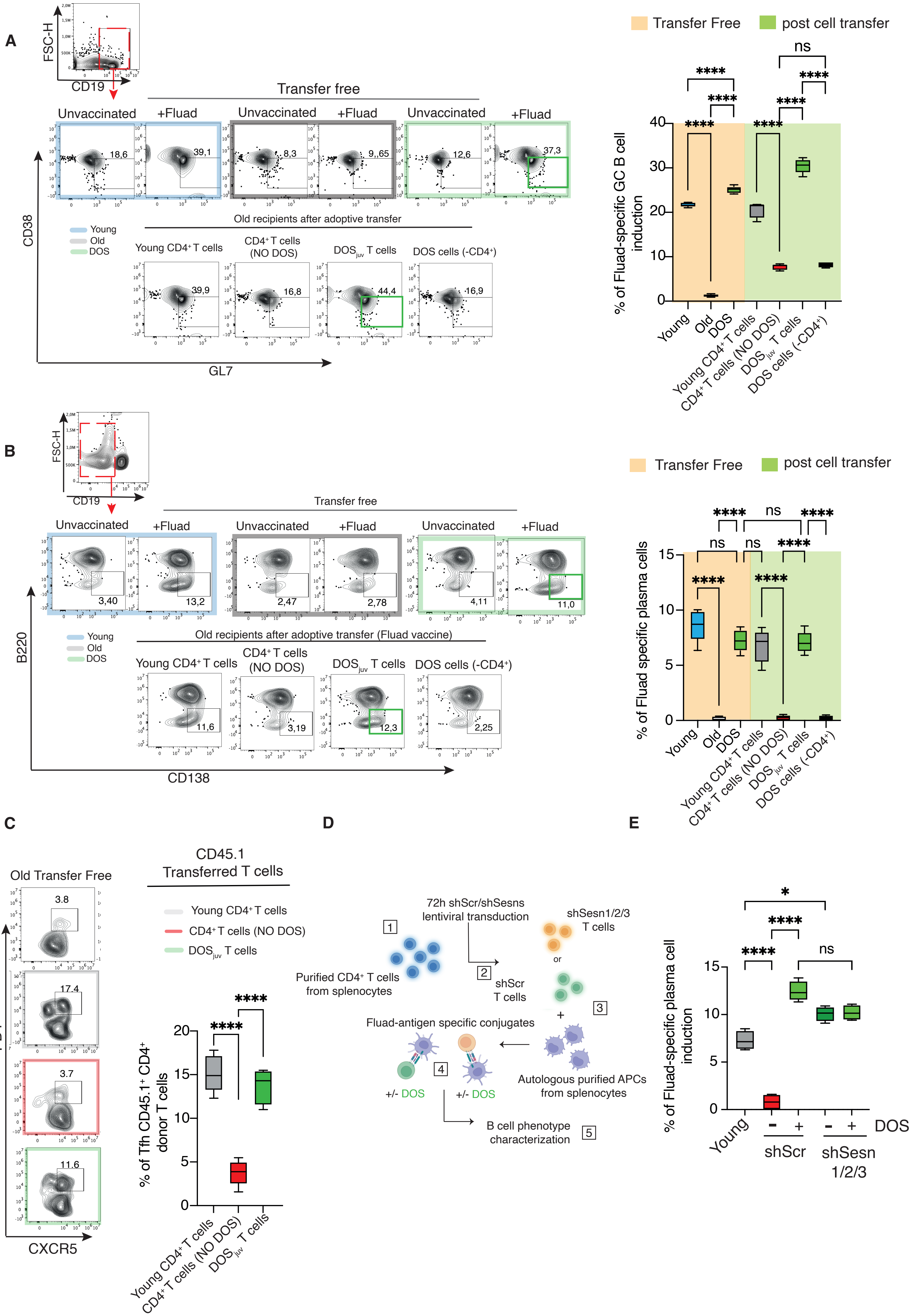

### S7

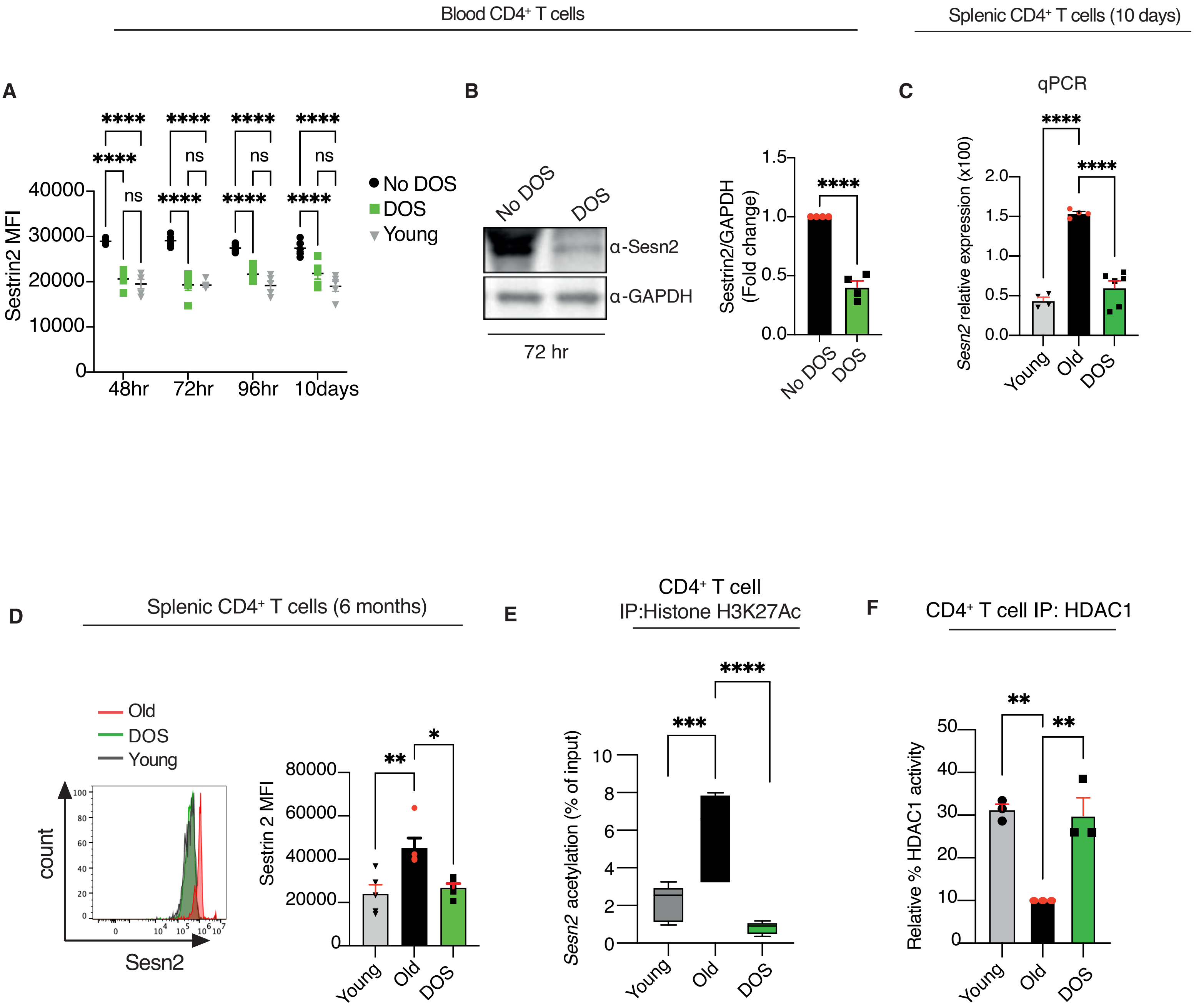

### S8

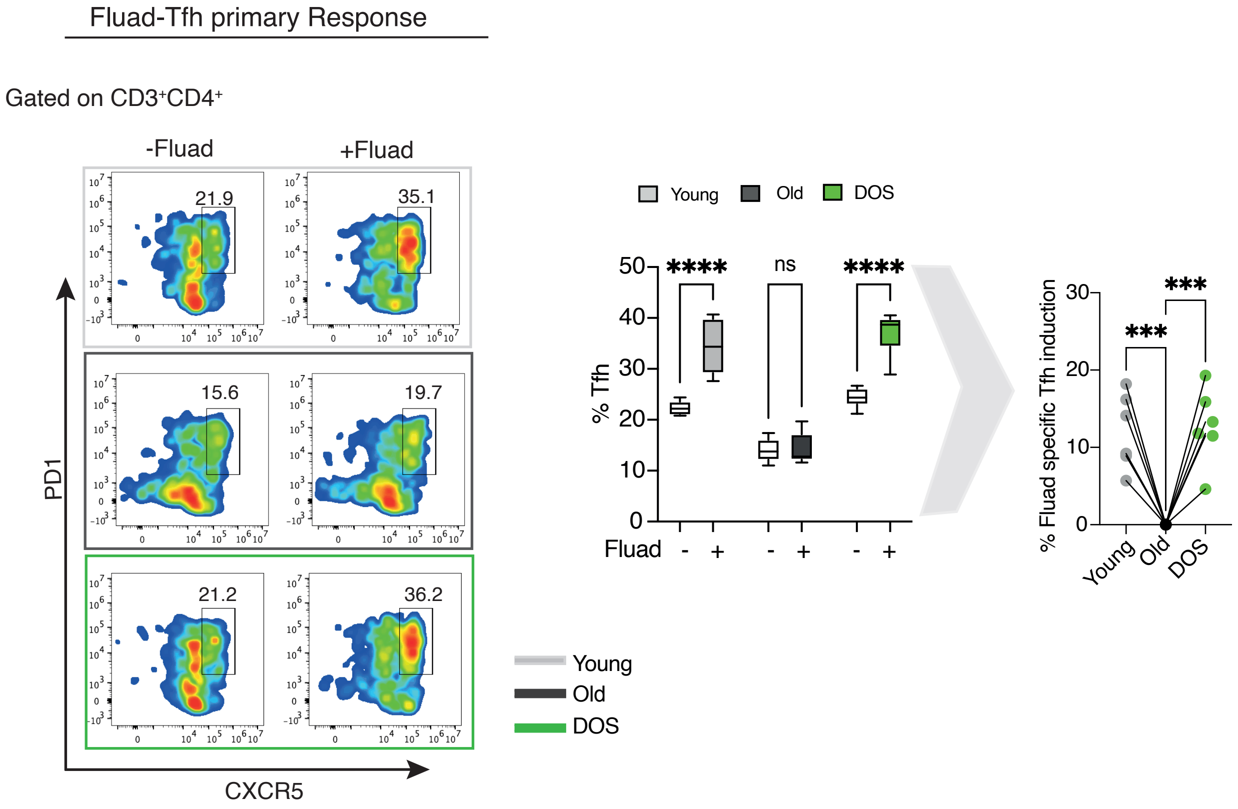

### S9

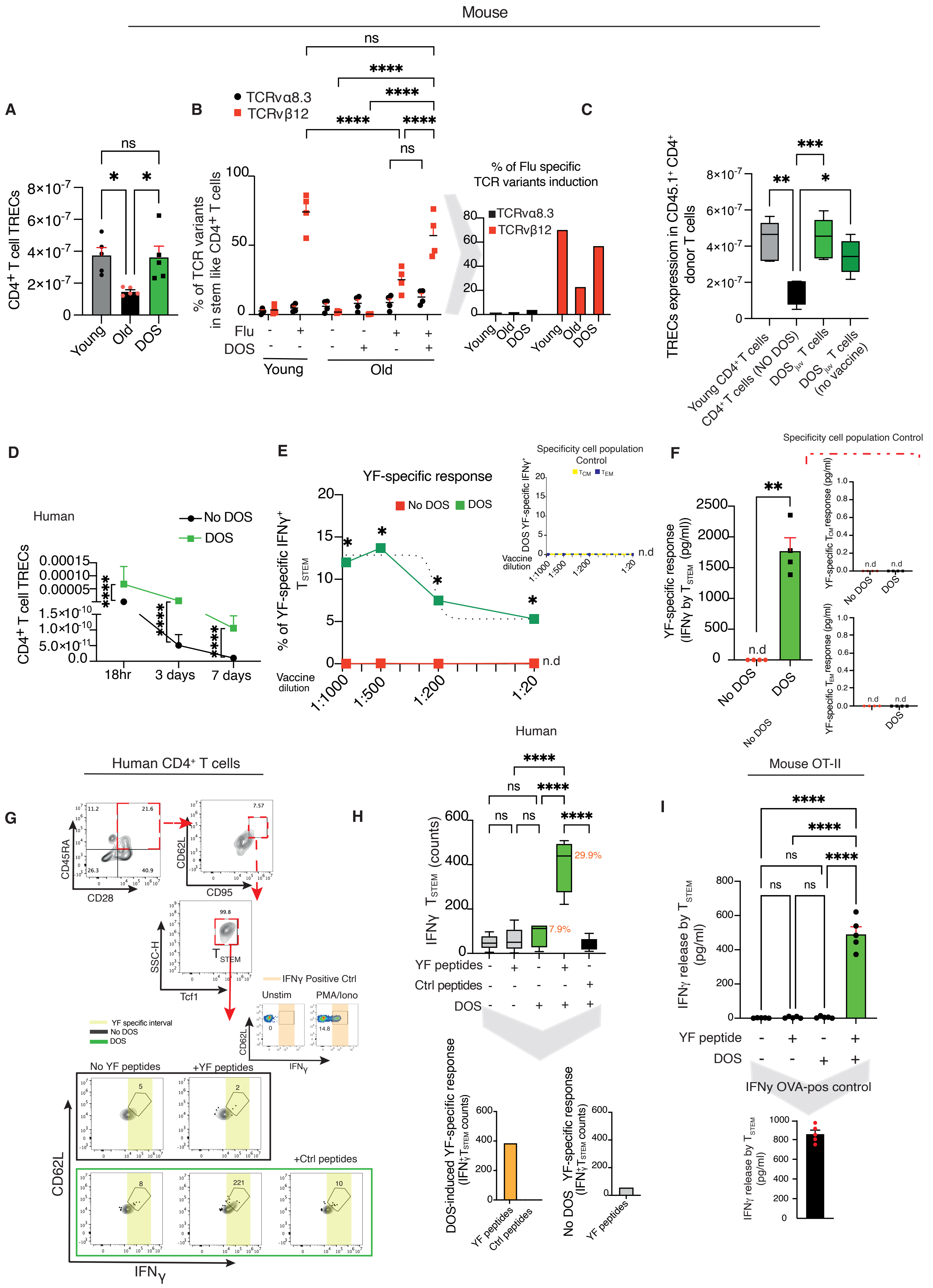

### S10

**A**

Donor A YF-induced TCRs

| Cell Subset | TCRVβ |
|---|---|
| $CD4^+$ T cell | V7-3 |
| $CD4^+$ T cell | V12-3 |
| $CD4^+$ T cell | V6-3 |
| $CD4^+$ T cell | V7-4 |
| $CD4^+$ T cell | V5-5 |

**B**

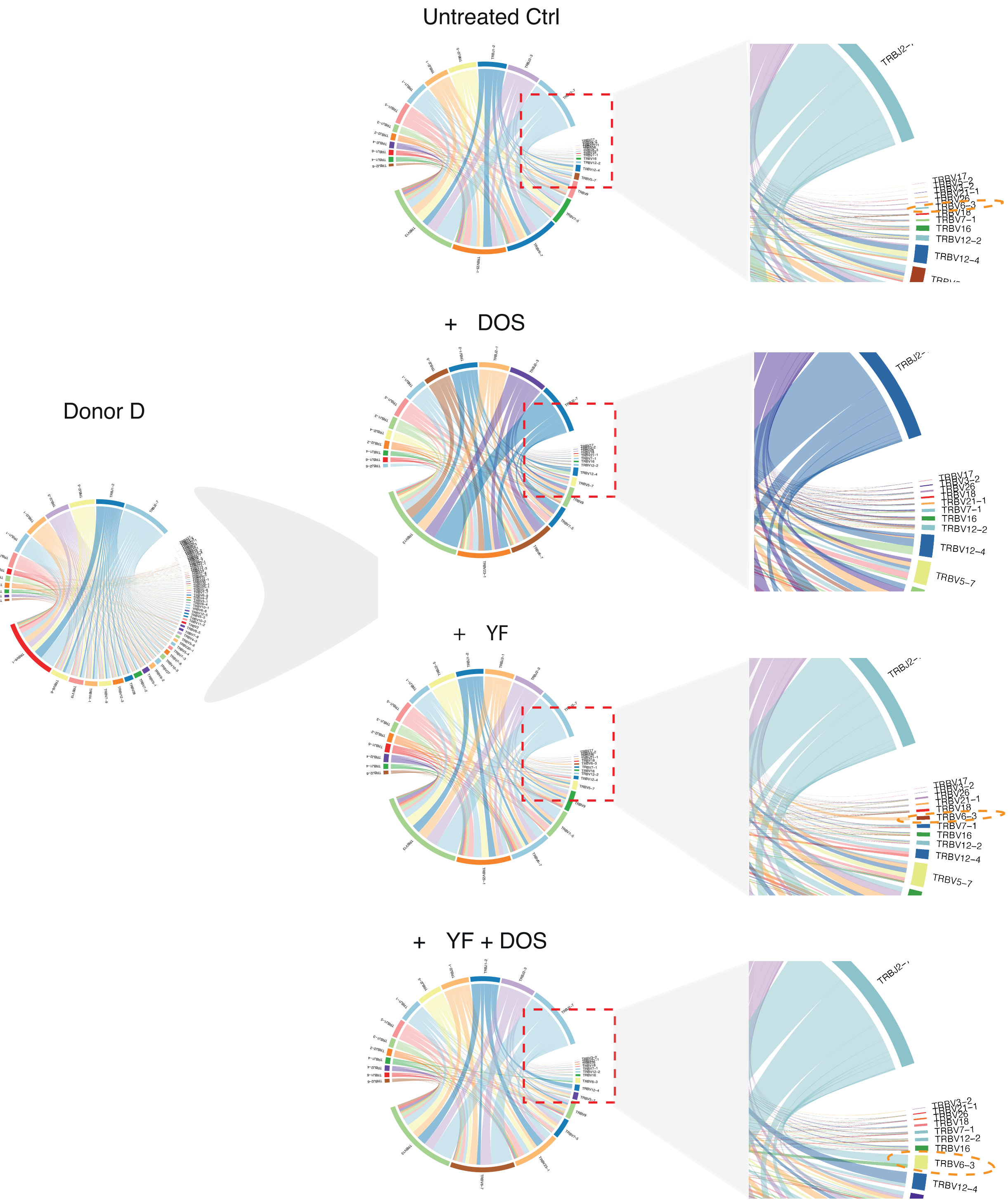

### S11

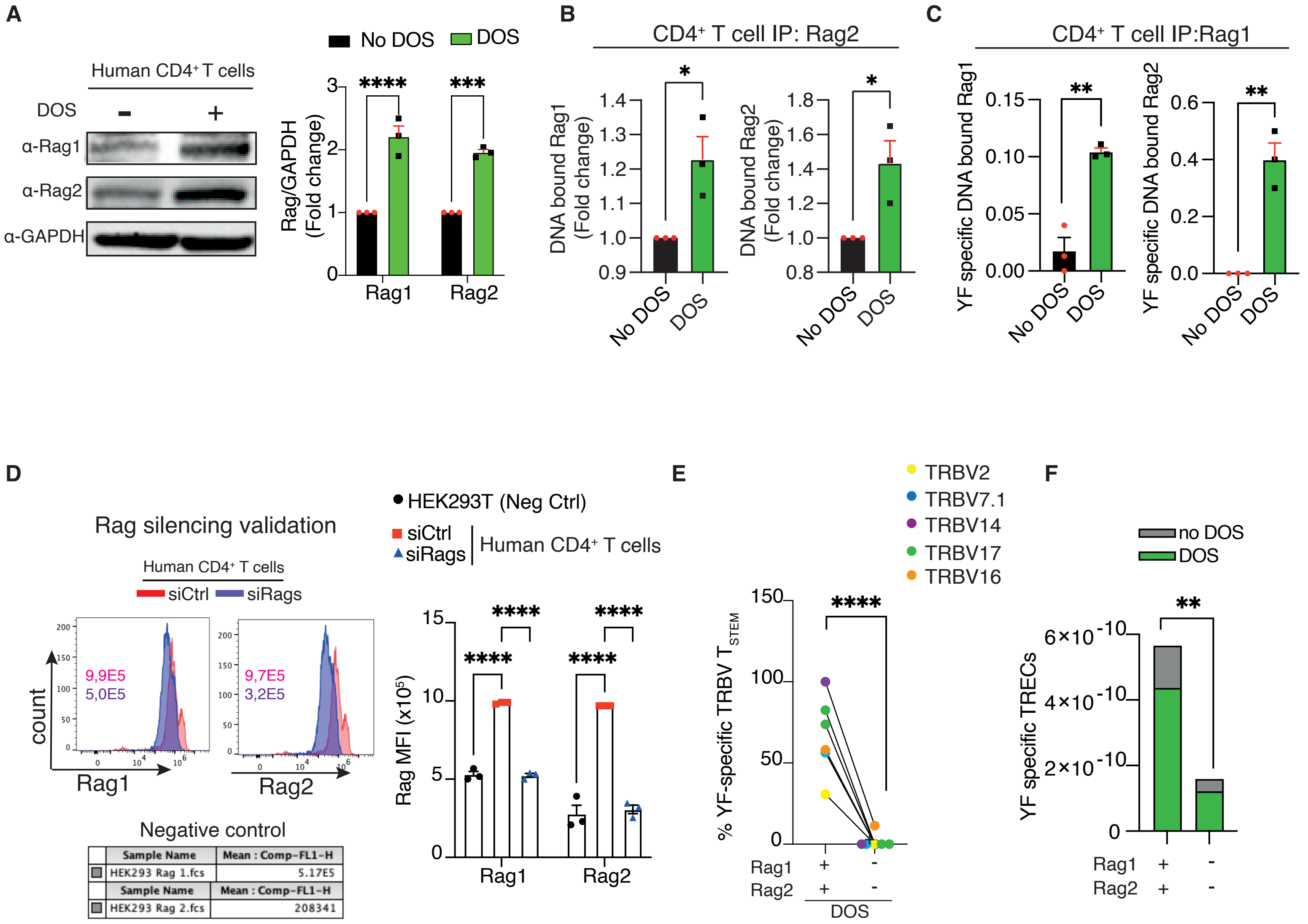

### S12

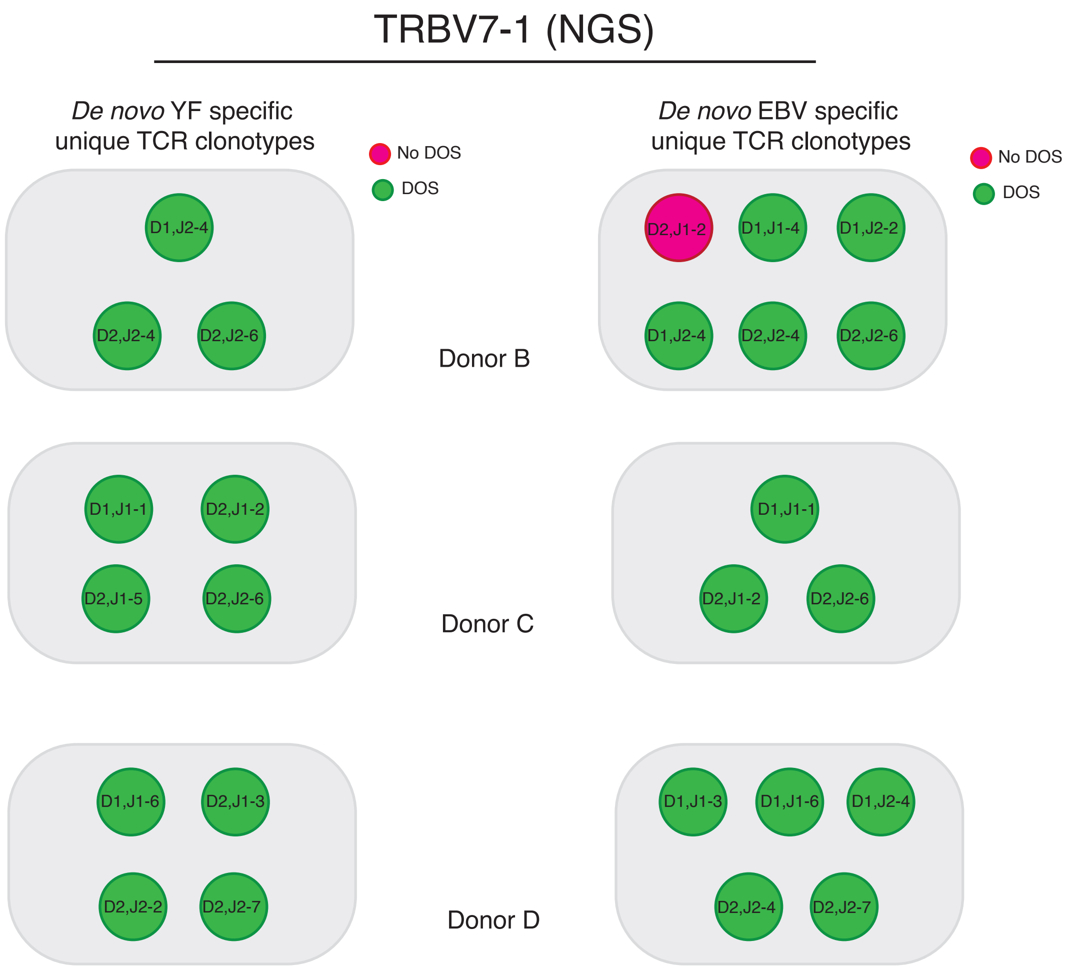

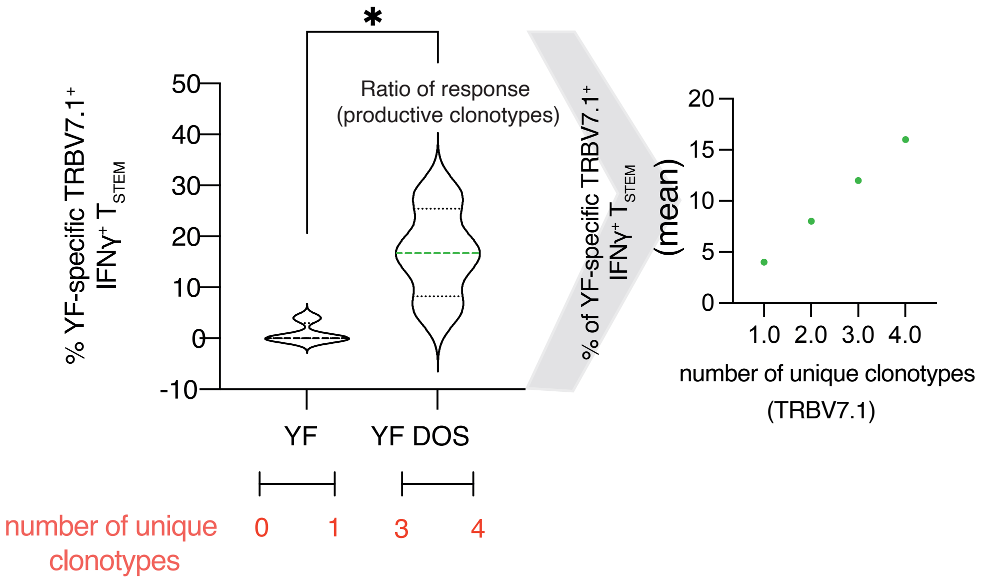
